# Colony context and size-dependent compensation mechanisms give rise to variations in nuclear growth trajectories

**DOI:** 10.1101/2024.06.28.601071

**Authors:** Julie C. Dixon, Christopher L. Frick, Chantelle L. Leveille, Philip Garrison, Peyton A. Lee, Saurabh S. Mogre, Benjamin Morris, Nivedita Nivedita, Ritvik Vasan, Jianxu Chen, Cameron L. Fraser, Clare R. Gamlin, Leigh K. Harris, Melissa C. Hendershott, Graham T. Johnson, Kyle N. Klein, Sandra A. Oluoch, Derek J. Thirstrup, M. Filip Sluzewski, Lyndsay Wilhelm, Ruian Yang, Daniel M. Toloudis, Matheus P. Viana, Julie A. Theriot, Susanne M. Rafelski

## Abstract

To investigate the fundamental question of how cellular variations arise across spatiotemporal scales in a population of identical healthy cells, we focused on nuclear growth in hiPS cell colonies as a model system. We generated a 3D timelapse dataset of thousands of nuclei over multiple days, and developed open-source tools for image and data analysis and an interactive timelapse viewer for exploring quantitative features of nuclear size and shape. We performed a data-driven analysis of nuclear growth variations across timescales. We found that individual nuclear volume growth trajectories arise from short timescale variations attributable to their spatiotemporal context within the colony. We identified a strikingly time-invariant volume compensation relationship between nuclear growth duration and starting volume across the population. Notably, we discovered that inheritance plays a crucial role in determining these two key nuclear growth features while other growth features are determined by their spatiotemporal context and are not inherited.

## Introduction

The question of how variations in the growth and shape of individual cells and their intracellular components arise from their immediate context and then influence population level behaviors is of profound importance to cell biology. We wished to explore this question in a tractable, data-driven manner that permits characterization and direct integration of measurements of growth across multiple scales in both time and space. We focused on the nucleus as a fundamental eukaryotic cellular structure and a compelling model system for exploring these types of cellular variations across timescales. By the time a normal cell, with no karyotypic abnormalities, divides, its nucleus necessarily contains twice as much DNA than when the cell cycle began, marking the nucleus as a key cellular structure exhibiting an extremely stereotyped behavior. So then, if all nuclei duplicate their DNA, do all nuclei also grow identically or is there variation in how individual nuclei grow, even within a tightly controlled population of identical cells in culture? And further, how would any observed variations in nuclear growth arise across timescales and spatial scales?

We chose cultured hiPS cells as an ideal tissue culture model for studying normal nuclear growth in a well-controlled experimental environment to address these questions. hiPS cells are rapidly cycling human cells that represent an early embryonic cell state, are naturally immortal and karyotypically normal.^1^ These cells grow in tightly packed, epithelial monolayer colonies of putatively identical cells on 2D substrates and have been previously used in studies quantifying normal cell to cell variability within populations.^2^ Nuclei in hiPS cells are also a very tractable model from the perspective of performing highly resolved quantitative analysis of the 3D shape of cellular structures through time due to the nucleus being a single structure, having a simple shape, and in hiPS cell colonies making up a large proportion of the cell volume.^2^

Multiple analytical approaches have been previously developed to study the causes and consequences of variability in cellular growth. The size control mechanisms by which cells regulate their volume growth are traditionally characterized as “adder,” “sizer,” and “timer” mechanisms based on whether cells grow to achieve a target added volume, final size or growth duration.^3^ Adder-like mechanisms have been observed in bacteria as well as adherent cell culture systems, mediated by individual cells tuning their growth rate, growth duration, or both based on their starting size to achieve a target added volume.^4,5^ In mice, cells (or nuclei representing cells) studied through intravital imaging were observed to control their size through modulating the duration of G1 to achieve sizer-like size control at the G1/S transition.^6,7^ Further, analysis of isolated cell volume and cell mass trajectories have demonstrated a variety of quantifiable cell growth behaviors over time including linear, bi-linear and exponential growth, often with little difference between model fits.^6,8,9^ These approaches largely rely on individual cell volume trajectories from the start to the end of a cell cycle and thus may potentially also extend to an analysis of variations in analogous individual nuclear growth trajectories.

In a tissue context, populations of cells in growing epithelia have been shown to reduce their size in response to mechanical constraints introduced by crowding within the tissue.^10–12^ As cell cultures approach confluency and begin to exhibit volume reducing divisions, the protein YAP becomes cytoplasmically localized - a behavior known to be correlated with increases in mass density.^13^ In another study the local cell density experienced by a cell’s progenitor was shown to be a good predictor of whether a newborn cell will exit to quiescence,^14^ revealing a potential role of cross-generational lineage in contributing to cellular growth control mechanisms. These studies all together demonstrate the impact the immediate cellular environment and can have on cell growth even across generations and, thus, the importance of developing an approach for analyzing variations in nuclear growth in hiPS cell colonies that explicitly accounts for the potentially varying spatial context surrounding the nuclei over multiple timescales.

We took a data-driven, image-based approach to analyze the variations in the growth of individual nuclei in growing hiPS cell colonies across multiple temporal and spatial scales. For this systematic analysis we produced a high-resolution 3D timelapse dataset capturing the growth dynamics of thousands of nuclei over multiple days. We then quantitatively characterized the distinct ways in which nuclear volume-dependent growth duration, coordinated growth in local cellular neighborhoods, more global colony-wide dynamics and cross-generational lineage each contribute to population-level nuclear growth dynamics.

Alongside the dataset, we necessarily developed novel image analysis and validation approaches, flexible, automated workflows for processing these timelapse images, and reproducible data analysis workflows. We also built an interactive viewer for visualizing and analyzing the resulting quantitative trajectories in a way conducive to data exploration and updated another viewer to facilitate exploration of this 3D timelapse image data. We provide these data, tools and workflows as a resource for future discovery and hypothesis generation.

## Results

### A high-throughput automated workflow to capture highly resolved quantitative features of nuclear shape dynamics in growing hiPS cell colonies

We performed long-term timelapse imaging using the mEGFP-tagged laminB1 cell line from the Allen Cell Collection of endogenously tagged hiPS cell lines (https://www.allencell.org/cell-catalog.html). This cell line retains proper expression level and localization of lamin B1, normal colony morphology, normal growth rate, and the ability to differentiate.^15^ Tagging this structural component of the nuclear lamina allowed us to image and subsequently extract the 3D shape of the nucleus in growing hiPS cell colonies throughout interphase. We performed timelapse imaging, collecting 3D bright-field and fluorescence images of growing colonies at 20x/0.8 NA magnification every five minutes over the course of two days (570 timepoints; Fig. 1A). This magnification, frequency and duration of imaging enabled us to capture nuclear growth dynamics of hundreds of cells within their larger colony context over multiple generations, while spanning a time period of extensive colony growth. We analyzed three baseline colonies with varying initial sizes, to capture colony-size dependent aspects of nuclear growth and shape; we will refer to these three baseline colonies as the “Small,” “Medium” and “Large” colonies, based on their relative starting areas of approximately 31,500, 63,000, and 110,800 µm^2^, respectively (Fig. 1B).

To maintain cell health during the acquisition of these frequent and long duration 3D fluorescence timelapse movies, cells were imaged at 20x/0.8 NA at very low laser powers. Accurate 3D segmentation of nuclear shape from these images was not possible with standard segmentation methods due to the poor axial resolution and low signal to noise ratio (SNR) that result from this imaging. To overcome this limitation, we developed and quantitatively validated a Vision Transformer-based deep-learning segmentation workflow. This model was trained on carefully aligned matched pairs of 3D fluorescence images of mEGFP-tagged lamin B1 acquired at 20x/0.8 NA and ground truth segmentations for those cells derived from imaging the same cells at 100x/1.25NA (Fig. 1C left, D, and E, Supplemental Fig. S1A and B, and Methods). We began with self-supervised pretraining of a Vision Transformer (ViT)^16^ encoder on live 20x nuclear timelapse images followed by supervised training of a convolutional decoder on 20x/100x matched image pairs to generate high-resolution instance segmentations of the nucleus via lamin B1 (Supplemental Fig. S2).

Evaluation of model performance on holdout data (i.e., data not used to train the model) together with a quantitative and application-appropriate validation^17^ approach showed it was able to predict highly accurate segmentations of the same volume and shape as the ground truth (Fig. 1E, Supplemental Fig. S1C-E). Applying this trained segmentation model to the timelapse data resulted in 646,034 segmented nuclei from 570 frames across all three baseline colonies. We call the dataset of all bright-field, fluorescence and segmentation images of these three baseline colonies the “hiPSC baseline colonies FOV-nuclei timelapse dataset,” as part of the “WTC-11 hiPSC FOV-nuclei timelapse dataset V1” which encompasses all timelapse datasets analyzed in this study. Next, to track the shape dynamics of individual nuclei throughout timelapse imaging, an automated workflow was designed to link segmented nuclei from frame to frame resulting in a total of 4,741 single nuclear trajectories (any nucleus that has at least five timepoints; Fig. 1C right). Because the nuclear lamina disassembles upon entry into mitosis, automated single nuclear trajectories were restricted to the interphase portion of the cell cycle and nuclei were not tracked through mitosis. We call this dataset of all nuclear trajectories from the three baseline colonies the “baseline colonies analysis dataset” (see Methods, Fig. 1).

**Figure 1.**
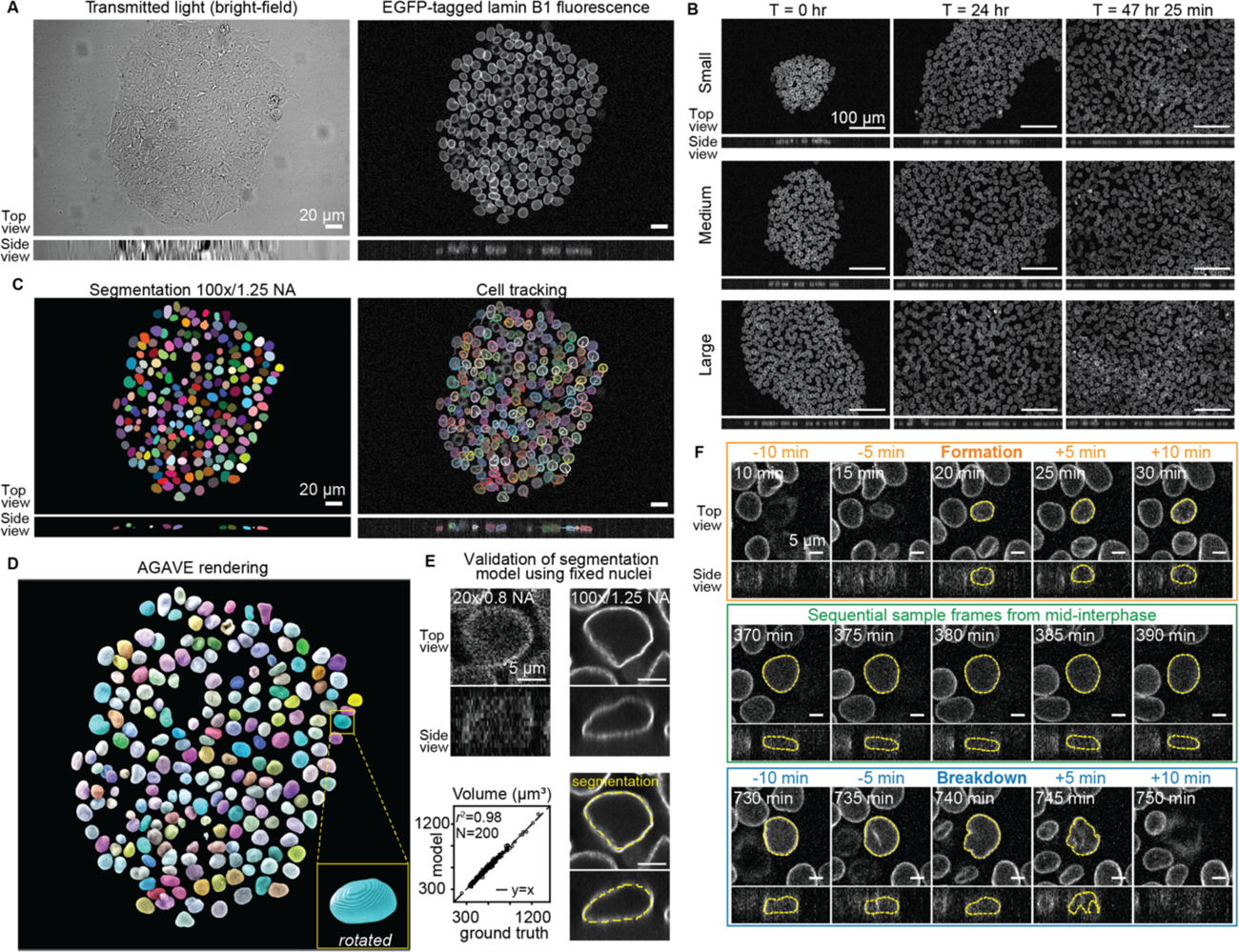
A high-throughput automated workflow to capture highly resolved quantitative features of nuclear shape dynamics in growing hiPS cell colonies. See Methods and Supplemental Fig. S1 and S2 for more details. A. Representative images were generated from the four hour time point of a 20x/0.8 NA, 3D timelapse movie of the Medium hiPS cell colony shown in B. The associated timelapse is provided in Movie S1. Left: Top and side views (middle slice) of the transmitted light bright-field z-stack. Right: Top and side views (maximum intensity projection and middle slice, respectively) of the lamin B1-mEGFP fluorescence z-stack. B. Top and side views (maximum intensity projection and middle slice, respectively) of 20x/0.8 NA, lamin B1-mEGFP images from timelapse imaging of three colonies with different starting sizes, referred to as Small, Medium and Large. Representative images shown for the 0, 24 and 47 hour and 25 minute timepoints. The associated bright-field and fluorescence timelapses for each colony are provided in Movies S1, S2 and S3 and are available for interactive viewing at http://volumeviewer.allencell.org. C. Left: Top and side view (maximum projection and middle slice, respectively) of nuclear segmentations of the images in A. The associated colored segmentation timelapses for each colony are provided in Movies S4, S5 and S6 and are available for interactive viewing together with the bright-field and fluorescence timelapses at volumeviewer.allencell.org. Right: Top and side view (maximum intensity projection and middle slice, respectively) of the mEGFP-tagged lamin B1fluorescence overlaid with the segmentation outline. The tracked centroid location of the 5 timepoints prior are shown for each segmented nucleus as a thin line. Colors indicate different instance segmentations of nuclei for easier viewing. D. AGAVE 3D visual rendering of the nuclear segmentations to highlight their 3D shape. Inset shows an enlarged and rotated view of the nucleus in the yellow box for visualization. E. Top, Left: Top and side view (middle slices) of the lamin B1-mEGFP fluorescence of fixed single nucleus crop imaged at 20x/0.8 NA (Methods). Top, Right: Top and side view (middle slices) of the same fixed single nucleus crop imaged at 100x/1.25 NA. Bottom, Right: Top and side views (middle slices) of the fixed single nucleus crop overlaid with the predicted 3D segmentation (yellow dashed outline). Bottom, Left: Volumes measured from the model-predicted segmentations from 20x/0.8 NA fixed nuclei compared to ground truth segmentations of the same fixed nuclei at 100x/1.25 NA. F. Automated prediction of lamin shell formation and breakdown timepoints for identifying full-interphase nuclear trajectories based on lamin B1-mEGFP images. Top and side views (maximum intensity projection and middle slice, respectively) of 20x/0.8 NA, lamin B1 mEGFP images of a single nucleus at the predicted formation (center, top) and breakdown (center, bottom) timepoints, with two preceding (left) and following (right) timepoints. Also shown is the middle of interphase (middle). Nuclear segmentation outlines for fully formed lamin shells are in yellow.

### Aspects of nuclear shape vary on three timescales, each driven by a distinct source of variation

To help explore dynamic nuclear growth features calculated from the tracked segmented nuclei, we developed the Timelapse Feature Explorer, a web-based tool that allows users to view timelapse segmentations with interactive and customizable colorization mapped according to relative intensities of quantitative features. In this viewer, nuclear segmentations from each individual colony timelapse are colorized by a chosen feature from the baseline colonies analysis dataset, with each nucleus in each frame being colored by its feature value in that frame. The resulting timelapses demonstrate how growth and shape features vary for all nuclei across the colony and over time, allowing for a qualitative assessment of spatiotemoral variation of these features.

We used the Timelapse Feature Explorer to visualize movies of each colony with nuclei colorized by three stereotypical aspects of nuclear shape: height, volume and aspect ratio (Fig. 2, Methods). We observed a gradual evolution of nuclear height over the course of the two-day timelapse, indicating that nuclear height varies slowly, on the timescale of colony growth (for the colony shown in Fig. 2A a gradual decrease in nuclear height). Nuclear volume increases on the timescale of hours (Fig. 2B), consistent with the population’s average cell cycle of 15.6 ± 1.9 hours (mean ± standard deviation), with each nucleus steadily increasing in volume throughout interphase. Finally, we considered the variation of aspect ratio within the colony plane as an indicator of how more complex aspects of shape vary over time. We found that this aspect ratio depends on the local neighborhood within the colony: the aspect ratio increases more rapidly, over a timescale of minutes when nuclei are “squished” by their neighbors, with increasing aspect ratios as they pass through narrow regions between neighboring nuclei (Fig. 2C). Overall, we found that how individual nuclear shapes change over time depends on which aspect of nuclear shape we consider, exemplified by the three different timescales of shape variation for the three different aspects of shape we explored, which result from three distinct sources of variation.

**Figure 2.**
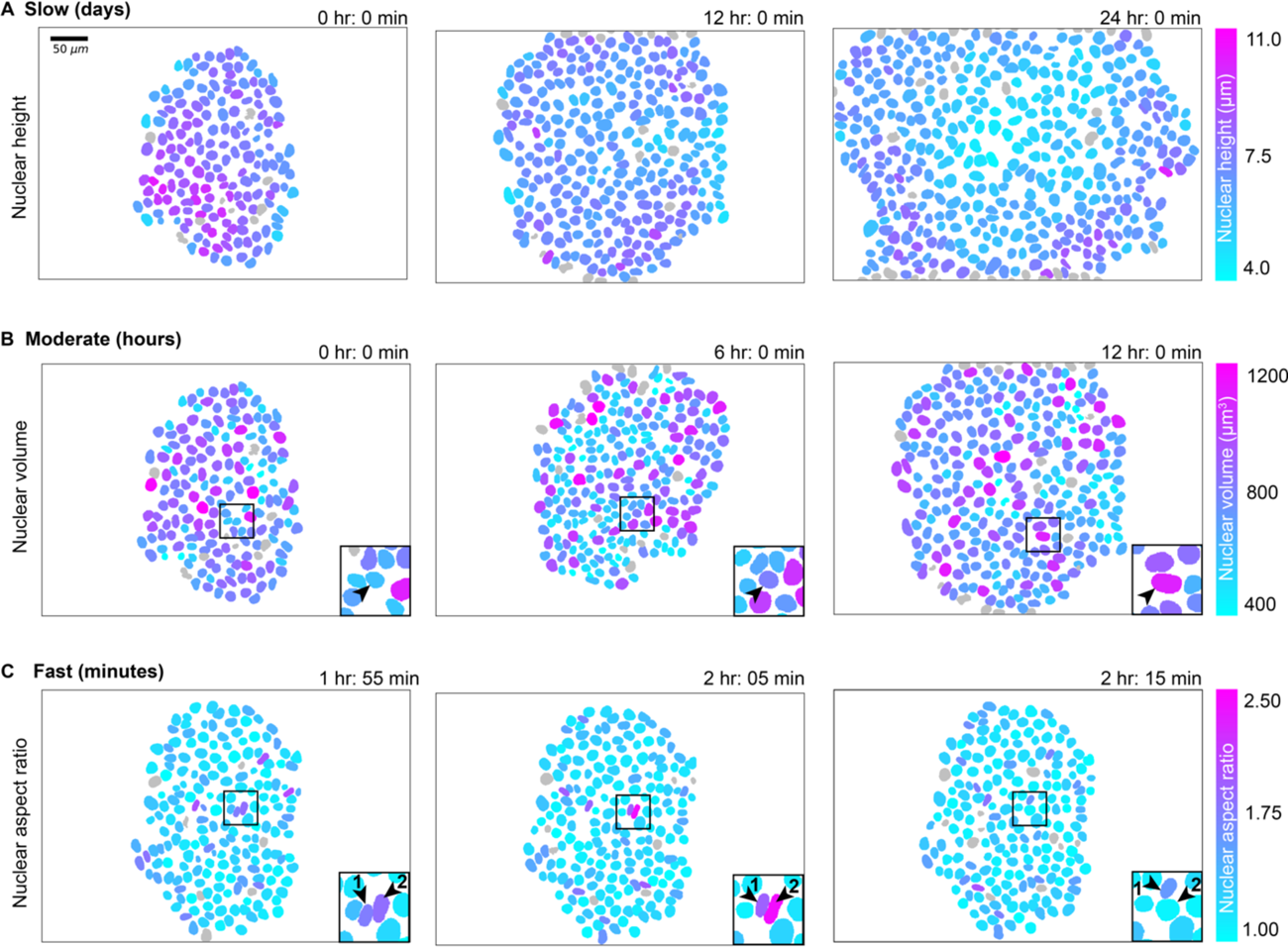
Aspects of nuclear shape vary on three timescales, each driven by a distinct source of variation. Maximum projections of 3D nuclear segmentations color-mapped to features of nuclear shape, created with the Timelapse Feature Explorer (Methods). The associated timelapses for these snapshots were also exported from the Timelapse Feature Explorer and are provided as Movies S7, S8 and S9. Due to the nature of viewing 3D segmentations as 2D maximum projections, the apparent nuclear size and shape in these images can be misleading; the color-mapping of these images is the best way to interpret their quantitative size and shape features. The colormap limits are chosen to highlight the variation of each feature across the population. **A.** Maximum projections of 3D nuclear segmentations colored by height at times t=0 hours, 12 hours and 24 hours. **B.** Maximum projections of 3D nuclear segmentations colored by volume at times t=0 hours, 6 hours, and 12 hours. Inset highlights an individual nucleus as its volume increases over its cell cycle. **C.** Maximum projections of 3D nuclear segmentations colored by aspect ratio (the length of the longest axis divided by the orthogonal in-plane axis) at times t=1 hour 55 minutes, 2 hours 5 minutes, and 2 hours 15 minutes. Inset highlights two neighboring nuclei that change aspect ratio on the timescale of minutes as they “squeeze” past each other. The same colormap is applied for each row of three images and its range is indicated on the right.

### Nuclei exhibit changing spatiotemporal patterns in height as colonies grow

To further explore the gradual changes in nuclear height as the colonies grew, we compared the height-colorized timelapses of all three baseline colonies and discovered a distinctive spatiotemporal pattern in the behavior of nuclear height in the baseline colonies analysis dataset, consistent across all three colonies (Fig. 3A). At the start of the Small colony timelapse, there is a radial height gradient, with the tallest nuclei (magenta) in the center of the colony, and shorter nuclei (cyan) in the surrounding, outer edges of the colony. As time passes and the colony grows, this gradient gradually diminishes and eventually reverses; by the end of the timelapse, the center of the colony is filled with shorter nuclei while the outer nuclei are taller. The Medium colony also begins with taller nuclei in the center, though to a lesser extent than the starting frames of the Small colony; we again see this gradient diminish and eventually reverse, with an even larger “lake” of shorter nuclei filling the colony center surrounded by taller nuclei at the colony edges. Finally, the Large colony starts with the lake of nuclei and we eventually see this spatial pattern reverse yet again as later in the timelapse, the height-colorized center nuclei become more magenta, indicating that they are again growing taller.

When we plotted the average height of nuclei across the field of view for all nuclei in the baseline colonies analysis dataset over the course of each of the colonies’ respective timelapses we found that initially the height of each colony decreased over time. However, for the Medium and Large colony, we observed that the average height of the nuclei increased towards the end of the timelapse. This consistent pattern of nuclear height dynamics and its overlapping quantitative behavior across all three colonies provided a way to temporally align the three colonies relative to one another. We determined the time lag that minimizes the mean squared difference between the average height trajectories of each colony, resulting in one single universal timeline of colony development (“aligned colony time”) across all three colonies (Fig. 3B, Methods).

The aligned colony time collapses spatial data from throughout the colony into a single measurement at each timepoint for easy comparison across colonies and across time. However, we were also interested in quantifying the evolution of the radial spatial pattern observed in different heights across colony development which required a quantitative metric for the radial location of a nucleus within the colony. Since the colonies were not uniform in their radial size, we developed a “normalized distance from colony center” metric, such that nuclei at the center and outer edge of the colony (or field of view once colonies grow larger) have a normalized colony depth of 0 and 1, respectively (Fig. 3C top left, Methods). We then determined the slope of a linear fit of the colony height relative to the colony depth at each timepoint for each colony; a positive slope indicates that height increases as the position moves radially outwards (i.e. shorter nuclei are in the center) while a negative slope would indicate that tallest nuclei are found in the center at that time (Fig. 3C top right, Supplemental Fig. S3A). Plotting these slopes across aligned colony time for all nuclei in the baseline colonies analysis dataset allows us to quantify the spatial pattern of nuclear height as the colony develops (Fig. 3C bottom). This plot showed a negative slope earlier in aligned colony time, which increased and became positive for larger colony sizes. This trend then reversed for the timepoints with largest colony sizes. This approach quantitatively captures the spatial pattern observed in Fig. 3A: colonies start with taller nuclei in the center, then this pattern reverses as the colony grows as a lake of shorter nuclei that fill the colony interior, and finally this pattern reverses again with taller nuclei pushing up in the colony center.

**Figure 3.**
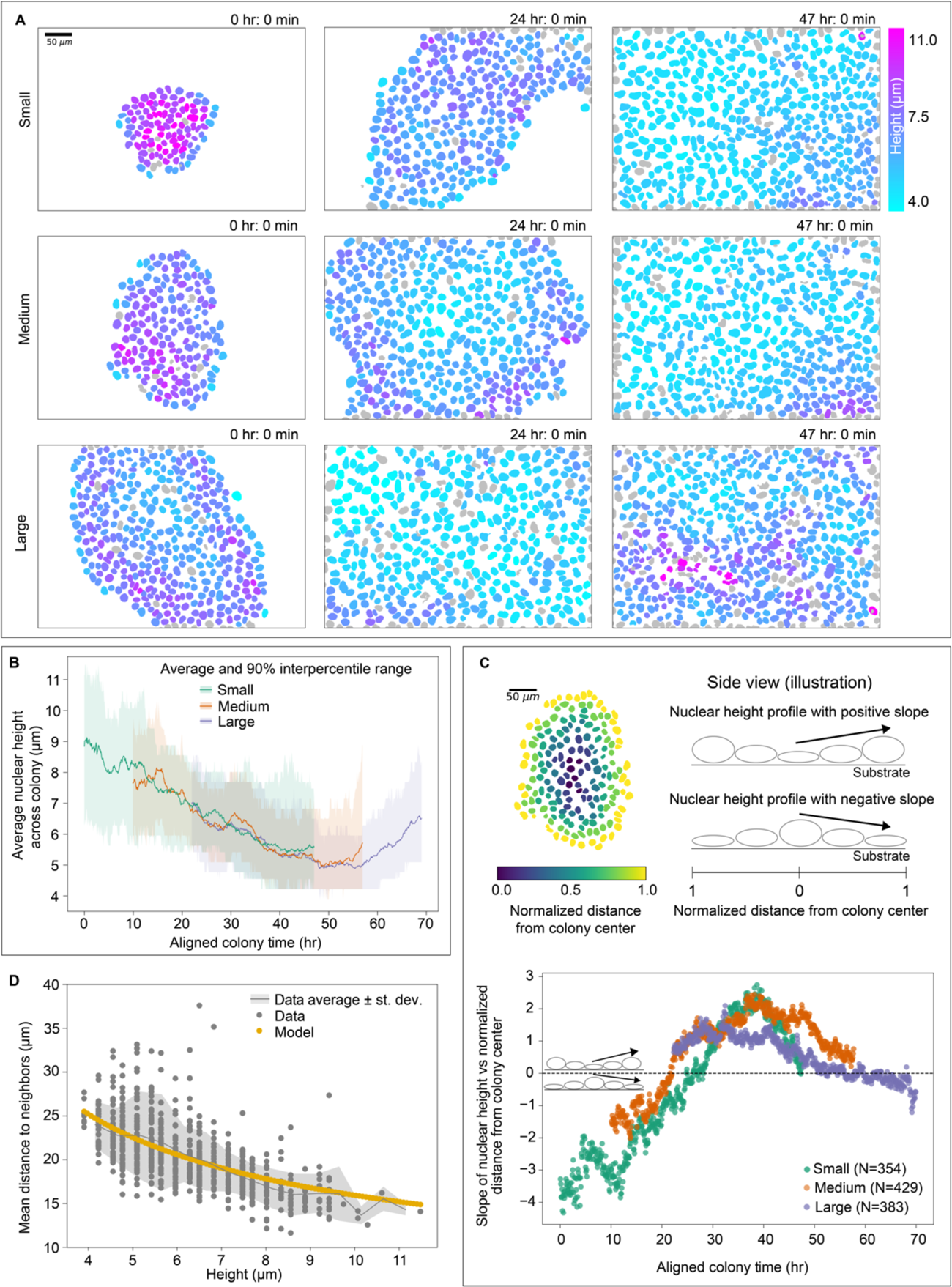
Nuclei exhibit changing spatiotemporal patterns in height as colonies grow. **A.** Maximum projections of 3D nuclear segmentations colored by height for the Small, Medium, and Large colonies at 0 hours, 24 hours, and 47 hours, created with the Timelapse Feature Explorer (Methods). The associated timelapses for these snapshots were also exported from the Timelapse Feature Explorer and are provided as Movies S7, S10 and S11. The same color map is applied to all nine images. **B.** The average nuclear height trajectory for the Small, Medium and Large colonies. The height trajectories for each colony have been aligned along the time axis to maximize the overlap between them (Methods). This represents the “aligned colony time” used in other analyses. The shaded region is the 5th to the 95th percentile (“90% interpercentile range”) for nuclear height. **C. Top, Left:** Maximum projection of the segmented image of the Medium colony at t=0, colored by the normalized distance from the colony center (Methods). **Top, Right:** A cartoon illustration of the side view of the colony with two example height profiles where the slope is measured radially from the colony center. The slope is positive when the nuclei at the center of the colony are shorter than the nuclei closer to the edge or image field of view (“FOV,” Methods). **Bottom:** The slope of the best linear fit line for the height vs. normalized distance from the colony center for all colony timepoints, for all three colonies (see Supplemental Fig. S3A for examples). The inset shows the cartoon of the side view of the colony to illustrate different regimes above and below the zero-slope reference line in the height profile. **D.** The relationship between the mean distance between neighbor centroids and the nuclear height is shown for nuclei with volumes of 630 µm^3^ (gray; Methods). Modeling the cells as hexagonal cylinders (yellow) allows them to pack closely in alignment with known data.

We hypothesized that nuclear height dynamics could be driven by crowding in the growing colony environment. To explore the link between nuclear height and local density while controlling for nuclear volume, we considered a model of nuclei as right hexagonal cylinders with fixed volumes which showed an excellent fit to the data without any free parameters (Fig. 3D, Methods). These results demonstrates that nuclei of consistent volume grow taller as the local density increases and they run out of room in the colony plane. We then quantified the average density in the colony, taken as the average over the entire colony of local density experienced by each nucleus at each time point (see Methods), and found, as expected, that this local density follows the same temporal pattern as the colony-averaged nuclear height (Fig. 3B, Supplemental Fig. S3B).

### Nuclear volume trajectories consistently contain two phases and, on average, double in volume over the growth phase that spans most of interphase

As expected, colony timelapses of nuclear volume demonstrated that volume increased steadily over the course of interphase. In order to ask how consistent the behavior of this volume change was across individual nuclei, we created a dataset containing only nuclear trajectories tracked through the entirety of interphase, referred to as the “full-interphase analysis dataset,” which is a subset of the baseline colonies analysis dataset (see Methods). During interphase, lamin B1 localizes to the nucleoplasmic side of the inner nuclear membrane^18^, forming a complete shell defining the shape of the nucleus. The lamin shell grows throughout interphase until it eventually breaks down before the cell enters mitosis, then reforms in each daughter cell at the start of the next interphase. We developed an automated image analysis workflow which, for each nucleus, identifies the start and end of interphase, defined respectively as the first frame in which the complete lamin shell has formed (“formation”) and the last frame before the first sign that this shell breaks down (“breakdown”) (Fig. 1F, Methods). After manual curation to identify outlier trajectories and errors in nuclear segmentation, tracking or timepoint classification from automated workflows (see Methods), a total of 1,166 nuclei were successfully tracked for the entirety of interphase, formation to breakdown, across the three colonies (354, 429, and 383 nuclei for the Small, Medium, and Large colony, respectively).

Taking advantage of this large and robust full-interphase analysis dataset, we examined individual nuclear volume trajectories across both real time (Fig. 4A) and a “Normalized interphase time”, defined as the time spanning from lamin shell formation to breakdown normalized by the interphase duration, such that formation and breakdown occur at *t=0* and *t=1,* respectively, for all full-interphase trajectories (Fig. 4B). These nuclear trajectories have a strikingly consistent biphasic volume growth behavior. Trajectories start with a short phase of rapid nuclear expansion which is then followed by a transition into a longer, slower nuclear growth phase that spans most of interphase (Fig. 4C). We developed an automated workflow to determine when this transition between these two phases occurs, finding an average transition time of 38 ± 10 minutes into interphase. We stained live nuclei in an hiPS cell line expressing mEGFP-tagged PCNA with a DNA dye (SPY650) and collected six-hour timelapse data. Based on a combination of the DNA and mEGFP-tagged PCNA patterns (Methods) we estimated the timing of this transition relative to the onset of S-phase. We found that S-phase begins much later than the transition point, shown by an example of a nucleus entering S-phase approximately 2.5 hours after mitosis (Supplemental Fig. S4, Methods), indicating that the transition from rapid expansion to growth is not due to the G1/S phase transition. Furthermore, perturbing cells with importazole, an inhibitor of nuclear import, caused significant disruption to nuclear growth during the rapid expansion phase, with nuclei born after the addition of importazole reaching volumes 86% that of the control unperturbed nuclei (Supplemental Fig. S5). Together, these results demonstrate that the early phase of rapid volume increase is likely an import-mediated expansion as the nucleus swells immediately following mitosis, consistent with recent observations.^19^ We therefore focused the subsequent analysis of nuclear volume trajectories on the post-expansion, nuclear growth phase that occurs for the majority of interphase. We also found that a small subpopulation of 42 nuclei (3.5%) exhibit growth dynamics that are distinct from the rest of the population (Methods and Supplemental Fig. S6). These nuclei exhibit dramatically longer growth phases and, out of the daughters they produce whose fates we could determine, two thirds subsequently undergo apoptosis, which may be related to observations that long mitosis durations in mother cells result in daughters who die in the next generation.^20^ For these reasons we excluded this sub-population from the full-interphase analysis dataset.

**Figure 4.**
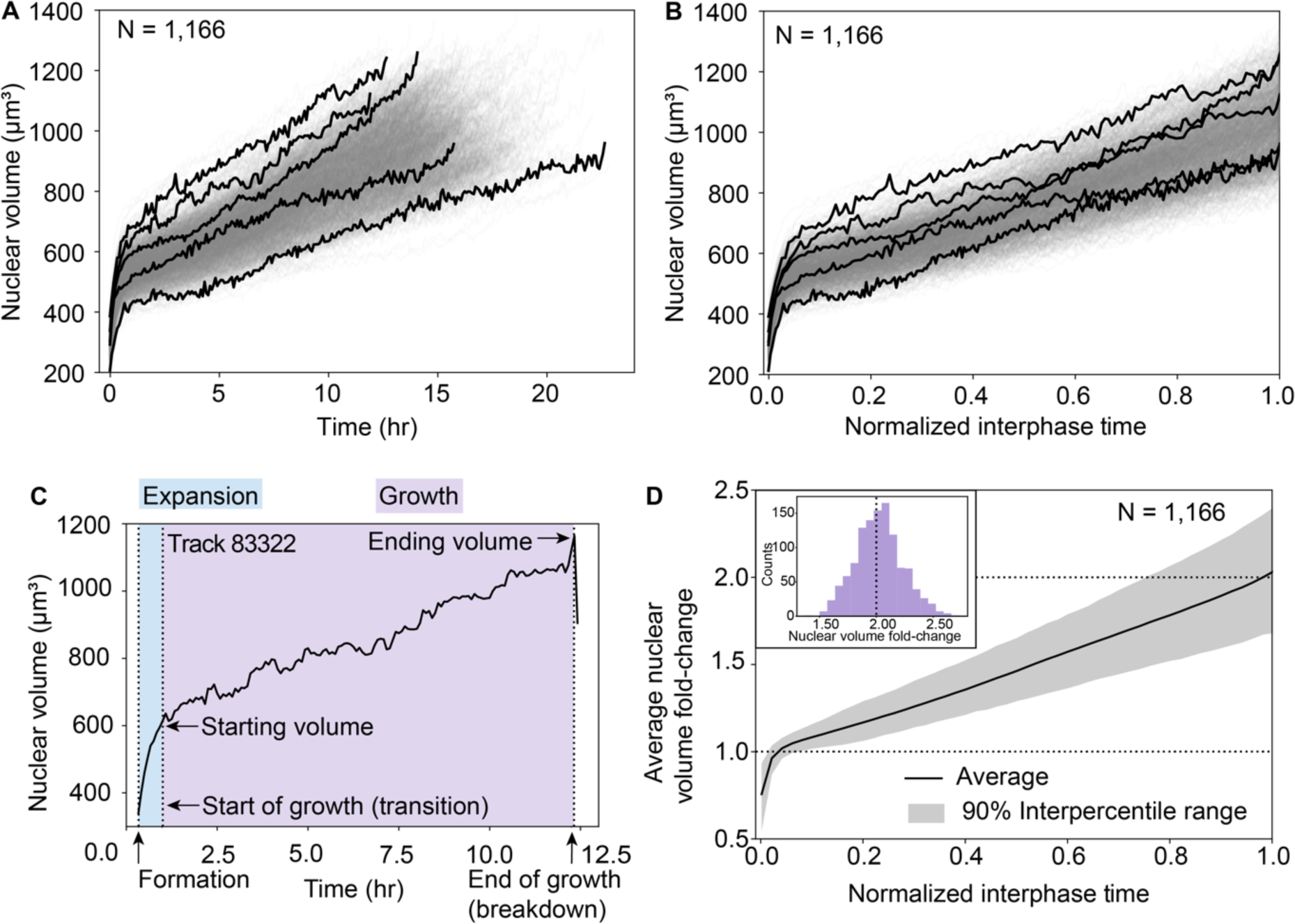
Nuclear volume trajectories consistently contain two phases and, on average, double in volume over the growth phase that spans most of interphase. **A** and **B.** Volume trajectories for all nuclei in the full-interphase analysis dataset, shown over real time synchronized by the time of lamin shell formation (A) and normalized interphase time (B). All 1,166 trajectories are shown in transparent gray and five trajectories are shown in black to highlight some of the variation in individual nuclear growth. **C.** A sample nuclear volume trajectory from formation to breakdown. The calculated transition point (Methods) defines the transition between a short, rapid nuclear expansion (“expansion”, shaded blue) and a longer, slower growth period (“growth”, shaded purple, begins at the “start of growth”). The “starting volume” and “ending volume” are the nuclear volumes at the start of growth (transition) and end of growth (breakdown). **D.** The average nuclear volume fold-change and 90% interpercentile range of full-interphase nuclear trajectories, synchronized by the time of shell formation and rescaled to normalized interphase time. Nuclear volumes at the start of growth were normalized to 1. Dashed reference lines at nuclear volume fold-change of 1 (start of growth) and 2 (doubling from the start of growth). Inset shows the distribution of nuclear volume fold-change for the population.

We observed a largely linear average nuclear growth trajectory and a near-perfect doubling in the average nuclear volume fold-change from the start to the end of nuclear growth for all nuclei in the full-interphase analysis dataset (peak of the distribution at 2.05-fold, Fig. 4D). Even though the average volume trajectories differ slightly between colonies (Supplemental Fig. S7A), this result of population-level average linear growth and volume-doubling holds true not only for the pooled population of all colonies, but also for each colony individually (Supplemental Fig. S7B). Based on these results, we tested a possible model in which DNA replication plays a major role in driving nuclear volume trajectory behaviors. When we treated cells with aphidicolin, an inhibitor of DNA replication, we found only a slight difference in average volume trajectories between perturbed and control nuclei, suggesting that while replication has some impact on growth, it does not dictate nuclear volume growth dynamics (Supplemental Fig. S7C and Methods). This is consistent with a recently published model suggesting that osmotic pressures at the nuclear membrane due to soluble proteins in nucleus and cytoplasm - rather than nuclear DNA content directly - are the dominant forces in determining nuclear size.^21^ Furthermore, we observe variations in individual nuclear volume doubling, with some nuclei exhibiting volume fold-changes that are significantly larger or smaller than two-fold, which also contradicts a model where genome content doubling during replication directly leads to nuclear volume doubling. We, therefore, next asked whether some nuclear size control mechanism exists to maintain population-level nuclear doubling despite the variation in individual nuclear growth trajectories that we observed.

### Individual nuclear volume trajectories vary widely in shape, ranging significantly from sub-to super-linear growth

To understand how individual nuclei control their growth and achieve population-level doubling, we first asked how individual nuclear volumes increase over time. Cellular volume growth has largely been shown to be exponential or near-exponential in the context of individual cells^5^; proteins are produced by ribosomes and ribosomes themselves are built out of protein, resulting in exponential production of cellular material. Is nuclear growth likewise exponential, and if so, for the same reasons? While linear and exponential growth models are appealing for their facile links to biological mechanisms, and the average nuclear growth trajectories seem quite linear (Fig. 4D), when we look at the individual nuclear volume trajectories in aligned colony time, we see that their trajectory shapes vary widely, ranging from what appears to be sub-to super-linear growth (Fig. 5A). We, therefore, began with a flexible quantitative approach to capture the overall shape of the nuclear volume trajectory. We fitted the growth phase of each of the trajectories in the full-interphase analysis dataset to a power-law in time 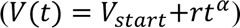, with the fitted time scaling factor (α) quantifying how super-linear (α>1) or sub-linear (α<1) a given trajectory is (examples of sub-and super-linear trajectories are shown in Fig. 5B top). We found that α values across the population take on a wide, continuous distribution, peaked at α∼1.15 (Fig. 5B bottom), consistent for all three colonies (Supplemental Fig. S8A). The breadth of the distribution is consistent with the observation that many nuclear volume trajectories display significantly sub- and super-linear behaviors with time (Fig. 5A). This makes it clear that a simple linear or exponential model of nuclear volume growth would be entirely inappropriate for this data.

**Figure 5.**
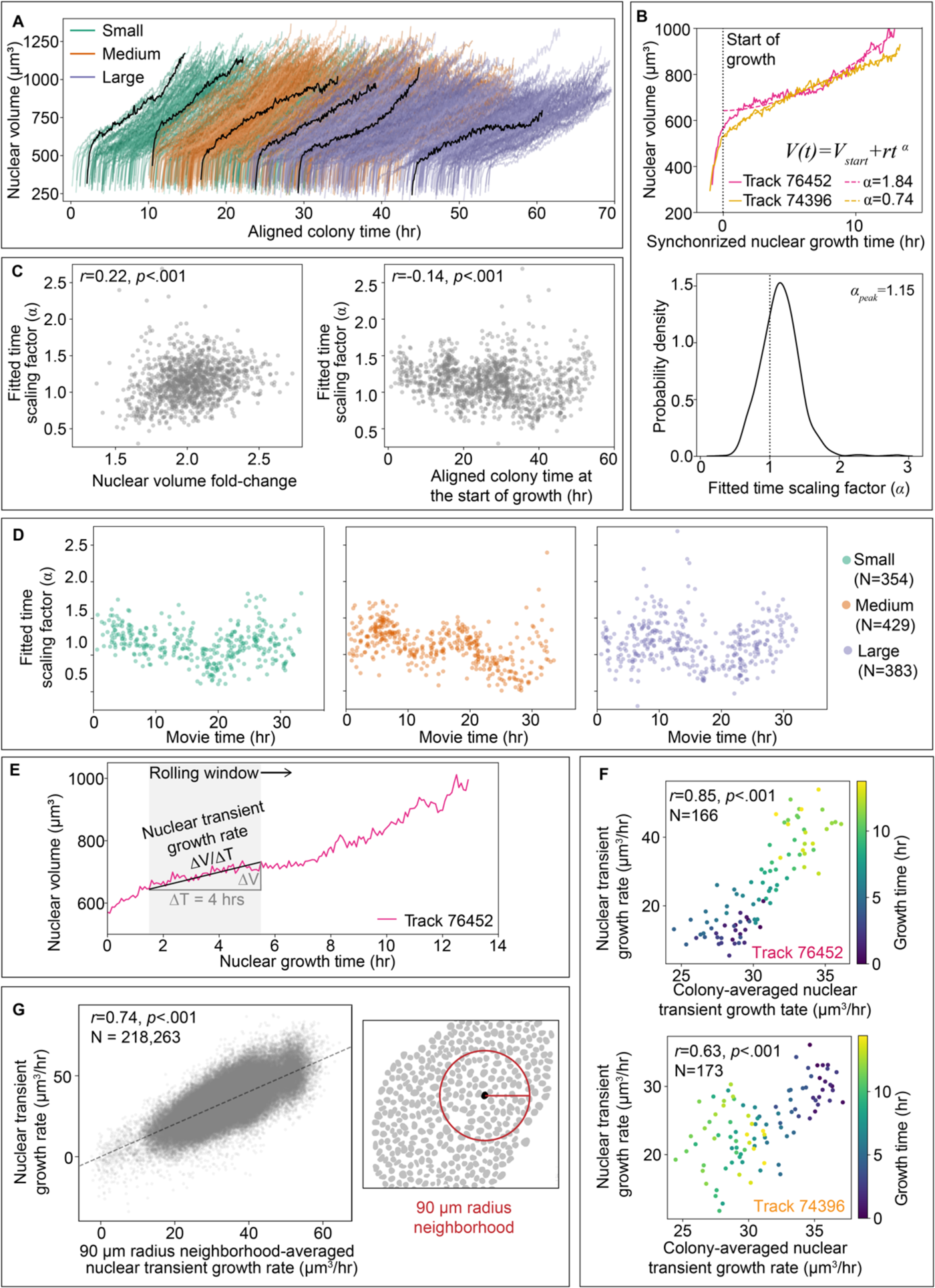
Transient nuclear growth rates are dominated by when and where nuclei are in the hiPS cell colony. **A.** Nuclear volume trajectories for all nuclei in the full-interphase analysis dataset from the Small, Medium, and Large colonies over aligned colony time (N=1,166). Six example full-interphase volume trajectories are shown in black to highlight the variation in trajectory shapes observed during nuclear growth. **B. Top:** Each full-interphase nuclear volume growth trajectory (from the start to the end of growth) was fit to a power-law in time 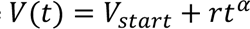 with fitted time scaling factor (α) (Methods). Example super-linear (α>1) and sub-linear (α<1) nuclear volume trajectories are shown, synchronized by their start of growth. **Bottom:** The probability density of the fitted time scaling factor (α) has a modal value of 1.15 for the full-interphase analysis dataset (N=1,166). **C.** Relationships between the fitted time scaling factor (α) and the nuclear volume fold-change **(Left)**, and aligned colony time at the start of growth **(Right)** for each full-interphase trajectory in the full-interphase analysis dataset (N=1,166) with Pearson’s correlation coefficient (*r*) and p-values (*p*) indicated. **D.** The fitted time scaling factor (α) along actual time within a single colony movie (movie time) for each of the Small, Medium, and Large colonies. **E.** Nuclear transient growth rates were calculated over a rolling window of 4 hours at each timepoint along each nuclear volume trajectory resulting in a series of sequential growth rates. **F.** The nuclear transient growth rates for the example trajectories shown in B (**Top:** super-linear, magenta**; Bottom:** sub-linear, yellow) compared to the colony-averaged nuclear transient growth rate in the same time window, colored by time throughout nuclear growth. **G. Left:** The nuclear transient growth rate for all individual nuclear growth time windows compared to the neighborhood-averaged nuclear transient growth rate in a 90 µm local neighborhood at the same time. The Pearson’s correlation coefficient (*r*), p-values (*p*) are sample size (N) are indicated. **Right:** Example colony segmentation image with 90 µm radius shown for scale.

### Transient nuclear growth rates are dominated by when and where nuclei are in the hiPS cell colony

We hypothesized that the observed variation in nuclear volume trajectory shapes might give rise to the individual variations around population-level nuclear volume doubling, with nuclei having sub- and super-linear volume trajectories under- and over-shooting volume doubling, respectively. Surprisingly, we found only a weak correlation of 0.22 between the fitted time scaling factor (α) and the nuclear volume fold-change (Fig. 5C left). Next, we investigated whether, instead, volume trajectory shapes were related to the changes in the overall colonies themselves that we observed over aligned colony time (Fig. 3B) by considering the relationship of fitted time scaling factors to aligned colony time (Fig. 5C right).

Interestingly, we did not see a consistent long-timescale shift in α values over aligned colony time, as we did for height. Instead, the cloud of α values contained what seemed like a vague appearance of some collective rising and falling fluctuating pattern, though this pattern is not particularly clear or strictly periodic. To investigate this further, we considered the variation of α values over individual colony movies; splitting out the nuclei from each movie in this way resulted in less variation in the fitted time scaling factor values at any given time and displayed somewhat more distinctive fluctuations (Fig. 5D). The timing of these fluctuations could capture the full range of α values within a 7-10 hour period. The shortness of this timescale relative to the average growth duration of 14.9 ± 1.9 hours suggests that nuclear growth trajectory shape may change on a timescale faster than an individual’s cell cycle and that a single power-law fit to the volume over all of growth may be insufficient to capture the subtleties of shorter timescale growth dynamics. This may also explain why these nuclear volume power-law fits have errors greater than can be explained by segmentation error (Supplemental Fig. S8B and Methods). We wondered whether an individual nucleus’ growth rate kinetics should, therefore, not be considered based on the entire growth trajectory, and instead be considered at a shorter timescale that could capture the varying behavior of nuclear growth over time. We calculated the transient growth rates of nuclei in the full-interphase analysis dataset on shorter timescales (four hours) over a rolling window in time resulting in a series of sequential growth rates from each individual trajectory (Fig. 5E). We reasoned that perhaps the value of α was reflective of the difference in transient growth rates towards the beginning and end of an individual nuclear volume trajectory. Indeed, we found a strong correlation between values of α and the difference between the average early and late transient growth rates, as measured by the average of transient growth rates calculated in the first 30% and last 25% of time windows, respectively (Supplemental Fig. S8C). This suggests that the transient volume growth rate metric still captures the main aspects of the change in trajectory shapes (as represented by α), but now provides the flexibility to explore growth rates at timescales shorter than the entire trajectory.

Following up on the collective fluctuations in α values observed across individual colonies, we next considered the relationship of nuclear transient growth rates to the average nuclear transient growth rate across the colony (“colony-averaged nuclear transient growth rate”). First, we consider the two example trajectories (Fig. 5B top) to gain insight into how their transient growth rates relate to the colony-averaged nuclear transient growth rate over time (Fig. 5F). While it is unsurprising that the example nuclei with super- and sub-linear growths individually have steadily increasing (top) and decreasing (bottom) growth rates over interphase time, it is remarkable the extent to which this reflects broader variations experienced across the entire colony on shorter, four-hour time scales. Furthermore, these examples highlight the fact that nuclear transient growth rates do not share a common cell-cycle dependence; trajectories may start with faster nuclear transient growth rates then slow, or vice versa.

Instead, nuclear transient growth rates appear to be related to the transient growth behaviors of their colony-level environment. We next pooled not the growth rates derived from entire nuclear growth trajectories, as is typically done, but instead the nuclear transient growth rates for all individual nuclei in the population. We then compared every nuclear transient growth rate measurement (multiple overlapping instances for each nuclear growth trajectory) with the colony-averaged nuclear transient growth rate from that same window in time and, consistent with the two example trajectories, found a strong correlation (Supplemental Fig. S8D). Furthermore, we find an even higher correlation when we instead consider a “neighborhood-averaged nuclear transient growth rate”, for which we pool only the nuclei within a local neighborhood of 90 μm radius to compare with the nuclear transient growth rate of each individual nucleus (Fig. 5G). The growth rate of nuclei is therefore spatially as well as temporally coordinated, with these shorter timescale spatial variations in nuclear growth rates imposed by the local neighborhood environment. This fundamental result highlights the important role of environmental context in driving individual growth dynamics within a colony context.

### Nuclei display an adder-like growth mechanism to maintain population-level volume doubling

Our analysis of nuclear trajectory shapes highlighted the deep impact of local context on individual variation in nuclear growth trajectories. How, then, do all of the variations in growth rates observed at the shorter, more transient timescales of hours and at the smaller spatial scales of cellular neighborhoods scale up and transform, via the behavior of entire individual nuclear growth trajectories, into the average nuclear volume doubling that is observed in the full-interphase analysis dataset? One hypothesis is that perhaps individual nuclei grow toward a doubling in their volumes (“volume fold-change” of two) between the start (“starting volume”) and the end (“ending volume”) of nuclear growth. In this case the nuclear starting volume should have no effect on its volume fold-change; regardless of whether a nucleus starts out smaller or larger at the onset of growth, it would still, on average, double in volume. Instead, we found that the nuclear volume fold-change was moderately and negatively correlated with nuclear starting volume (*r*=-0.52) demonstrating a systematic effect in which smaller nuclei undergo a larger fold-change and larger nuclei a smaller fold-change (Fig. 6A). Further, we found that nuclear volume fold-change was also not invariant to any other basic features extracted from nuclear growth trajectories (Supplemental Fig. S9A), suggesting that the nuclear volume fold-change itself is not directly targeted or regulated and instead seems to be an important outcome of other mechanisms of size control.

The negative correlation between nuclear volume fold-change and starting volume is an observation consistent with an adder model of cell growth.^22^ In comparison to the sizer model of growth, the hallmarks of the adder model, here considered for nuclei instead of cells, is that, on average, the amount of volume that nuclei in a population grow (“added volume”) is invariant to nuclear starting volumes and that the distribution of added volume has a coefficient of variation (CV) that is ∼1.73-fold larger than the coefficient of variation of the distribution of nuclear ending volumes 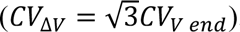.

**Figure 6.**
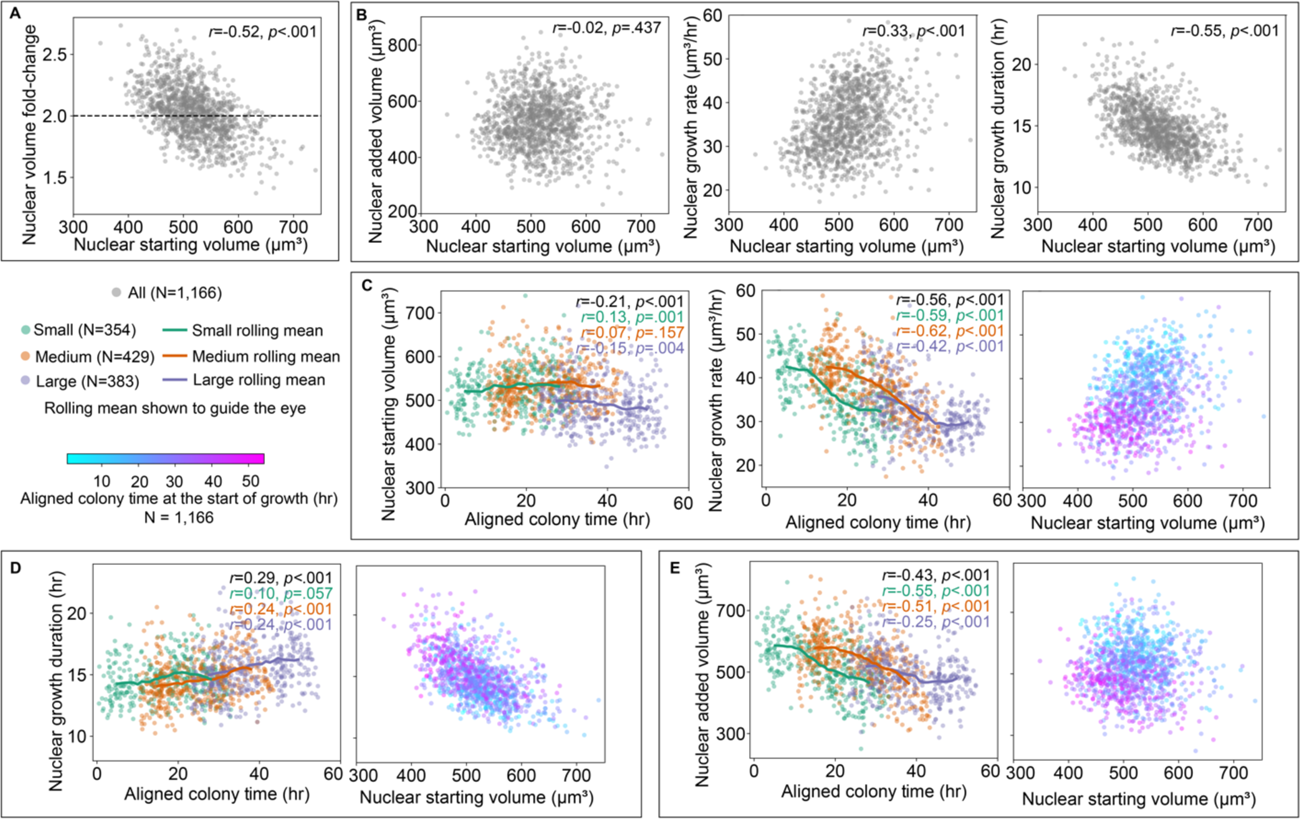
Nuclear added volume is an outcome of colony context-dependent nuclear growth rates and colony context-independent nuclear growth durations. All correlations reported in this figure are Pearson’s correlation coefficients (*r*) with p-values (*p*) for all nuclei in the full-interphase analysis dataset, reported in black for all nuclei pooled across all three colonies and in green, orange, or purple for each colony separately. The N for each population is indicated in the legend along with the color map for aligned colony time. **A.** Relationship between the nuclear volume fold-change and the nuclear starting volume for all full-interphase nuclear trajectories. Dashed reference line at nuclear volume fold-change of 2 (doubling from the start of growth). **B.** The relationships of nuclear added volume, growth rate, and growth duration with nuclear starting volume for all full-interphase nuclear trajectories. **C. Left, Middle:** Nuclear starting volume and nuclear growth rate over aligned colony time at the start of growth for the Small, Medium, and Large colonies, with rolling mean shown to guide the eye. **Right:** Relationship between nuclear growth rate and nuclear starting volume for all full-interphase nuclear trajectories (shown in B) colored by aligned colony time at the start of growth. **D-E, Left:** Nuclear growth duration and nuclear added volume over aligned colony time at the start of growth, with rolling mean shown for the Small, Medium, and Large colonies. **D-E, Right:** Relationships between the nuclear growth duration and nuclear added volume for all full-interphase nuclear trajectories (shown in A, B) colored by aligned colony time at the start of growth.

We found that the added volume was uncorrelated with the nuclear starting volume (*r*=-0.02, Fig. 6B left). We also found that the measured coefficient of variation for added volume (0.177) was 1.72-fold greater than the measured coefficient of variation for ending volume (0.103). While the wide range of nuclear volume trajectory shapes we observed is not consistent with the use of strict modeling approaches that presume exponential growth toward identifying mechanisms of size control, these observations together are consistent with an adder-like regime of size control.^22^ Both the dependence of nuclear volume fold-change on nuclear starting volume and the overall average population-wide volume doubling during growth are outcomes of this type of size control mechanism.

### Nuclear added volume is an outcome of colony context-dependent nuclear growth rates and colony context-independent nuclear growth durations

Where, then, do the observed individual nuclear added volumes come from? For any nucleus the added volume is the outcome of the combination of how fast and for how long a nucleus grows. Here, we define the nuclear “growth rate” for the entire growth phase of an individual nucleus as the change in nuclear volume (same as the “added volume above”) divided by the “growth duration” from the start to end of growth (as opposed to the “transient growth rate” previously described). While analysis of the nuclear volume fold-change and added volume tells us about the relationship between nuclear volumes at the two endpoints of the growth phase, it does not tell us anything about how growth rate and growth duration between these endpoints together give rise to the resulting added volume population distribution. To address this, we explored the relationships of both growth rate and growth duration with nuclear starting volume (Fig. 6B middle and right). When data is pooled from all three colonies, we found that both these features were correlated with the starting volume - a weak positive correlation in the case of the growth rate (*r*=0.33) and a moderate negative correlation in the case of growth duration (*r*=-0.55). It may be tempting to suggest that the positive correlation between growth rate and nuclear starting volume, together with the negative correlation between growth rate and growth duration, indicates a potential size control mechanism whereby smaller nuclei grow slower and thus would need to grow longer to be within the range of added volume seen in the population. In this scenario, perhaps the variations we found in nuclear growth trajectories that arise at shorter timescales from the local neighborhood context serve to weaken the strength of the correlation between growth rate and starting volume. However, as we explored this result further, we discovered it to be misleading for two reasons. First, we found that average nuclear starting volumes remain largely constant over the majority of aligned colony time but take an appreciable dip late in aligned colony time seen only for the Large colony (Fig. 6C left). Meanwhile, we found that the growth rate drops steadily over aligned colony time for all three colonies (Fig. 6C middle). When the scatter plot of growth rate vs. starting volume is colored by aligned colony time (Fig. 6C right), we see from the downward progression of cyan to magenta that this correlation for the pooled data only exists because of these two time-dependent behaviors, primarily due to the Large colony (Supplemental Fig. S9B). This analysis demonstrates that in addition to the observations of local colony neighborhood effects contributing to growth rates at a shorter, transient timescale (Fig. 5), there exists a second, longer timescale colony context-dependent effect on nuclear growth, in which nuclei grow more slowly over all of interphase later in aligned colony time. This is consistent with observations that increasing colony size tends to lead to slowing of cell growth due to increased tissue confinement.^10^ We also considered whether the decrease in nuclear growth rate (and therefore also nuclear ending volume and added volume) could result from a depletion of nutrients in the colony. We performed a set of control experiments comparing the nuclear growth rates in colonies with various adjustments to when they receive fresh media (see Methods), and found that nutrient depletion does not explain the observed decrease in growth rate (Supplemental Fig. S10).

We next analyzed the effect of aligned colony time on the moderate negative correlation between nuclear growth duration and starting volume. We found a slight increase in growth durations over aligned colony time (Fig. 6D left). However, in this case there is little, if any difference in the relationship between nuclear growth duration and starting volume across colonies and thus, an almost complete overlap between data from earlier and later timepoints (Fig. 6D right). The result directly interpreted from the pooled dataset in (Fig. 6B right) therefore holds true: the growth duration has a negative correlation with the starting volume throughout the population, and that duration remains the same across aligned colony time. This suggests that cells consistently compensate for their lower nuclear starting volumes by growing for the same longer duration regardless of aligned colony time.

If the growth rate (which is independent of starting volume) drops but the growth duration relationship (which is dependent on starting volume) stays the same over aligned colony time, we would predict that the resultant added volume would also depend on aligned colony time. Indeed, we found that the average added volume drops with aligned colony time (Fig. 6E) while remaining independent of nuclear starting volume, suggesting that nuclear growth in hiPS cell colonies seems to occur via a “colony time-dependent adder-like” size control mechanism. Consistent with this observation, we found a similar dependence on aligned colony time for the resulting nuclear ending volume (Supplemental Fig. S9C) and for the relationship between nuclear volume fold-change and starting volume (diagonal striation in Supplemental Fig. S9D). This would also explain the decreased average volume over normalized interphase time for the large colony, which has more data at these later aligned colony timepoints (Supplemental Fig. S7A). Therefore, the nuclear added volume seems to arise as an outcome of the interplay between the local and more global context-dependent growth rates and the inherent control of growth duration based on nuclear starting volume.

### Variations in nuclear starting volumes arise from the variations in nuclear ending volumes of their progenitors in the previous generation

We have shown that variation in nuclear growth rates within individual nuclear growth trajectories can be attributed to local neighborhood-level spatial and temporal variations on a short timescale, while a more global colony development-level behavior causes slower, subtler shifts in nuclear growth features such as initial, final and added volume. Despite this, two major nuclear growth features - growth duration and volume fold-change - demonstrate consistent dependence on the starting nuclear volume for individuals across the population. Furthermore, the distribution of starting volumes in the full-interphase analysis dataset is quite broad (517.9 ± 54.9 μm^3^; Fig. 7A). How then do the starting volume and its variations arise? One hypothesis is that the source of variations in the starting nuclear volumes of cells arises from the variations in ending nuclear volumes of their direct progenitors. In support of this idea, we found that the population distributions of starting and ending volumes have equal coefficients of variation (Fig. 7A). We can link the starting and ending volumes and quantify their correlation within individual trajectories (Supplemental Fig. S9C). However, from these histograms alone we cannot determine how much of this variation in starting volumes arises from variations in the ending volumes of the previous generation (the progenitors) and to what extent this variation is inherited along individual lineages. To link nuclear trajectories across two to three generations we manually curated the Small and Medium sized colonies and created 315 independent lineage trees resulting in a “lineage-annotated analysis dataset,” which is a subset of the “full-interphase analysis dataset.” (Fig. 7B, Methods). We considered related cells with a single generation gap (“mother” and “daughter” nuclei, N=359 “mother-daughter” pairs) or with no generation gap (“sister” nuclei born from the division of a common “mother” nucleus, N=257) for which we had full-interphase trajectories. Data from these colonies were pooled into a single population for analysis, as no significant differences were observed between colonies when treated separately. We then addressed whether the nuclear volumes of cells at the start of the growth phase following rapid expansion are related to the ending nuclear volumes of their mothers. We found that the sum of nuclear starting volumes of sister pairs (“combined nuclear starting volume”) was strongly correlated with the mother’s ending volume (*r*=0.83). Further, this combined nuclear starting volume summed to, on average, 95.9 ± 4.2% of the mother’s ending nuclear volume (Fig. 7C and Supplemental Fig. S11A) falling systematically just slightly below the unity line. Thus, combined starting volumes of daughter nuclei are strongly related to and recapitulate the population-wide variation of the ending nuclear volumes of their mothers.

Another possible contribution to the variation in overall nuclear starting volumes is how much of the mother’s nuclear starting volume each sister in the pair inherits; if inheritance were perfectly equal, then the variation seen in individual nuclear starting volumes would be a direct outcome of the variations in ending volumes of the mother nucleus. On the other hand, if sister nuclei showed very unequal starting volumes, this might contribute significantly to the population-level variations in nuclear starting volumes.

We therefore compared sister nuclear starting volumes to each other and found a moderate positive correlation (*r*= 0.46, Fig. 7D), while the nuclear starting volumes of unrelated nuclei born at the same location and time within the colony were much less correlated (*r* = 0.15, Supplemental Fig. S12A, Methods). We next explored how much asymmetry in sister starting volumes contributes to the overall starting volume variation by comparing sister nuclei that are most symmetric with those that are most asymmetric (Supplemental Fig. S11B left, Methods). If sister asymmetry was a strong source of the overall starting volume variation, we would expect the group of most asymmetric sisters to show a significantly broader distribution in starting volumes than the most symmetric group of sisters. Further, we would expect the asymmetric sister pairs to be located at opposite ends of the distribution centered around the average starting volume of the population. Instead, we found that while the group of asymmetric sisters does show a broader distribution centered around a similar average starting volume, it is only moderately broader (coefficient of variation ± 90% confidence intervals of 0.11 ± 0.01 and 0.08 ± 0.01 for the asymmetric and symmetric groups, respectively; Fig. 7E). Furthermore, the asymmetric sister pairs do not themselves represent the opposite tails of the distribution and are only 140 μm^3^ apart (Fig. 7E); the starting volume values from the most asymmetric sister pair is indicated with arrows on the distribution plot as an example. Consistent with the observation that the variation in nuclear starting volumes variation is largely independent of sister asymmetry, neither the relationship between nuclear starting volume and growth duration nor nuclear volume fold-change differ significantly between the most symmetric and most asymmetric sister pairs (Supplemental Fig. S11B center and right, Supplemental Fig. S11C). Together, these results indicate that while asymmetric division has some impact on the variation in starting volumes and on the dependence of growth durations and fold-changes on those starting volumes, it is not the primary or sole source of these growth behaviors. Together these analyses suggest that nuclear starting volumes are inherited from mothers to their daughters, that this inheritance is largely independent from when and where within the colony the daughters are located, and that the overall variation in nuclear starting volumes observed in the population primarily arises from the variation in nuclear ending volumes of the progenitor.

**Figure 7.**
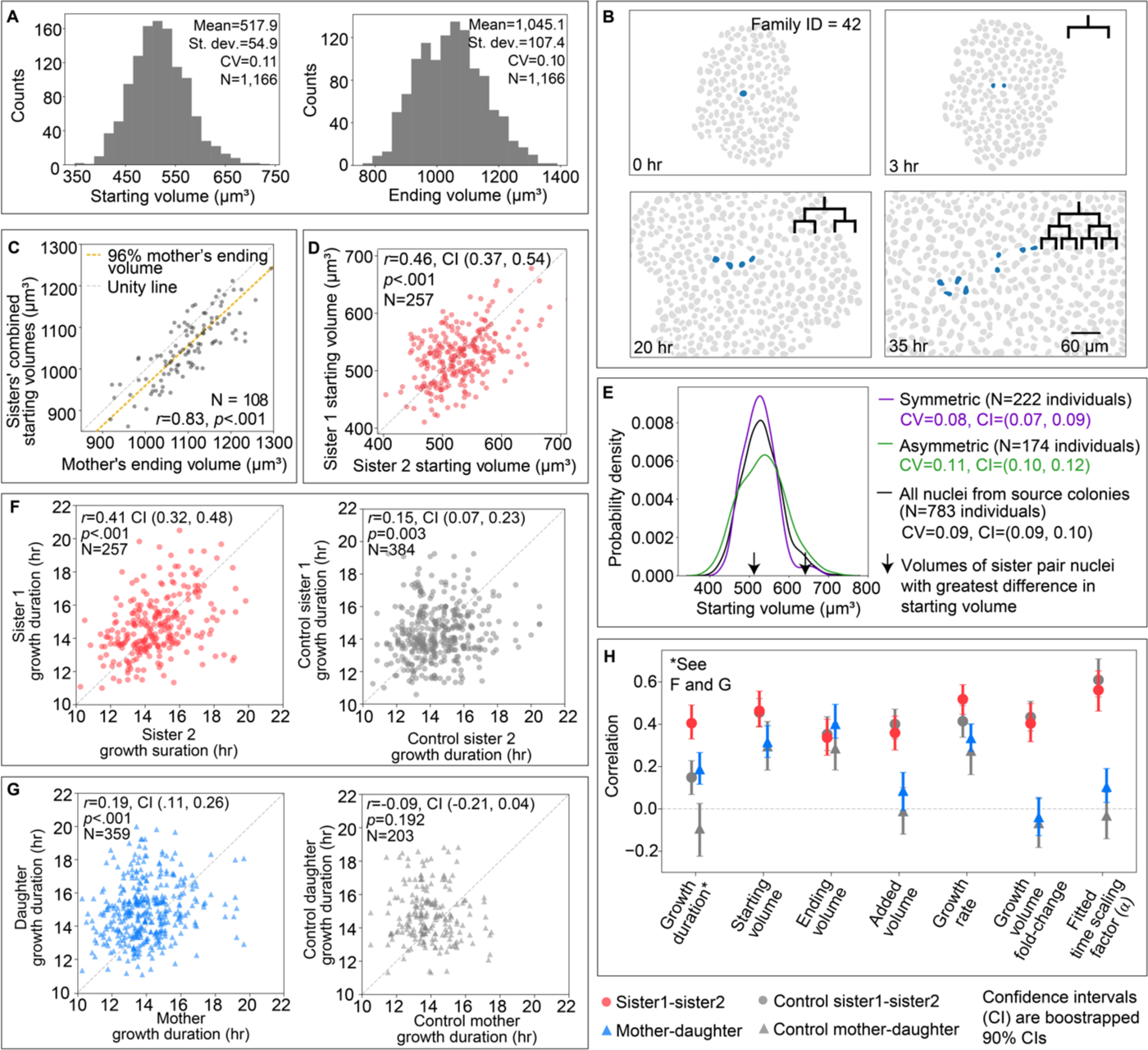
Nuclear starting volume and growth duration are inherited from one generation to the next, while other nuclear growth features depend on their colony context. All correlations reported in this figure are Pearson’s correlation coefficients (*r*) with p-values (*p*). All confidence intervals (reported numerically or shown as error bars) are bootstrapped 90% interpercentile confidence intervals (see Methods for details). **A.** Distribution of nuclear starting volumes and ending volumes with the mean, standard deviation, and coefficient of variation (CV) for all nuclei in the full-interphase analysis dataset. **B.** Maximum projection of nuclear segmentations for the Medium colony colored by an example lineage tree at the start of each new generation (0, 3, 20, and 35 hour timepoints). In this representative lineage tree, two generations are captured for their entire interphase (3-20 hrs and 30-35 hrs). **C.** The relationship between the sisters’ combined starting volumes and their mother’s ending volume. Dashed reference lines are shown for the unity line (black) and for 96% of the mother’s ending volume, the average percent of the mother’s nuclear ending volume reached by the sisters’ combined nuclear starting volume (red; Supplemental Fig. S11A). **D.** The relationship between the starting volumes for sister pairs. The unity line is shown for reference. **E.** The probability density of the starting volume for symmetric sisters, asymmetric sisters, and all nuclei from the lineage-annotated analysis dataset with the CV for each (Methods). Arrows denote the starting volumes for the most asymmetric sister pair as a reference for the maximum distance sister pairs would span in the starting volume distribution. **F. Left:** The relationship of growth durations for all sister pairs in the lineage-annotated analysis dataset. **Right:** The relationship of growth durations for mother-daughter pairs. **G. Left:** The relationship between two control sister pairs which are unrelated nuclei born within 10 minutes, a 60 µm radius, and a difference in starting volume less than 80 µm^3^ (Methods, Supplemental Fig. S12B). **Right:** The relationship between control mother-daughter pairs which are unrelated nuclei where the control mother’s breakdown is within 60 minutes and a 60 µm radius of the control daughter. Their size is constrained such that the control daughter’s starting volume is within 60 µm^3^ of half the control mother’s ending volume (Methods). **H.** The growth duration correlation (*r*) between sister pairs, mother-daughter pairs and their respective control pairs from the scatter plots in panel G are plotted in H, with error bars indicating the bootstrapped 90% confidence intervals on these correlations. Correlation values with confidence intervals between sister pairs, mother-daughter pairs and their respective controls are also shown for nuclear starting volume, ending volume, growth rate, volume fold-change, and fitted time scaling factor (α) (See Supplemental Fig. S12C for the equivalent scatter plots as in F and G).

### Nuclear growth duration is inherited from one generation to the next while other nuclear growth features depend on their colony context

To address whether other aspects of nuclear growth in addition to the starting volumes are inherited, we took advantage of the lineage-annotated analysis dataset. An approach to determining whether a particular feature is inherited is to compare the strength of the relationship of this feature either between mother and daughter or between sister pairs with unrelated pairs in the population. Given the earlier results that the transient, local neighborhood context of nuclei significantly contributes to nuclear growth, we chose to compare mother-daughter and sister pairs with other cells in the immediate vicinity (both in space and time; Supplemental Fig. S12B left and middle, Methods) of these cells. Further, since we found that the starting volumes of daughters are largely inherited from the ending volumes of their mothers, and many other growth features are correlated with nuclear starting volume, we chose to further control for the starting volumes as well (Supplemental Fig. S12B right, Methods). So, for example, the correlation of growth duration between sister pairs (*r*=0.41) is significantly greater than the correlation of growth duration between “control sister” nuclei in the local vicinity and with similar starting volumes (*r*=0.15) (Fig. 7F and H). The significance of the difference in these correlations is determined by their non-overlapping bootstrapped 90% confidence intervals (Methods). Similarly, the correlation of growth duration between mother-daughter pairs (*r*=0.19) is significantly greater than that of an “control daughter” and an “control mother” nucleus in its local vicinity, whose ending volume is twice the “daughter’s” starting volume (*r*=-0.09, Fig. 7G). This example demonstrates that growth duration is indeed more similar for related nuclei (both mother-daughter and sister pairs) when compared to proper controls and thus likely an inherited feature of nuclear growth. All other key features of nuclear growth, however, showed no difference between related and controlled unrelated pairs, consistent with the finding that each of these features either were correlated with the local neighborhood of other nuclei or the nuclear starting volume (Fig. 7H, Supplemental Fig. S12C).

## Discussion

Throughout this study, we found the nuclear growth duration to have striking behavior, distinct from other nuclear growth features in its consistency over generations and throughout colony development. The growth duration and nuclear starting volume are the only growth features we measured that were inherited, while for all of the other growth features their correlation between related cells was accounted for by being born at their inherited starting volume and at the same time and place within the colony (Fig. 7). Furthermore, the size-dependent growth duration compensation mechanism is uniquely invariant to aligned colony time (Fig. 6D), leading us to speculate that the tuning of nuclear duration based on inherited nuclear starting volume and the inheritance of growth duration itself may provide the means by which consistent nuclear growth behaviors at the population level arise despite colony context-dependent variations in nuclear growth dynamics.

We found that the nuclear ending volume, added volume and growth rate all decreased slowly over the timescale of colony development (Fig. 6, Supplemental Fig. S9). The time-invariance of the size-dependent nuclear growth duration together with the slowly decreasing growth rate results in what can be considered a time-dependent adder-like growth mechanism, in which the size-invariant distribution of nuclear added volumes decreases slowly over time. Meanwhile, this set of nuclear growth features that decrease over long timescales are found to be most similar among nuclei found not only at the same time in colony development but also in the same local neighborhood (Fig. 7H, Fig. 5G). In fact, while sister cells inherit a similar nuclear starting volume and growth duration, their growth trajectories diverge via these more local, transient effects such that by the end of growth they are just as similar in volume to cells in their local neighborhood as to each other (Fig. 7H). Interestingly, the local and global behaviors of these growth features may both be related to changes in the density of the epithelial colony. On a local level, we speculate that coordination of transient growth rates within local neighborhoods arise as nuclei respond to shorter-timescale changes in the local spatial density of cells due to mitotic events, motility-associated squeezing (as nuclei and cells move past one another; see Fig. 2C) and, perhaps, filling in gaps between cells that have left the colony after apoptotic events.^7^ Meanwhile, slow decreases in these aspects of nuclear growth over time may be linked to changes to the overall spatial cell density as the colony grows and develops with nuclei growing taller in later aligned colony time (Fig. 3). This hypothesis is consistent with observations of YAP, the primary transcription factor of the size-control implicated Hippo pathway,^23,24^ changing its localization to the cytoplasm as colonies reach confluency.^10^ Unexpectedly, we found that as the nuclear growth rates decline steadily over earlier parts of aligned colony time, the average nuclear height and density are also decreasing, suggesting that other mechanisms may be at play. Together these variations and their possible sources highlight the deep impact of the environment context on nuclear growth dynamics.

A common thread of these systematic data-driven analyses is the careful consideration of how to disentangle and interpret sources of variations of nuclear growth dynamics across a population and over multiple timescales. While it may not be entirely surprising that the immediate spatial context in which cells grow and the inheritance across generations contribute to variations in nuclear growth dynamics, this dataset and our analysis approach are able to *directly measure* to what extent distinct aspects of nuclear growth are impacted by which sources of variations. This specific mapping of variations in growth features to their sources offers an impactful resource for the emerging field of multiscale modeling, in which the specification of rules for cell behavior relies deeply on quantitative evidence of the targeted impacts of intrinsic and external factors. Our analyses also demonstrate the challenges in analyzing variations in timelapse data across large populations, exemplified by our analysis of nuclear growth rates and their relationship to nuclear starting volume. Firstly, the interpretation of the growth rate as measured across the entire growth phase of individual nuclei is limited, given the observed short timescale, local variation in transient growth rates. Furthermore, the weak correlation of this growth rate to starting volume (*r*=0.33) was found to be a “red herring” - arising only because of temporal variations of growth dynamics in a single colony, and not reflecting a population-wide size-dependent tuning of growth rate.

While the complexities of variations in nuclear growth and shape over multiple scales of space and time inherent in this dataset present subtleties and challenges for analysis, they also offer a rich resource for future discovery and hypothesis generation. We provide open access to this dataset along with the tools and workflows used to process, visualize and analyze it. While we have focused the study largely on features of the volume growth dynamics of nuclei, additional nuclear size and shape features, including those measured in this dataset and future features that can be derived from the nuclear segmentations in this dataset, remain ripe for exploration. Furthermore, the quantitative and dynamic nature of this data make it a lasting resource for the scientific community to use for producing and testing models of nuclear growth and shape dynamics across timescales.

## Acknowledgments

We thank Ben Gregor for automated stem cell maintenance, Chamari Wijesuriya and Jie Yao for microscopy support, Janani Gopalan for program management, Kevin Mitcham for software infrastructure support and the Allen Institute for Cell Science team for helpful discussions and support. We also thank Hongxiao Wang and Danny Chen for exploring the feasibility of computational “transfer functions”, a predecessor of the transformer-based models in this paper. We thank Rick Horowitz, Gaudenz Danuser, Quincey Justman, Gant Luxton, and Wallace Marshall for helpful scientific discussions. The WTC line that we used to create our gene-edited cell lines was provided by the Bruce R. Conklin Laboratory at the Gladstone Institute and UCS. J.A.T. was supported by the Howard Hughes Medical Institute. This article is subject to HHMI’s Open Access to Publications policy. HHMI lab heads have previously granted a nonexclusive CC BY 4.0 license to the public and a sublicensable license to HHMI in their research articles. Pursuant to those licenses, the author-accepted manuscript of this article can be made freely available under a CC BY 4.0 license immediately upon publication. S.M.R. and C. L.F. were supported for some of this work by the National Human Genome Research Institute of the National Institutes under Award Number UM1HG011593. The content is solely the responsibility of the authors and does not necessarily represent the official views of the National Institutes of Health. We wish to acknowledge the Allen Institute for Cell Science founders, Jody Allen and Paul G. Allen, for their vision, encouragement and support.

## Author contributions

Conceptualization, J.C.D, C.L.Frick, C.L.L, M.P.V., J.A.T., and S.M.R.; Methodology, J.C.D, C.L.Frick, C.L.L., B.M., N.N., P.A.L., P.G., R.V., S.S.M., C.L.Fraser, D.J.T., M.F.S, J.C., K.N.K., L.K.H., D.M.T., M.P.V., J.A.T., and S.M.R.; Software, J.C.D, C.L.Frick, C.L.L., B.M., N.N., P.A.L., P.G., R.V., S.S.M., C.L.Fraser, M.F.S., J.C., L.W., D.M.T., and M.P.V.; Validation, J.C.D., C.L.Frick., C.L.L., B.M., N.N., P.G., R.V., S.S.M., M.F.S., J.C., M.V.P., and S.M.R.; Formal Analysis, J.C.D., C.L.Frick, C.L.L., N.N., R.V., S.S.M., M.V.P., and S.M.R.; Investigation, J.C.D, C.L.Frick, C.L.L., B.M., N.N., R.V., S.S.M., M.F.S., J.C., K.N.K., L.K.H., M.C.H., R.Y., M.V.P., J.A.T., and S.M.R.; Resources, J.C.D, C.L.Frick, C.L.L., B.M., N.N., P.A.L., P.G., R.V., S.S.M., C.L.Fraser, D.M.T., and M.V.P.; Data Curation, J.C.D. C.L.Frick, C.L.L., B.M., N.N., P.G., C.R.G., M.F.S., S.A.O., D.M.T., and M.V.P.; Writing-Original Draft, J.C.D., C.L.Frick, C.L.L, and S.M.R.; Writing – Reviewing and Editing, J.C.D., C.L.Frick, C.L.L, B.M., P.G., R.V., S.S.M., M.F.S., M.V.P., J.A.T., and S.M.R; Visualization, J.C.D., C.L.Frick, C.L.L, N.N., P.A.L., R.V., S.S.M., C.L.Fraser, L.W., D.M.T., and M.V.P.; Supervision, J.C.D., G.T.J., D.M.T., M.V.P., and S.M.R.; Project Administration, J.C.D., C.L.Frick, C.L.L, C.R.G., D.M.T., M.V.P, and S.M.R.

## MATERIALS AND METHODS

### RESOURCE AVAILABILITY

#### Lead contact

Further information and requests for resources and reagents should be directed to and will be fulfilled by the lead contact, Susanne Rafelski.

#### Materials availability

Using the Wild Type WTC-11 hiPS cell line background,^25^ we previously generated the Allen Cell Collection of hiPS cell lines in which each gene-edited cell line harbors a fluorescent protein endogenously tagged to a protein representing a distinct cellular structure of the cell.^15^ The cell line AICS-0013 cl 210 (RRID:CVCL_IR32), which express mEGFP-tagged lamin B1; and AICS-0088 cl 83 (RRID:CVCL_A8RT), which express mEGFP-tagged proliferating cell nuclear antigen (PCNA) are described at https://www.allencell.org and are available through Coriell at https://www.coriell.org/1/AllenCellCollection. For all non-profit institutions, detailed MTAs for each cell line are listed on the Coriell website. Please contact Coriell regarding for-profit use of the cell lines as some commercial restrictions may apply.

#### Data and code availability

We release all timelapse data used in this study in the OME-Zarr format to democratize their access. Users can download all the timelapse, training, and analysis datasets from Quilt programmatically or using the Quilt user interface using the links below.

WTC-11 hiPSC FOV-nuclei timelapse dataset V1: https://open.quiltdata.com/b/allencell/tree/aics/nuc-morph-dataset/hipsc_fov_nuclei_timelapse_dataset/

- hiPSC FOV-nuclei timelapse data for pretraining Vision Transformer containing FOV bright-field and fluorescence 20x images: https://open.quiltdata.com/b/allencell/tree/aics/nuc-morph-dataset/hipsc_fov_nuclei_timelapse_dataset/hipsc_fov_nuclei_timelapse_data_for_pretraining_vision_transformer/
- hiPSC FOV-nuclei timelapse data used for analysis containing FOV bright-field and fluorescence 20x images, with 3D nuclear segmentation images as a separate file: https://open.quiltdata.com/b/allencell/tree/aics/nuc-morph-dataset/hipsc_fov_nuclei_timelapse_dataset/hipsc_fov_nuclei_timelapse_data_used_for_analysis/

- Baseline colonies FOV timelapse dataset: https://open.quiltdata.com/b/allencell/tree/aics/nuc-morph-dataset/hipsc_fov_nuclei_timelapse_dataset/hipsc_fov_nuclei_timelapse_data_used_for_analysis/baseline_colonies_fov_timelapse_dataset/
- DNA replication inhibitor FOV timelapse dataset: https://open.quiltdata.com/b/allencell/tree/aics/nuc-morph-dataset/hipsc_fov_nuclei_timelapse_dataset/hipsc_fov_nuclei_timelapse_data_used_for_analysis/dna_replication_inhibitor_fov_timelapse_dataset/
- Nuclear import inhibitor FOV timelapse dataset: https://open.quiltdata.com/b/allencell/tree/aics/nuc-morph-dataset/hipsc_fov_nuclei_timelapse_dataset/hipsc_fov_nuclei_timelapse_data_used_for_analysis/nuclear_import_inhibitor_fov_timelapse_dataset/
- Feeding control FOV timelapse dataset: https://open.quiltdata.com/b/allencell/tree/aics/nuc-morph-dataset/hipsc_fov_nuclei_timelapse_dataset/hipsc_fov_nuclei_timelapse_data_used_for_analysis/feeding_control_fov_timelapse_dataset/
- Fixed control FOV timelapse dataset: https://open.quiltdata.com/b/allencell/tree/aics/nuc-morph-dataset/hipsc_fov_nuclei_timelapse_dataset/hipsc_fov_nuclei_timelapse_data_used_for_analysis/fixed_control_fov_timelapse_dataset/
- PCNA + DNA dye FOV timelapse dataset (this set has no segmented nuclei): https://open.quiltdata.com/b/allencell/tree/aics/nuc-morph-dataset/hipsc_fov_nuclei_timelapse_dataset/hipsc_fov_nuclei_timelapse_data_used_for_analysis/pcna_dna_dye_fov_timelapse_dataset/

hiPSC single-nuclei timelapse analysis datasets containing DataFrames with quantitative features of tracked segmented nuclei: https://open.quiltdata.com/b/allencell/tree/aics/nuc-morph-dataset/hipsc_single_nuclei_timelapse_analysis_datasets/

- Baseline colonies unfiltered feature dataset (for exploratory purposes)
- Baseline colonies analysis dataset
- Full-interphase analysis dataset
- Lineage-annotated analysis dataset
- Feeding control analysis dataset
- DNA replication inhibitor analysis dataset
- Nuclear import inhibitor analysis dataset

hiPSC nuclei image datasets for training deep-learning models: https://open.quiltdata.com/b/allencell/tree/aics/nuc-morph-dataset/hipsc_nuclei_image_datasets_for_training_deep_learning_models/

- Segmentation decoder training FOV dataset contains paired 20x and 100x FOV TIFF images with companion 100x watershed segmentations (used as ground truth) and model segmentation predictions: https://open.quiltdata.com/b/allencell/tree/aics/nuc-morph-dataset/hipsc_nuclei_image_datasets_for_training_deep_learning_models/segmentation_decoder_training_fov_dataset/
- Interphase detector single-nuclei image dataset contains sequences of maximum intensity projections of 3D images at the single nucleus level used for training the interphase detector: https://open.quiltdata.com/b/allencell/tree/aics/nuc-morph-dataset/hipsc_nuclei_image_datasets_for_training_deep_learning_models/interphase_detector_single_nuclei_image_dataset/

Timelapse Feature Explorer datasets containing maximum projections of 3D FOV-nuclei segmentations together with analysis dataset features, formatted as input for Timelapse Feature Explorer: https://open.quiltdata.com/b/allencell/tree/aics/nuc-morph-dataset/timelapse_feature_explorer_datasets/

- Baseline colonies dataset
- Full-interphase dataset
- Lineage-annotated dataset
- Exploratory dataset

Custom written code was central to the analysis and conclusions of this paper. All necessary code to reproduce the results of this paper have been shared publicly on GitHub. The released custom code repositories use the following Python packages in parts: Jupyter,^26^ numpy,^27^ ipython,^28^ pandas,^29,30^ scipy,^31^ qhull,^32^ matplotlib,^33^ seaborn,^34^ imageio,^35^ vtk,^36^ bigtree,^37^ Colour displays for categorical images,^38^ opencv,^39^ shapely,^40^ dask^41^ and hydra^42^.

- Original/source data for figures in the paper are available in GitHub: https://github.com/AllenCell/nuc-morph-analysis
- The Vision Transformers are implemented using CytoDL, a Python package that we developed for configurable 2D and 3D image-to-image deep learning transformations and representation learning, which is available at https://github.com/AllenCellModeling/cyto-dl
- Code to perform nuclear instance segmentation, formation and breakdown classification, and extraction of quantitative shape features are deposited in morflowgenesis GitHub repository: http://github.com/AllenCell/morflowgenesis
- Configuration files to execute image processing workflows have been deposited on quilt: https://open.quiltdata.com/b/allencell/tree/aics/nuc-morph-dataset/supplemental_files/
- Code used to track single nuclei over timelapse movie: http://github.com/AllenCell/aics-timelapse-tracking
- Code for custom Paraview plugin used to support manual curation of nuclear trajectories: http://github.com/AllenCell/aics-track-curator
- Code used to create the Timelapse Feature Explorer, an open-source, web-based viewer developed for interactive visualization and analysis of segmented time-series microscopy data: http://github.com/allen-cell-animated/nucmorph-colorizer.
- Code used to prepare data for the Timelapse Feature Explorer is available in the http://github.com/AllenCell/nuc-morph-analysis and http://github.com/allen-cell-animated/colorizer-data
- Additional code used to support this paper are available and referenced in the resource table
- Any additional information required to reanalyze the data reported in this paper is available from the lead contact upon request.

### EXPERIMENTAL MODEL DETAILS

#### Cell lines and cell culturing

The two cell lines used in this study are described above. Briefly, the lines were created as described in Roberts et al., 2017^15^, by gene editing the parental WTC11-hiPS cell line derived from a healthy male donor,^25^ using CRISPR-Cas9 to achieve endogenously encoded tagging of proteins representing specific cellular structures. The identity of the unedited parental line was confirmed with short tandem repeat (STR) profiling testing (29 allelic polymorphisms across 15 STR loci compared to donor fibroblasts (https://www.coriell.org/1/AllenCellCollection). Since WTC-11 is the only cell line used by the Allen Institute for Cell Science, edited WTC-11 cells were not re-tested because they did not come into contact with any other cell lines.

The culture and handling protocols for hiPS cell lines were internally approved by an oversight committee and all procedures performed in accordance with the National Institutes of Health, National Academy of Sciences, and Internal Society for Stem Cell Research guidelines. Cells were cultured by hand in 6 cm dishes (Corning, 353002) or in a 6-well plate (Corning, 353046), on an automated cell-culture platform developed on a Hamilton Microlab STAR Liquid Handling System (Hamilton Company).^2,43^ Cells were maintained at 37 °C and 5% CO^2^ in mTeSR1 medium with and without phenol red (STEMCELL Technologies 85850, 05876), supplemented with 1% penicillin– streptomycin (Thermo Fisher Scientific, 15070063) on surfaces pre-coated with growth factor-reduced Matrigel (Corning, 356231). Media was changed (cells fed) daily. When cells reached a confluency of 70-85% (every three or four days), they were dissociated into single cells using Accutase (Gibco, A11105-01) and then re-seeded in supplemented mTeSR1 medium with 1% penicillin–streptomycin and a 10 μM concentration of the ROCK inhibitor Y-27632 (Stem Cell Technologies, 72308). Further details for cell culture reagents and consumables can be found in the Resource Table and standard protocols can be found at www.allencell.org.

#### Cell culture for image acquisition

##### Baseline conditions mEGFP-tagged lamin B1 expressing cells

For imaging, cells expressing mEGFP-tagged lamin B1 were plated on growth factor-reduced Matrigel-coated glass-bottom, black-skirt, 96-well plates with 1.5 optical grade cover glass (Cellvis, P96-1.5H-N). Cells were plated at a density ranging from 1,300 to 1,800 cells per well and cultured for three days. Roughly 1-3 hours before timelapse imaging, cells were fed with a mTeSR1 medium without phenol red that had been pre-equilibrated at 37°C and 5% CO^2^ for between 30-90 minutes. Cells were then moved to the microscope incubation for imaging. No media change occurred during the acquisition over two days.

##### Feeding control conditions

In normal culturing conditions, cells are “fed” daily with fresh media. In the standard two-day timelapse condition described above, however, cells are not fed with new media for the full two days once imaging begins. This means that any of the following could be occurring to the media: the cells are reducing growth factor concentrations, the cells are metabolizing the media which could reduce nutrient availability, and the cells are producing waste. We also know that incubation systems are prone to evaporation (on our system evaporation of 10-30 µL is possible) meaning the media volume could be reduced up to 80% of its original volume, causing a significant change in concentration of media components. To address these possibilities, we prepared cells for comparison between the following four media conditions: the control condition, which is identical to the condition of the three baseline colonies above. The pre-starved condition, where cells are not fed at the beginning of the experiment. This means the last time the cells were fed is 24 hours prior to the imaging experiment. The pre-starved re-fed condition, where cells are not fed at the beginning of the experiment but are re-fed during the timelapse acquisition after 21.5 hours, and the re-fed condition where cells are fed at the beginning and at the middle of timelapse acquisition (21.5 hours).

##### Cells treated with nuclear import inhibitor importazole

Cells were plated with the baseline conditions above. To test the effect of import on nuclear growth, importazole (Sigma-Aldrich 401105), an inhibitor of nuclear import,^44^ was added to the cells at a concentration of 127.3 µM approximately 3 hours into the timelapse acquisition. Control wells were treated by mixing the media already in the well.

Importazole was added to the cells in the following way. The stock solution of the inhibitor was diluted in cell culture media to 9.5x the final desired concentration and 20 µL of this solution was put into a well of an empty plastic 96-well cell culture plate. This plate was then taken to the microscope. To add the inhibitor to the cells, the microscope was opened, 75 µL of media was removed from each well where inhibitor was to be added with a multichannel pipette. These 75 µL were pipetted into the individual wells containing the 20 µL of inhibitor solution and mixed (this is a 4.75x dilution). Next 75 µL of the mixed inhibitor solutions were removed and added back into the same wells the media was taken from (75 µL inhibitor solution added to the remaining 75 µL of cell culture media) and mixed by pipetting up and down 3x times. This last step is a 2x dilution bringing the final dilution to 9.5x.

##### Cells treated with DNA replication inhibitor aphidicolin

Cells were plated with the baseline conditions above. To test the effect of DNA replication on nuclear growth, aphidicolin (Sigma-Aldrich, 178273), an inhibitor of DNA replication,^45^ was added to the cells at a concentration of 20.2 µg/mL approximately 3 hours into the timelapse acquisition following the same procedure described above. Control colonies were treated by the addition of media only (without drug) and mixed.

##### Cells treated with additional small molecule inhibitors for pretraining data purposes

To expand the range of nuclear morphologies for pretraining of the Vision Transformer for nuclear segmentation, we treated cells with additional small molecule inhibitors. For these data, cells were plated with the baseline conditions above. In addition to the importazole and aphidicolin conditions described above, additional importazole (127 and 47.2 µM) and aphidicolin (4.8, 5.9, 11.2 and 20.2 µg/mL) concentrations as well as the following additional small molecule inhibitors were collected: puromycin (2 µg/mL; ThermoFisher, A1113803), rapamycin (9.1 nM; Selleckchem, S1039), and 2-aminopurine (25 µg/mL; Sigma Aldrich, A3509). Inhibitors were added to cells as described above in “Cells treated with nuclear import inhibitor importazole”. Control cells were treated with the addition of the media only and mixed, by mixing only, or nothing was done.

##### Cells treated with DNA replication inhibitor for pulse-chase experiment

To test aphidicolin’s efficacy, we designed the following pulse chase experiment. Cells were plated with the baseline conditions above. Cells were treated with 5-ethynyl-2’-deoxyuridine (EdU,10 µM) for 30 minutes to label active sites of replication. After 30 minutes cells were rinsed and given fresh media without (control) or with 25 µg/mL aphidicolin (treatment). Cells were then incubated for 4 hours at which point they were fixed with 4% paraformaldehyde (PFA) for 15 minutes. The Click-iT™ Edu protocol was followed to label Edu with Pacific Blue Azide (ThermoFisher Scientific, C10418). The cells were then fixed with cold (4°C) methanol for 5 minutes, and then stained with a 1:2000 dilution of mouse-anti-PCNA (Cell Signaling, 2586S) followed by staining with 1:500 dilution of goat anti-mouse AlexaFluor 568 (ThermoFisher Scientific, A-11031).

##### Fixed cells used in training and validating segmentation model

Cells were plated with the baseline conditions above, but with a matrix of increasing densities and varied feeding time to create a fixed-cell sample for training a segmentation model that represented the same wide range of nuclear morphologies observed in the acquired baseline timelapse movies. Along one axis, cells expressing mEGFP-tagged Lamin B1 were plated at a density of 1,500, 3,000, and 5,000 cells per well. Along the other axis, cells were last fed 1 hour, 1 day or 2 days prior to fixation (Supplemental Fig. S1A). To test the segmentation model accuracy, cells were plated with baseline conditions and then fixed.

Cells were fixed with 4% PFA (Electron Microscopy Sciences 15710) in 1x DPBS -/-(Thermo Fischer Scientific, 14190) that had been pre-warmed to 37°C for ∼ 7 minutes. Cells were removed from the incubator, media was aspirated and PFA was added directly. Cells were incubated in the PFA for 10 minutes at room temperature. The PFA was then removed, and cells were washed with 1x DPBS -/-. The fixed plate was either moved directly to the microscope or wrapped in parafilm and stored at 4℃. The plate was warmed to and maintained at 37°C during imaging.

##### PCNA EGFP expressing cells with DNA dye to determine timing of S-phase

Cells expressing mEGFP-tagged PCNA were plated on Matrigel-coated glass-bottom, black-skirt, 24-well plates with 1.5 optical grade cover glass (Cellvis, P24-1.5H-N). Cells were plated at a density of 15,000 cells per well and cultured for three days. Cells were fed with a mTeSR1 medium without phenol red. One hour before moving cells to the microscope, a DNA dye, Spy650 (Spirochrome SC501), was added to a final concentration of 0.1x.

### METHODS DETAILS

#### Image acquisition

##### Spinning-disk confocal microscopy

Imaging was performed on ZEISS spinning-disk confocal microscopes equipped with a 1.2x tube lens adapter with 10x/0.45 NA Plan-Apochromat, 20x/0.8 NA Plan-Apochromat, or 100x/1.25 NA W C-Apochromat Corr M27 objectives (ZEISS) for final magnifications of 12x, 24x or 120x, respectively. Image acquisition routines were designed in the ZEN 2.3 or 2.6 software (blue edition; ZEISS). The spinning-disk confocal microscopes were equipped with a CSU-X1 spinning-disk scan head (Yokogawa) and two Orca Flash 4.0 cameras (Hamamatsu). A custom-made long pass (LP) dichroic beam splitter 660LP-3mm (Chroma) was used to split the emission light path to the two cameras. The microscope stage was outfitted with a humidified environmental chamber (Pecon) to maintain the cells at 37 °C with 5% CO^2^ during imaging. A Prior NanoScan Z 100 mm piezo Z stage (ZEISS) was used for fast acquisition in z. Optical control images were acquired regularly to monitor microscope performance. Laser power at the objective was measured monthly. Unless otherwise specified, a standard 488 nm laser line was used at a laser power of 0.271 mW (measured with 10x objective). An Acousto-Optic Tunable Filter (AOTF) was used to modulate the intensity of the laser. A 525/50 nm Band Pass (BP) filter set (SEMROCK) was used to collect emissions from EGFP. Images were acquired with an exposure time of 50 ms unless otherwise specified. Transmitted light (bright-field) images were acquired using a red LED light source (Sutter TLED+) with a peak emission of 740 nm with a narrow range and a BP filter 706/95 nm (Chroma) for bright-field light collection. The nominal Z-step size for 3D image stacks was 0.53 µm for the 20x/0.8 NA objective and 0.29 µm for the 100x/1.25 NA objective. In timelapse imaging experiments, 3D image stacks were acquired at 5-minute intervals. Each stack consisted of 42 image planes (i.e. Z-slices) which acquired the EGFP fluorescence emission and bright field transmitted light simultaneously on two separate cameras split by the 660LP-3mm beam splitter. Focus was maintained with Definite Focus 2 (ZEISS).

Due to a refractive index mismatch from imaging with a 20x/0.8 NA air objective (RI=1.000) into cells and aqueous media (RI=1.360-1.380 and RI=1.333, respectively), there exists a spherical aberration-based axial distortion that makes objects appear flatter in Z than they are. We determined the axial distortion correction factor on these microscopes to be ∼1.43 using the axial correction macro provided by Diel et al., 2020^46^. This means that while the nominal (i.e., set in the software) Z-step size for the 20x/0.8 NA Plan-Apochromat objective was 0.53 µm, the actual Z-step size was 1.43 times larger or about 0.758 µm (for explanation see Diel et al., 2020^46^). We confirmed that this rescaling factor of 1.43 achieved good alignment between the tops and bottoms of thick samples imaged with both the 20x/0.8 NA Plan-Apochromat objective and the 100x/1.25 W C-Apochromat Korr UV Vis IR objective.

##### Acquisition of multi-day, 3D timelapse movies of mEGFP-tagged lamin B1 expressing cells

A total of nine colonies of hiPS cells (see “Baseline conditions for lamin B1 EGFP expressing cells”) were imaged as described above every 5 minutes for 47 hr and 25 min. This resulted in nine timelapse movies of cells growing from day 3 to day 5 after initial plating.

A total of ten multi-day timelapse movies were acquired to determine the effect of possible nutrient depletion (see “Feeding control conditions” above). 20x colony positions were chosen to optimize colonies of matched size between the control condition and the re-fed, pre-starved, and pre-starved-refed conditions. This resulted in four control scenes, three re-fed scenes, one pre-starved scene and two pre-starved re-fed scenes. To maintain consistent microscope focus and incubation temperature, the plate containing cells was not removed from the microscope during the re-feed. Instead, media was added to the cell plate while on the microscope stage, by opening the incubation chamber and pipetting the inhibitors directly. The system was then closed gently as soon as possible, such that the incubation chamber was open ∼2-3 minutes. Cells were otherwise imaged as described above every 5 minutes for 47 hr and 55 min.

##### Acquisition of importazole, aphidicolin, and inhibitor-treated lamin B1 EGFP expressing cells

A total of 44 colonies of hiPS cells expressing EGFP-tagged lamin B1 were imaged across three separate inhibitor treatment experiments (see “Cells treated with small molecule inhibitors producing varied nuclear morphologies”). To ensure that imaging duration was sufficiently long to capture nuclear growth behavior both before and after inhibitor addition, the timelapse experiments were acquired for 3 hours before and after inhibitor addition. Similarly to the addition of media described above for the feeding control experiments, cells were not removed from the microscope during addition of inhibitors. Instead, inhibitors were added to the cell plate while on the microscope stage, by opening the incubation chamber and pipetting the inhibitors directly. The system was then closed gently as soon as possible, such that the incubation chamber was open ∼2-3 minutes.

The first timelapse contained 15 colonies with the following conditions: aphidicolin at 20.2, 11.2 and 5.9 µg/mL, rapamycin at 9.1 nM, and a media-only control. The cells were imaged for 185 minutes before inhibitor addition and then an additional 240 minutes after for a total of 425 minutes (7 hours and 5 minutes) The second timelapse contained 15 colonies with the following conditions: importazole at 127 and 47.2 µM, 2-aminopurine at 25 µg/mL, and puromycin at 2 µg/mL, and a mixing-only control. The cells were imaged for 220 minutes before inhibitor addition and then an additional 325 minutes after for a total of 545 minutes (9 hours and 5 minutes). The third timelapse contained 14 colonies imaged under the following conditions: aphidicolin at 20.2 and 4.8 µg/mL, media-only control, and a control where nothing was done (no media or mixing). The cells were imaged for 175 minutes before inhibitor addition and then an additional 570 minutes after for a total of 745 minutes (12 hours and 25 minutes).

##### Acquisition of DNA replication inhibition pulse-chase experiment

To test the efficacy of DNA replication inhibition for the duration of the previous timelapse experiment, we designed a pulse chase experiment (Cells treated with DNA replication inhibitor for pulse-chase experiment). The fixed cells were imaged on using the Zeiss spinning disk confocal systems described above using a 100x/1.25 NA W C-Apochromat Corr M27 objectives (ZEISS) and 405 and 561 laser lines with 450/50 nm and 600/50 nm filters. The colocalization of PCNA (anti-PCNA 561) with Edu (Pacific blue 405) in the treatment condition indicates that aphidicolin causes rapid and sustained inhibition of DNA replication progression (Supplemental Fig. S13).

##### Acquisition of matched pairs of 20x and 100x training data images

To create a nuclear segmentation model that works on 20x images trained via 100x matched pairs, we need to capture pairs of 20x and 100x images of the same fixed cells. Because fixation reduces the mEGFP fluorescence intensity, the 488 laser power was increased from 0.271 mW for 20x imaging so that the pixel-intensity histograms from the images matched those from the 20x live-cell timelapse images. For the 100x images, the 488 laser power was set to ∼2.3 mW (as used in Viana et al., 2023^2^) to facilitate accurate lamin B1 segmentation. The 20x positions were chosen to maximize the variability in nuclear size (small, large, flat, tall, etc) and shape (wrinkly, oblong, smooth, etc) and colony densities (dense, thick, spread, etc) so the model would see as much of the natural variation as in the acquired timelapse images (Supplemental Fig. S1A). A total of 80 20x positions were acquired, with 9 100x positions tiled across each 20x FOV (Supplemental Fig. S1B) yielding 720 paired image regions.

##### Acquisition of fixed control timelapse

To measure the consistency in the model-produced nuclear segmentations and downstream volume measurements, we created a fixed plate of hiPS cells expressing mEGFP-tagged lamin B1. We imaged the same colony of cells 20 times, translating the sample 2.5 µm in the X,Y plane between each acquisition to simulate the distance nuclei move in a typical 5 minute interval. Images were taken with the same image acquisition settings as the timelapse images, except the 488 laser power was increased slightly from 0.271 mW so that the image pixel-intensity histograms matched that of the timelapse acquisition image histograms.

##### Acquisition of PCNA with DNA dye to determine timing of S-phase

Cells were imaged on a Zeiss Lattice Light sheet 7 microscope equipped with a 44.83x/1.0 NA detection objective, laser lines with wavelengths of 488 nm and 640 nm and an LBF405/488/561/640 laser blocking emission filter. Lightsheet setting 30x1000 was used to perform the imaging. 488 nm and 640 nm laser power used for imaging were approximately ∼11.6 µW and ∼36.5 µW (measured at the meniscus lens), respectively. The images of PCNA and DNA dye were acquired sequentially (using 488 and 640 nm excitation, respectively) with 10 ms exposures on an ORCA Fusion sCMOS camera (Hamamatsu), taking 0.4 µm steps while stage-scanning through the sample. This image acquisition was repeated at 108 s intervals for a total duration of 6 hours.

#### Creation of FOV-nuclei timelapse datasets

##### Measure of initial colony size

To measure the initial colony size, the boundary of the colony is segmented from the first timepoint of the bright-field image. To crop the z-stack to the slices with the most colony information, the approximate top and bottom of the colony were detected using the coefficient of variation (CV) profile computed from the standard deviation/mean of each Z-slice. The minimum in the CV profile is the in-focus, middle slice of the colony. The maxima above and below the center slice are the approximate tops and bottom of the cells. The Z-stack is cropped to contain only the slices between the top and bottom of the cells. A standard deviation Z-projection of the image stack is made and normalized by the mean intensity of the whole cropped Z-stack. A Sobel filter (implemented in OpenCV) using first order derivatives with a kernel size of 5 is applied to the image and thresholded (threshold=0.025) to binarize.^39^ Then a series of post-processing steps are applied to remove small objects <734 µm^2^, fill holes, apply gaussian smoothing and binarize, and fill holes again. The pixel size in the 20x/0.8 NA bright-field image is 0.271 µm/pixel and the area of the resulting colony segmentation mask is used as a measure of initial size.

##### Baseline colonies FOV dataset

Three of the nine colony movies were selected for downstream image processing and creation of the baseline colonies analysis dataset. They were selected based on several key criteria, including that they displayed different starting sizes, stayed centered within the field of view, and the fraction of cells coming from colonies outside the FOV is less than 10%. The three colonies are referred to as “Small,” “Medium,” and “Large” to indicate their relative starting sizes (approximately 31,500, 63,000, and 110,800 µm^2^, respectively).

##### Feeding control FOV dataset

Three of the ten colony movies acquired under different feeding control conditions were selected for downstream image processing and creation of the feeding control dataset based on colony size and analysis tractability. A representative colony for each of the following media conditions was chosen: a control matching the baseline condition, a re-fed condition, and a pre-starved condition (see “Feeding control conditions”). The pre-starved, re-fed condition was not chosen for analysis due to significant stage jitter that caused downstream tracking issues. To control for variation in colony development, the colonies were chosen to be similar in starting size with initial an initial colony area of approximately 51,700, 44,000, and 40,500 µm^2^ for the control, re-fed and pre-starved conditions, respectively.

##### DNA replication inhibitor FOV dataset

Five colonies were chosen for downstream image processing and creation of the DNA replication inhibitor FOV dataset. Two colonies treated with 127.3 µM and three control colonies treated with mixing only were chosen. Colonies were selected such that the aphidicolin treated colonies had a paired control colony of a similar size (initial colony area of approximately 30,600, 32,400 µm^2^, and 42,900, 47,400 µm^2^ for each aphidicolin, control pair, respectively). The control and aphidicolin treated colony pairs were chosen from the same inhibitor timelapse acquisition. An additional control colony with initial colony area of ∼77,500 µm^2^ was used to control across experiment days.

##### Nuclear import inhibitor FOV dataset

Two colonies of the same size (initial colony areas of approximately 109,900 and 143,000 μm²) and from the same inhibitor timelapse acquisition were chosen for downstream image processing and creation of the nuclear import inhibitor FOV dataset. One colony was treated with 127.3 µM approximately 3 hours into the timelapse acquisition and a control colony treated with mixing only.

##### Fixed control timelapse FOV dataset

The fixed control FOV dataset contains all 20 frames acquired of the same fixed colony, translated between each frame, to create a pseudo timelapse.

##### PCNA + DNA dye FOV dataset

Raw images were deskewed in ZEN Blue 3.7 using “Coverslip Transformation’’ option. Nuclei on the colony’s edge were much brighter due to significant uptake of the DNA dye. Nuclei in the center of the colony that were dim and did not uptake a significant amount of DNA dye were manually tracked for analysis (Supplemental Fig. S4). This timelapse dataset was used to measure the timing of S-phase (Supplemental Fig. S4).

##### Interphase detector training single-nuclei image dataset

To train a model to detect nuclei in interphase (Supplemental Fig. S14), we created 64x64 maximum intensity projections of 505 single cell trajectories from the Large colony.

##### Pretraining FOV dataset

To create a robust, generalizable 3D segmentation model for nuclei via lamin B1, a large pretraining dataset comprised of timelapse data was created. This dataset includes all 19 multi-day, 3D timelapse movies of mEGFP-tagged lamin B1 expressing cells colonies corresponding to all of the baseline and feeding control conditions, as well as all 44 3D timelapse movies of mEGFP-tagged lamin B1 expressing cell colonies from the importazole, aphidicolin, and additional inhibitor-treated timelapse experiments, resulting in 63 timelapse movies for pretraining.

##### Segmentation decoder training FOV dataset

Training a segmentation model to produce 100x segmentations from 20x images requires that the training data is accurately aligned between imaging modes. We registered the images based on translational transformations only, as opposed to rotational, shearing, or elastic transformations that might distort the image data through interpolation. In rescaling images of different resolutions to each other, rounding errors can occur from multiplying by a pixel size ratio with decimals. To ensure rounding errors do not introduce issues in registration after rescaling, we crop the 20x and 100x images such that the ratio of dimensions between the images match the ratio in pixel sizes as closely as possible.

We extracted nine tiles from each 20x FOV corresponding to the regions where 100x images were acquired (Supplemental Fig. S1A). Registration and rescaling was performed in three steps. First, the 100x image was downsampled to 20x resolution and the maximum intensity projections (MIPs) of the image pairs were registered using ORB feature detection^47^ RANSAC optimization^31^ to estimate a 2D Euclidean transformation that aligns the images in the XY-plane. Based on this alignment, the 20x image was then automatically cropped to be 5 pixels larger than the downsampled 100x image in the X and Y dimensions. We then used gradient descent optimization to determine an optimal 3D translation that maximizes the Mattes Mutual Information^48^ between images at the 20x scale. We refined this 3D registration by performing the optimization a second time with both images scaled to 100x resolution, using the previous alignment as the initial starting position. The results of this final registration were used to crop the 100x images (maximally by 15 pixels in XY and 1 pixel in Z) and the 20x image such that it can be upsampled to 100x resolution and be accurately aligned when overlaid with its corresponding 100x image with identical image/pixel dimensions. The results of this registration process were manually curated by examining each image pair overlaid on top of each other to ensure all final data is accurately aligned, resulting in 410 20x-100x pairs in the segmentation decoder training FOV dataset.

#### Generating a Vision Transformer-based 3D segmentation model for nuclei via lamin B1

##### Generating lamin B1 3D segmentation ground truth data from 100x images

The 100x images from the aligned 20x-100x pairs in the Segmentation decoder training FOV dataset were used to generate high quality ground truth segmentations for 20x/0.8 NA 3D images of lamin B1. We performed a watershed-based segmentation on the 100x images by first clipping the fluorescence profiles at the 1st and 99th percentile, then smoothing in the XY-plane for each Z-slice using a Gaussian filter with unitary standard deviation, then finally, performing the watershed with manually placed seeds. We evaluated all segmentations manually by overlaying the contours of the watershed segmentations onto the 100x raw images. Segmentations were considered inaccurate and not used if the segmentation did not fall directly along the midline of the nuclear boundary (as indicated by mEGFP-tagged lamin B1 fluorescence) at all points across all angles (top view and both side views; Supplemental Fig. S15). Resulting in 410 labeled 20x/100x segmentation pairs split 372/20/18 for training, validation, and testing respectively.

##### Vision Transformer encoder pretraining on nuclear timelapse images

Obtaining high quality segmentations posed a significant challenge due to the limited number of labeled 20x/100x segmentation pairs and the domain gap between the few fixed 20x images (e.g., significant image-to-image differences in features like signal to noise ratio). For these fixed images we can generate a paired 100x segmentation ground truth, whereas for the numerous live 20x images, collecting paired ground truth data is infeasible. Experiments with fully convolutional models trained in a supervised manner exhibited poor generalization (data not shown). Noting the large quantities of unlabeled live 20x movies, we instead opted to perform self-supervised pretraining of a Vision Transformer (ViT^16^; Supplemental Fig. S2A and B) encoder on live 20x images followed by supervised training of a convolutional decoder on 20x/100x matched image pairs to generate high-resolution instance segmentations of the nucleus via lamin B1.

We first pretrained a Masked Autoencoder (MAE)^49^ using a XYZ patch of size 8x8x4 voxels, a mask ratio of 0.75, and learnable positional embeddings.^50^ The encoder is made up of 12 identical ViT blocks^51^ each with eight heads and an embedding dimension of 512. We use a single-layer transformer decoder with 8 heads and an embedding dimension of 128. Following CrossMAE,^50^ only the masked tokens in the input image were reconstructed by the decoder through cross attention from a set of mask tokens with learned positional embeddings to the output of the Inter-block Attention (a learned weighting of the encoder’s intermediate representations). After the transformer decoder, tokens were linearly projected and rearranged back into an image and mean squared error (MSE) loss was calculated between input-masked tokens and reconstructed images.

We used all timepoints from 63 timelapse movies in the pretraining FOV dataset for MAE training, corresponding to 19,692 fields of view for training and 50 for validation. As the full dataset was too large to fit into memory, we cached 10% of the dataset for training a single epoch and replaced 15% of the cache per epoch. To augment and normalize data, we used transforms from the MONAI library.^52^ First, we Z-scored each FOV’s intensity distribution independently by subtracting its mean and dividing by its standard deviation. We then randomly extracted 96 patches of size 192x192x24 voxels from each FOV to form a single batch that was augmented using random flips across the X and Y axes, and random 90-degree rotation around the z axis. The model was trained for 1000 epochs over four days on a single A100 GPU using PyTorch^53^ implementations of the AdamW optimizer^54^ with a weight decay of 0.05 and a OneCycle learning rate scheduler^55^ with a 10 epoch warmup to a maximum learning rate of 10^-4^.

##### Segmentation decoder training on matched pairs of 20x 100x ground truth nuclear segmentations

After self-supervised pretraining, we froze the image encoder (except for the Inter-lock Attention weights) and replaced the transformer decoder with a multi-scale decoder inspired by the skip connections of the UNET architecture^56^ (Supplemental Fig. S2C and D). Each layer in the decoder increases spatial size while decreasing the number of channels through two 3x3x3 convolutions with residual connections followed by trilinear upsampling by 2x in each spatial dimension. The input to each layer is a distinct output from the inter-block attention (serving as a skip connection) concatenated with the output from the previous decoder layer. The final layer upsamples the Z dimension by 2.6134 and XY dimensions by 2.5005 to match the pixel size of the 100x segmentation image. This is followed by 3x3x3 convolutional block and a 1x1 convolutional block yielding a 6-channel output image. We used a custom 6-channel instance segmentation ground truth format inspired by recent flow field-based instance segmentation methods^57–59^ (Supplemental Fig S. 2D) designed to avoid computationally intensive postprocessing steps. Channel 1 represents the distance transform of a connectivity-preserving erosion of the instance masks, optimized via a spatially weighted MSE Loss emphasizing object boundaries. Channel 2 is a semantic segmentation optimized via Tversky Loss.^60^ Channels 3, 4, and 5 are ZYX gradient fields directed from each point in a 10-pixel shell on the interior of each object boundary to the closest point on the connectivity-preserving eroded object used in Channel 1. These gradient field channels are optimized via the same spatially weighted MSE loss. Finally, Channel 6 is a boundary segmentation optimized via Tversky loss.

During decoder training we randomly selected 3 patches of size 192x192x24 voxels in XYZ from the low-resolution images at 20x and corresponding patches of size 480x480x62 voxels from the high-resolution segmentations at 100x to form a single batch. Training pairs were augmented using random flips across the X and Y axes and the low-resolution images were additionally augmented with random histogram shifts, random intensity shifts, random contrast adjustment, and random Gaussian noise. We used 372 image pairs (generated as described above) for training, 20 pairs for validation, and 18 for testing (see Supplemental Fig. S2).

##### Generating instance segmentations of nuclei

During inference (Supplemental Fig. S2D), we applied the model in a sliding window over a full field of view. Neighboring windows were overlapped by 30x30x4 voxels in XYZ and a Gaussian weighting was used to prevent the occurrence of stitching artifacts in the 6-channel predicted image. To convert the 6-channel model prediction to a single-channel instance segmentation, we first computed rough instance segmentations based on a labeled thresholding of the semantic segmentation. We then refined connected components containing multiple eroded objects (indicating that multiple instances have been merged) using the gradient fields.

##### Validating instance segmentations of nuclei

To evaluate the accuracy of the trained segmentation model, we utilized the test set of 18 pairs of 20x images and 100x segmentations that were not used in training the model. We applied the trained segmentation model to the 20x images to generate instance segmentations that can be compared with the 100x ground truth segmentation (generated with seeded watershed as described in “Generating lamin B1 3D segmentation ground truth data from 100x images”). The accuracy of the segmentations was evaluated by visual inspection of segmentation overlays (Supplemental Fig. S1C) and by comparing quantified features from the segmentations (Supplemental Fig. S1D and E).

We evaluated a wide range of nuclear features including volume, height, surface area, and surface area to volume ratio. The model predicted segmentation gave an unfitted *r*^2^ of >0.9 for all features (Supplemental Fig. S1E) with an RMSE of 20.1 µm^3^ for nuclear volume. Notably, there was almost no over-or under-estimation bias in the predictions–the largest being -9.79% (or -0.4 µm) for nuclear height. We also evaluated the accuracy of the shape of the model-produced segmentations by using a spherical harmonic expansion (SHE) to parameterize each 3D nuclear shape (as in Viana et al., 2023^2^). We performed principal component analysis on the SHEs (as in Viana et al., 2023^2^) to obtain 8principal components for nuclear shape (PC1-8) which were also predicted with a high degree of accuracy (Supplemental Fig. S1E). “Percent error” is defined as the RMSE divided by the 90% interpercentile range (95th percentile value – 5th percentile value). A value of ∼50% will occur for uncorrelated data. “Percent bias” is defined as the average error divided by the 90% interpercentile range.

#### Generating single-nucleus timelapse datasets

##### Segmentation of single nuclei using the Vision Transformer-based deep-learning nuclear segmentation model

The Vision Transformer-based deep-learning nuclear segmentation model was applied to each 20x mEGFP-tagged lamin B1 image chosen for downstream analysis and returns a 100x FOV nuclear segmentation where each nucleus has its own label (as described in “Segmentation decoder training on matched pairs of 20x/100x ground truth nuclear segmentations***”*** and “Generating instance segmentations of nuclei”). From these FOV instance segmentations, we generate crops for single nucleus feature extraction.

##### Single nucleus feature extraction

Quantitative shape features were computed for each segmented nucleus as in Viana et al., 2023.^2^ Briefly, each segmented nucleus was cropped then rescaled to isotropic voxel sizes by interpolating along the z dimension to upscale the voxel size (XYZ) from 0.108333x0.108333x0.29 μm to 0.108333x0.108333x0.10833 μm. Nuclear volume is calculated as the number of non-zero voxels in the single nucleus crop. The single-voxel volume (0.108 μm^3^) was used to rescale this feature into units of µm^3^. Nuclear height is calculated as the distance in voxels along the Z-axis between the bottom-most and top-most voxels in the input cropped segmentation image. The single-voxel height of 0.108 μm was used to rescale this feature to units of µm. To determine nuclear surface area, we first generate a mesh-based representation of the nucleus segmentation. First, because of the lower resolution of the Z dimension relative to X and Y, the binary input image is convolved with a Gaussian kernel with size σx=σy=σz=2, which is enough to smooth the image while retaining the overall nuclear shape, improving the quality of the output mesh that can be derived. Next, this gaussian-smoothed binary segmentation is converted into a 3D triangular mesh using a traditional marching cubes algorithm from VTK Python library.^36^ The surface area of the mesh is then estimated using the GetSurfaceArea function within vtkMassProperties which uses Heron’s formula to calculate and sum over the areas of each triangle in the mesh. The single-voxel-side area of (0.108 μm)^2^ was used to rescale the estimated surface area value into units of µm^2^.

##### Automated tracking of single nuclei

Single nucleus tracking was performed using a segmentation-based object tracking algorithm, using the results of the nuclear instance segmentations and the nuclear volume feature as the input data. Tracking was performed through the solving of an Earth Mover Distance (EMD) problem on a frame-by-frame basis.^61^ Individually segmented nuclei are represented as nodes in two sets, one for objects in the current frame and another for those in the subsequent timepoint. Edges are drawn between nodes in these sets if the centroids of their corresponding objects are within a maximum distance of each other. Each edge is assigned a cost which is proportional to the distance between centroids and difference in volume of the segmented objects. Each node is also connected to a set of “virtual” nodes not corresponding to a segmented nucleus, representing the condition that the nucleus is missing in one of the frames due to mitosis, segmentation error, or the nucleus leaving the field of view. The optimization problem seeks to distribute a fixed amount of weight across all edges while minimizing the cost contributed by each edge. The edges which are assigned the greatest amount of weight are determined to be connecting instances of the same nuclei between frames. This process is performed for each pair of subsequent frames in the timelapse until all segmented objects are connected in continuous trajectories.

The last step in the tracking workflow attempts to link trajectories that were broken due segmentation errors. Because we prevent links being made between nuclei of significantly different volumes or centroid positions, a nuclear trajectory could end prematurely due to a segmentation error, e.g. no nucleus is segmented or two nuclei are segmented as one. To enable nuclear trajectories to be linked across this problematic frames, we performed a second round of tracking in which the optimization is performed only between objects in the current timepoint which have no link to the next timepoint and objects from the next three timepoints with no trajectory links to previous timepoints. Objects which touch the edges of the image are excluded from this step, due to the possibility that their tracking has ended due to leaving the field of view (FOV). Importantly, this linking skips the frame with the segmentation error, so that nuclear trajectory will not include the bad segmentation from the problematic timepoint; instead, all nuclear features at that missing timepoint will shown as NaN values.

##### Automated interphase detector

We used an image-based classifier to demarcate the beginning and end of interphase for single nucleus trajectories. To train the model, we identified the first timepoint with a fully formed lamin B1 shell (formation) and the last timepoint with an intact lamin B1 shell (breakdown) through visual inspection of 64x64 maximum intensity projections (MIPs) of 505 single cell trajectories from the Large colony (Interphase detector training single-nuclei image dataset). We assigned each timepoint in a trajectory with a 0 if the lamin shell is intact (interphase), and if it is 1 if it is broken, resulting in ∼65,000 labeled single-cell crops used for training a convolutional classifier (see Supplemental Fig. S14). We used random flipping, rotation, intensity shifts, and cropping and resizing transforms during training of a simple regressor convolutional network^52^ with three blocks, each of which performs convolution (with 8, 16, and 32 filters respectively) and 2x downsampling, followed by a linear layer with two outputs. We trained the network for 40,000 steps using a OneCycle learning rate with a max learning rate of 3×10^-4^, weight decay of 0.01, and cross entropy loss. Despite significant class imbalance (97.4% of examples have a fully formed lamin shell), the model achieves a macro-averaged F1 score of 0.98 on the validation set.

To apply the trained model to a single cell trajectory, we extracted crops using a bounding box centered on the nuclear instance segmentation over time, created a MIP and resized them to 64x64 pixels. We temporally padded the beginning and end of each trajectory by three timepoints by extending the coordinates of the first and last crops. To make a prediction, we take the class argmax of the model predictions across the trajectory and define lamin shell “formation” and “breakdown” as the first and last timepoints where the nucleus was classified as having an intact lamin shell, respectively (Supplemental Fig. S14B). If no such transition between the two classes (intact or broken lamin shell) occurs, the trajectory is marked as not having a formation or breakdown timepoint.

### QUANTIFICATION AND STATISTICAL ANALYSIS

#### Dataset curation and filters

##### Trajectory quality control filters (automated)

Nuclear segmentations that touch the edge of the FOV are excluded from analysis, as they do not capture the entirety of the nucleus (this is flagged by the “fov_edge” column in the manifest). Nuclei with trajectories shorter than five frames are excluded from all analyses, as these were found to often be debris or incidents of merged object segmentations. Nuclear trajectory data from frames when nuclear volumes deviate by greater than 15% from the median volume within a rolling window of 15 frames (75 minutes) for a given nuclear trajectory are also excluded from analysis. These deviations are indicative of segmentation error, usually associated with merge events, such as cell debris being included in a nuclear segmentation. When calculating this rolling-window median volume, the beginning of the trajectory was padded with the first volume value, and at the end of the trajectory was padded with the median of the volumes in the last three frames; this was chosen to prevent the real rapid growth phase from A-B from being called an outlier, while still appropriately identify “outlier” jumps in volume at mitosis at the end of the trajectory. Outliers detected by this automated method are flagged by the “is_tp_outlier” in the manifest. While present, these deviations are infrequent, making up an average of only 0.64% of timepoints in full-interphase trajectories.

##### Baseline cell death annotation and filter (manual)

When cells die, the lamin B1 has a district phenotype where the nuclear envelope blebs into multiple pieces which is visually distinct from lamin shell breakdown in mitosis. These pieces turn into debris that are eventually extruded from the colony. To ensure that the main analysis is restricted to healthy cells, all cell death events were manually annotated in the baseline colony FOV dataset using the custom-built macro (https://github.com/aics-int/aics-track-curator) in Paraview^62^ and removed from the baseline analysis dataset.

##### Calculation of the start of growth

To calculate the transition point between expansion and growth phases in single nucleus trajectories, each potential full-interphase trajectory of volume over time was truncated to the first 40 frames. These truncated trajectories were parameterized as arcs and linearly interpolated to account for missing values (e.g., removed outliers). This parameterization gave uniform sampling along the length of the curve rather than being uniform in time, increasing the sampling density in the region of the transition. Each interpolated arc was fit to a hyperbola defined by two asymptotes which intersect at a center point. The point on the hyperbola (sharing an x-value with the interpolated data) closest to the hyperbola’s center was chosen as the approximated transition point. Then the nearest frame (within two frames) with a successful segmentation was chosen as the transition point, which marks the start of growth (Supplemental Fig. S16).

##### Full-interphase trajectory filter (automated)

A trajectory is considered a full-interphase trajectory when it has both formation and breakdown events predicted (Interphase detector), the transition point is successfully calculated (Supplemental Fig. S16) and the trajectory is at least 120 frames (10 hours) long. These trajectories must pass all downstream feature calculations to be included in the full-interphase analysis dataset. For example, nuclear trajectories cannot have negative growth rates.

##### Baseline full-interphase trajectory quality control annotation and filter (manual)

All full-interphase trajectories derived from the baseline analysis dataset were manually checked for errors which could have incurred during automated segmentation, tracking, formation and breakdown detection, or transition point calculation. Trajectories with errors were identified by their Track ID and manually removed from the dataset (N=27).

##### Baseline annotation and filter for extremely long growth durations (manual)

53 nuclei were identified as having especially long growth durations (∼22-37 hours) compared to the rest of the population (average duration of 14.9 ± 2.0 hours). Many of these nuclei were outliers in nuclear volume fold-change (some up to four-fold) and nuclear ending volume (some up to nearly 2,000 µm^3^). We characterized the fate of these nuclei and found that 11 ended in cell death before completing interphase; 21, while successfully dividing, had daughters that died shortly after interphase; and 11 divided at the end of the timelapse movie so the fate of their daughters could not be observed. The 15 that die before division were annotated as undergoing cell death above and the 32 whose daughters died or had unknown fate were flagged as outliers. While the remaining 10 nuclei divided successfully and had daughters that did not die, they were flagged as outliers since they grew for a long time and had abnormally large fold-changes and ending volumes (i.e. grew similarly to the outliers that ended in death). The nine second generation outliers were also flagged as outliers. Thus, in total 51 full-interphase nuclear trajectories were flagged as long growth duration outliers. We have flagged all 51 of these full-interphase nuclear trajectories as growth outliers because our goal is to understand healthy, normal hiPS cell growth variation on a population and individual level and we cannot rule out that there might be something inherently different about the growth of this subset of nuclei that represents 3.5% of the full-interphase trajectories. (Supplemental Fig. S6).

##### Curation of lineage relationships and single nucleus tracking (manual)

Lineage trees were created for the Small and Medium sized colony movies by manually annotating mother and daughter relationships (i.e. assigning each daughter nucleus with the appropriate ID for its mother). The manual annotations to these trajectories were made using a custom-built macro (https://github.com/AllenCell/aics-track-curator) in Paraview^62^, which allows simultaneous viewing of data and interactive annotation. Maximum intensity projections of the 20x mEGFP-tagged lamin B1 movies with the centroid trajectory for each nucleus overlayed were displayed and single nuclei were followed through time until division, at which point the daughter nuclei would be assigned the appropriate “Parent ID” based on the Track ID of the mother nucleus. Additionally, during this round of annotations, if the trajectory incorrectly switched to a different nucleus at a given frame (e.g. when two nuclei move past one another in close proximity), its trajectory was manually adjusted to follow the correct nucleus. All nuclei in any one lineage tree were assigned a common “Family ID.” These manual lineage annotations and trajectory corrections were then applied to the initial automated tracking output resulting in a total of 316 lineage trees.

#### Analysis datasets generation

##### Baseline colonies analysis dataset

The baseline colony FOV dataset was processed using the following steps. First, the Vision Transformer-based deep-learning segmentation model was applied to each timepoint (e.g., 3D image of mEGFP-tagged lamin B1 fluorescence), transforming the 570 3D 20x images of size 42x1248x1824 into 570 3D instance segmentations of size 108x3120x4560, yielding a total 646,034 individually segmented nuclei (179,223, 206,106, and 260,705 nuclei per movie) (see Fig. 1C). Next, feature extraction and automatic tracking of single nuclei were done resulting in 29,511 trajectories. Finally, to create the baseline colonies analysis dataset, the baseline trajectory quality control filters and the baseline cell death filter were applied resulting in 4,741 trajectories of nuclei.

##### Full-interphase analysis dataset

To create the full-interphase trajectory analysis dataset, both the full-interphase trajectory filter and the baseline full-interphase trajectory quality control filters were applied to the baseline trajectory analysis dataset. This is the dataset containing all quality controlled full-interphase trajectories including ones that grow for abnormally long durations as shown in Supplemental Fig. S6 (full-interphase trajectory with outlier analysis dataset). Lastly, we remove these growth outliers to create the full-interphase trajectory analysis dataset used in the main analysis of this paper.

##### Lineage-annotated analysis dataset

The lineage-annotated analysis dataset is derived from the full-interphase trajectories analysis dataset for the Small and Medium colonies. This subset includes only the full-interphase trajectories that have lineage relationship annotations.

##### Feeding control analysis dataset

The feeding control FOV dataset was processed using the same steps as the baseline dataset (Vision Transformer-based deep-learning segmentation model, feature extraction and automatic tracking of single nuclei). Because the cells were fed directly while on the microscope, the microscope stage shifted beyond the normal range at the 21.5 hour timepoint, causing some trajectories to break with the automatic tracking of single nuclei. A manual shift and track matching step were applied to this frame to rescue the trajectories that were broken. Next the automated trajectory quality control and the full-interphase trajectory filters were applied. No manual curations or annotation filters were applied to the feeding control analysis dataset. The final feeding control analysis dataset contains 316, 384, 377 full-interphase trajectories for the control, re-fed, and pre-starved conditions respectively.

##### DNA replication inhibitor analysis dataset

The five colony movies from the DNA replication inhibitor FOV dataset (3 control and 2 aphidicolin-treated) were processed using the same steps as the baseline dataset (Vision Transformer-based deep-learning segmentation model, feature extraction and automatic tracking of single nuclei). Three colony pairs were matched: the two aphidicolin-treated colonies were paired with an appropriate control colony of similar starting size, and a control pair was created by pairing two control colonies together. Then we applied the automated trajectory quality control filter to remove problematic segmentations (such as those touching the edge of the FOV) or short trajectories. Next, timepoints were removed that were within 15 minutes of the end of interphase (lamin shell breakdown) or 60 minutes after the start of interphase (lamin shell formation) to avoid the rapid changes that occur around mitosis from affecting the analysis. To avoid effects due to cell death, all timepoints after the point at which the 25% of the nuclei in the colony appeared to be dead or dying were removed. Nuclear trajectories were required to be present at least 60 minutes before aphidicolin addition and at least 120 minutes after. To account for any effects due to nuclear size differences, trajectories were selected one by one for analysis such that for each trajectory in the inhibitor condition, a single trajectory from the control condition was selected that had the same average volume within at the time immediately before inhibitor addition. Thus, the sample sizes for both conditions (control and treatment) are identical, and the volume distribution for all nuclei at the time of inhibitor addition are identical for both conditions. Resulting in 43 and 67 nuclear trajectories for the two aphidicolin:control pairs, and 74 nuclear trajectories for the control:control pair.

##### Nuclear import inhibitor analysis dataset

The two colony movies in the Nuclear import inhibitor FOV dataset were processed using the same steps as the baseline dataset (Vision Transformer-based deep-learning segmentation model, feature extraction and automatic tracking of single nuclei). Next the automated trajectory quality control filter was applied to remove problematic segmentations (such as those touching the edge of the FOV) or short trajectories. To compare nuclear volume trajectories from before and after importazole addition from the same colony, nuclear trajectory selection process was performed in the following way. Nuclear trajectories were required to be at least 45 minutes long, to begin with classifier-predicted lamin shell formation event, and have an initial volume less than 450 µm^3^. The “before” trajectories were required to begin at least 90 minutes prior to importazole addition so that the full process of rapid expansion could be observed prior to truncating the trajectory at the time of inhibitor addition. Because long exposure to importazole can disrupt mitotic spindle formation^44^ only trajectories that began within the 1 hr window after importazole addition were included in the “after” trajectories. The “after” trajectories were also truncated 2 hours after importazole addition (30 minutes before the 150 minute mark where 25% of the importazole treated colony is dead or dying). Lastly, to be certain that only nuclei exiting mitosis were included, a manual curation was performed to remove 3 nuclei that did not exhibit volume growth indicative of expansion, resulting in 64 nuclear trajectories “before” importazole addition and 24 “after” for analysis. The same protocol was applied to the control colony resulting in 38 nuclear trajectories “before” mixing and 39 trajectories “after.”

##### Fixed control analysis dataset

The Vision Transformer-based deep-learning segmentation model was applied to generate instance segmentation of nuclei. Next, we performed feature extraction and nuclear tracking on the segmented images resulting in 248 nuclei that were tracked for all 20 frames (Supplemental Fig. S17A). We used these trajectories to assess how the noise pattern in the raw data and cells spatial displacement impact the resultant segmentations and therefore the measured nuclear features. To calculate the percent and absolute error in the measured nuclear features we assumed that the median value for a given nuclear trajectory is representative of its true value and compared each of the 20 pseudo timepoints measurements to the median. The segmentation model predictions were very consistent for the same nucleus over time– the average percent error for nuclear volume was only 0.64% (5.54 µm^3^), and the maximum percent error for volume was only 4.64% (or 31.94 µm^3^, Supplemental Fig. S17).

#### Data Analysis

##### Calculation of nuclear volume, height, and XY aspect ratio

The nuclear volume was calculated by summing the voxels present inside the nuclear volume segmentation. The nuclear height was calculated as the distance between the lowest and highest pixels of the nucleus segmentation in the Z-dimension. The XY aspect ratio was calculated as the ratio of length to the width of the nuclear segmentation. The length is defined as the longest axis of the nuclear segmentation in the XY-plane. The width is defined as the length of nuclear segmentation in the plane perpendicular to the longest axis.

##### Alignment of baseline colonies into an “aligned colony time”

The change in mean nuclear height over time was used to align colonies in their development. The time of best alignment between two colonies was found by minimizing the mean squared difference between their mean height trajectories. First, the medium colony is time-shifted to align with the small colony. Then the large colony is time-shifted to align to the medium colony. The result of this alignment is shown in Fig. 3B.

##### “Density” feature measured from neighbor distances used for quantifying crowding

We first computed a Voronoi tessellation graph from the X and Y positions of nuclear centroids within the colony and defined nuclei to be neighbors if they were “adjacent,” or shared an edge in this tessellation graph. We determined the set of neighbors for each nucleus at each timepoint, and computed the average of centroid-to-centroid distances between each nucleus and all its neighbors. The inverse square of the average distance results in a per area metric that approximates the local density (Fig. S3B). In the single nucleus manifest, this feature was given the short name “density” and serves as a simple metric for quantifying how densely packed nuclei are in the local environment of any given single nucleus.

##### Parameter-free model for exploring the link between nuclear height and local density

To explore the relationship between nuclear height and local density while controlling nuclear volume, we considered a model of cells as packed right hexagonal cylinders. We first compute cell volume from nuclear volume using a nuclear to cytoplasmic ratio of 0.28, as measured for this cell type previously.^2^ To account for variability in volume, we sample nuclei that have the same sample volume (630 μm^3^). We then modeled each cell as a hexagonal prism with *Volume* = 3*ash* where *a* is the apothem length, *s* is the side length, and *h* is the height of the prism. The radius was calculated as 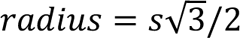. We then plot this model cell radius as a function of nuclear height values interpolated within an expected range, as a model of cell crowding relative to nuclear height. For comparison to the real data, we then approximated the cell radii directly from single nucleus segmentations, estimating the radius of each cell to be half the mean distance between the nucleus and the centroids of its nearest neighbors. We compared the radius of this hexagonal prism toy model to a gaussian weighted moving average of the approximate cell radii (Fig. 3D).

##### Calculation of the normalized distance from colony center

To quantify and compare the positions of nuclei within colonies, which are naturally asymmetric, we computed the “normalized distance from colony center” giving a relative radial position of the nucleus between the colony center and edge. This metric ranges from 0 at the colony center to 1 at the colony edge (or FOV edge if the colony is larger). To create this metric, we used the Voronoi tessellation graph adjacency definition described in the section “*Density feature measured from neighbor distances used for quantifying crowding.”* Starting by defining nuclei on the colony boundary to have a depth of one, we used the adjacency definition to assign a colony depth to all nuclei in the field of view (FOV). We then normalized the colony depth metric such that each frame consisted of nuclei with a “normalized distance from colony center” ranging from 0 at the colony center to 1 at the outer boundary of the colony, according to the formula *d_N_* = (*d_max_* – *d*)/(*d_max_* - *d_min_*), where *d* is the calculated colony depth, *d_N_* is the normalized depth, *d_min_* and *d_max_*. are the minimum and maximum colony depths for a given frame, respectively (Fig. 3C, top left panel). This normalization allows us to compare positions of nuclei within asymmetric colonies of varying sizes due to growth from frame to frame or size differences across colony datasets. Once the colony edge grows beyond the FOV, this metric measures from the center (0) to the boundary of the imaged region (1). The colony edge is no longer within the FOV at hour 40, 32, and 18 for the Small, Medium, and Large colony respectively.

##### Radial slope of colony height

For each frame in a colony timelapse, we fit a linear regression between the nuclear heights and their radial locations within the colony (“Calculation of the normalized distance from colony center” above) to obtain the slope of the best-fit line. A positive slope indicates that nuclei with higher values of the “normalized distance from colony center” (closer to the colony boundary) are taller, and vice versa for a negative slope (Fig. 3C, top right). To quantify the strength of the relationship between the height and the normalized distance from colony center, we used the Pearson correlation coefficient to identify monotonic trends without assuming linearity (Supplemental Fig. S3A). We then created a scatter plot of the slope of the best-fit line against the aligned colony time to visualize the variation in the spatial patterning of nuclear height over the course of colony development (Fig. 3C, bottom). The size of the plot markers indicates the strength of the relationship between the height and the normalized distance from colony center, with larger markers indicating a stronger relationship.

##### Normalized interphase time

The normalized time is a measure of time for a given full-interphase trajectory that ranges from 0 to 1, where 0 represents the time of nuclear lamin shell formation and 1 represents the time of nuclear lamin shell breakdown. Normalized time is found by performing the min-max normalization by subtracting the starting time (formation) and dividing by the difference between the ending time (breakdown) and the starting time for each timepoint in the trajectory.

##### Calculation of nuclear volume scaling with time

To quantitatively characterize the shape of nuclear volume trajectories, we fitted each volume trajectory from transition to breakdown to a power law scaling with time *V*(*t*) = *V_start_* + *rt^a^* , where *V_start_* gives the fitted trajectory volume at the start of growth, *r* is a rate whose units depend on α, and α gives the scaling of the volume with time. As α = 1 would give linearity, the value of this time scaling relative to one describes the super- or sub-linear behavior of any trajectory.

##### Calculation of transient growth rate

The transient growth rate at any time=t (Fig. 5) is the change in volume divided by change in time for a time window centered at time t (Fig. 5E). It is calculated in the following way:

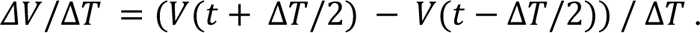

If no value is present at a given time point, *ΔV*/*ΔT* is returned as NaN. To ensure that the transient growth rate is only analyzed during the growth phase and does not capture expansion or lamin shell breakdown dynamics, only timepoints more than two hours after lamin shell formation and at least 30 minutes before lamin shell breakdown were used in the calculation of *ΔV*/*ΔT* unless otherwise noted. To determine the transient growth rate of a nucleus’ neighbors, the transient growth rates of all nuclei within a 90 µm radius surrounding the specific nucleus were averaged.

##### Nuclear surface area

We found that the growth dynamics of nuclear surface area as measured directly from the segmentation mesh (see “Single nucleus feature extraction”) generally track as expected with the nuclear volume growth dynamics. For example, the nuclear surface area increases by, on average, 1.7-fold during growth. This is unsurprising, given that the nuclear volume on average doubles and the nucleus is roughly an oblate spheroid. If the nucleus was a sphere, a doubling of volume would result in a 1.6-fold surface area increase; for an oblate spheroid we would expect it to be slightly higher, consistent with the results of this study. The segmentation mesh used to calculate the surface area does not have high enough resolution to capture fine details of shape, therefore this surface area measurement cannot be analyzed as though it biologically represents the nuclear membrane surface area.

##### Bootstrapping confidence intervals for correlations

To get the 90% confidence intervals for the correlation between two features, we bootstrap resample the data 500 times with replacement. In each iteration, a Pearson correlation is calculated between the specified features. We then calculate the 5th and 95th percentiles of all the resampled correlations.

##### Lineage analysis

We used the lineage-annotated analysis dataset with 316 lineage trees to find nuclear pairs that were traced for their full-interphase resulting in 262 sister pairs, 362 mother-daughter pairs and 191 cousin pairs. Due to the length of the hiPS cell cycle durations (∼15 hours), the length of the timelapse movies (48 hours), and the expansion of the colony outside the imaging field of view it was rare to capture 3 generations of full-interphase nuclear trajectories (N=34 grandmother-daughter pairs). Because the relationships decay for cousin pairs (data not shown) and there were very few grand mother-daughter pairs, we focus on the relationships between mother-daughter and sister pairs. Growth features were compared across these two types of relationships; the correlation values given are Pearson correlations. Error bars are the 90% lower and upper bounds of the bootstrap confidence interval. To control for effects of the colony environment on sister correlations, pairwise comparisons of unrelated nuclei born at the same time (less than ten minutes apart, the time difference in which sister frame formation was detected, see Supplemental Fig. 12B) and the same place (less than 60 μm apart which is the maximum distance sister nuclei move after 1 hour of being born, see Supplemental Fig. S12B) were made. To control for the environmental effects in the mother-daughter correlations, we compared growth features of unrelated pairs of nuclei, selected such that one of the nuclei is born within 60 minutes and 60 μm of the division time and location of the other nucleus. To control for the inheritance of size, control sister pairs were additionally constrained to have less than 80 µm^3^ difference in starting volume. To control for the volume relationship between mother and daughter pairs, control daughter’s starting volume was required to be within 60 µm^3^ of half the control mother’s ending volume.

We categorized sister pairs into classes of asymmetric and symmetrical dividing nuclei. Sisters were considered symmetric when their starting volumes differed by less than 30 µm^3^. Between 30 and 50 µm^3^, differences in starting volumes could be attributed to actual volume differences or differences in the time at which transition was calculated to occur therefore, nuclei that fall in this range were not used in this analysis. Sister nuclei with differences in volume at transition greater than 50 µm^3^ were considered asymmetric (Supplemental Fig. S11B).

#### Analysis of aphidicolin timelapse experiments

To determine how aphidicolin affected growth, we determined the “change in volume” for each nucleus. To do this, a nucleus volume trajectory is normalized by subtracting its volumes at each timepoint by its own average volume over a 15 minute window immediately preceding drug addition. We then plotted the mean and 90% interpercentile ranges for aphidicolin-treated and control trajectories (Supplemental Fig. S7 and S13).

#### Analysis of importazole timelapse experiments

To perform this analysis, the “before” and “after” importazole addition trajectories were aligned at the time of lamin B1 shell formation and their means and 90% interpercentile ranges were plotted (Supplemental Fig. S5). We reported the reduction in nuclear volume for importazole-treated cells at the end of expansion using the ratio of the average “after” and “before” nuclear volumes.

### Data Visualization

#### Colorized segmentation snapshots and supplemental movies

We used the Timelapse Feature Explorer (see Additional Resources), to visualize the spatiotemporal variation of nuclear features across the colony. The resulting colorized maximum projections of 3D nuclear segmentations were exported movies (Movies S9-11) and single-time point field of view snapshots (Fig. 2 and 3A). In figures where the same feature was colorized in snapshots from multiple timepoints, or in multiple colonies, the colormap was set to be consistent across those snapshots, with the colormap range set to demonstrate the variation across the population. Nuclei colored in gray were filtered out due to several criteria: the nuclear segmentation was cut off by the edge of the field-of-view, the nuclear segmentation was marked as an outlier via automated outlier detection, the feature could not be calculated for that nucleus at that time point.

### ADDITIONAL RESOURCES

#### Timelapse Feature Explorer

Timelapse Feature Explorer is a novel web application for interactive visualization and analysis of segmented, time-series microscopy data. The viewer is open-source, with source code available at https://github.com/allen-cell-animated/timelapse-colorizer. To interactively view datasets for analyzing the baseline colonies in Timelapse Feature Explorer, visit: https://timelapse.allencell.org. To create the input data for these interactive visualizations, maximum projections of 3D nuclear segmentations from the baseline colonies FOV timelapse dataset were colormapped based on quantitative features available for the nuclei in each analysis dataset (the baseline colonies, full-interphase, and lineage-annotated analysis datasets). A fourth “exploratory analysis dataset,” which includes outliers, apoptotic nuclei and additional features, is also provided. The formatted input datasets resulting from combining image-based and quantitative data for input into the Timelapse Feature Explorer are provided on Quilt (see Data and code availability), along with documentation of all explorable features. The script to perform this data formatting, along with the scripts to calculate the features available in the Timelapse Feature Explorer, are available at https://github.com/AllenCell/nuc-morph-analysis.

While it was originally developed to view the data presented here, the viewer is broadly useful for visualizing segmented, time-series data. To process and view your own data in Timelapse Feature Explorer, see the code packages and documentation available at https://github.com/allen-cell-animated/colorizer-data.

#### Volume viewer

The web-based volume viewer at allencell.org was enhanced to support loading the FOV datasets associated with the paper as OME-Zarr time series. The viewer can load multiresolution 5D (TCZYX) OME-Zarr data. It is optimized for speed and interactivity and so it relies on having precomputed downsampled images as part of the Zarr. When the source data is sufficiently downsampled, the viewer can auto-select the appropriate resolution, and perform interactive time series playback of 3D volumes or single Z-slices. The viewer can also combine images with matching dimensions from separate URLs. In this way raw microscope images can be distributed separately from segmentation images, but while still brought together easily in the viewer. Timelapses of the baseline colonies are available for viewing in the browser without download of the data at http://volumeviewer.allencell.org/.

#### AGAVE

Renderings provided from AGAVE software^63^ use directed lighting to enhance appearance of 3D shapes. An orthographic camera projection is chosen to retain relative sizes across the rendered colony image. We used a prerelease version of the software to apply color labels to the segmented objects. For more info about AGAVE, visit: https://www.allencell.org/pathtrace-rendering.html.

### The Allen Cell Collection

The hiPSC Single-Cell Image Dataset, protocols, the Allen Cell Discussion Forum and additional information can be found here: https://www.allencell.org/.

---

| REAGENT or RESOURCE | SOURCE | IDENTIFIER |
| --- | --- | --- |
| Chemicals |  |  |
| mTeSR™1 Medium | STEMCELL Technologies | Cat# 85850 |
| mTeSR™1 Medium Without Phenol Red | STEMCELL Technologies | Cat# 05876 |
| Penicillin-Streptomycin | Thermo Fisher Scientific | Cat# 15070063 |
| Matrigel Growth Factor Reduced (GFR) Basement Membrane Matrix, Phenol Red-free, LDEV-free | Corning | Cat# 356231 |
| DMEM/F12 (1:1) 1X | Thermo Fischer Scientific | Cat# 11039021 |
| Gibco DPBS (1X) | Thermo Fischer Scientific | Cat# 14190 |
| Gibco StemPro™ Accutase™ Cell Dissociation Reagent | Thermo Fischer Scientific | Cat# A1110501 |
| Y-27632 ROCK inhibitor | STEMCELL Technologies | Cat# 72308 |
| Importazole | Sigma-Aldrich | Cat# 401105 |
| Aphidicolin | Sigma-Aldrich | Cat# 178273 |
| Puromycin | ThermoFisher | Cat# A1113803 |
| Rapamycin | Selleckchem | Cat# S1039 |
| 2-aminopurine | Sigma Aldrich | Cat# A3509 |
| Click-iT™ EdU Pacific Blue™ Flow Cytometry Assay Kit | ThermoFisher Scientific | Cat# C10418 |
| Paraformaldehyde | Electron Microscopy Sciences | Cat# 15710 |
| DNA dye, Spy650 | Spirochrome | Cat# SC501 |
| Antibodies |  |  |
| Mouse-anti-PCNA | Cell Signaling | Cat# 2586S |
| Goat anti-mouse AlexaFluor 568 | ThermoFisher Scientific | Cat# A-11031 |
| Experimental models: Cell lines |  |  |
| Human induced pluripotent stem cell: AICS-0013 cl 210, lamin B1 | Available through Coriell | RRID:CVCL_IR32 |
| Human induced pluripotent stem cell: AICS-0088 cl 83, proliferating cell nuclear antigen | Available through Coriell | RRID:CVCL_A8RT |
| Deposited data |  |  |
| WTC-11 hiPSC FOV-nuclei timelapse dataset V1 | This paper | https://open.quiltdata.com/b/allencell/tree/aics/nuc-morph-dataset/hipsc_fov_nuclei_timelapse_dataset/ |
| hiPSC FOV-nuclei timelapse data for pretraining Vision Transformer | This paper | https://open.quiltdata.com/b/allencell/tree/aics/nuc-morph-dataset/hipsc_fov_nuclei_timelapse_dataset/hipsc_fov_nuclei_timelapse_data_for_pretraining_vision_transformer/ |
| hiPSC FOV-nuclei timelapse data used for analysis | This paper | https://open.quiltdata.com/b/allencell/tree/aics/nuc-morph-dataset/hipsc_fov_nuclei_timelapse_dataset/hipsc_fov_nuclei_timelapse_data_used_for_analysis |
| Baseline colonies FOV timelapse dataset | This paper | https://open.quiltdata.com/b/allencell/tree/aics/nuc-morph-dataset/hipsc_fov_nuclei_timelapse_dataset/hipsc_fov_nuclei_timelapse_data_used_for_analysis/baseline_colonies_fov_timelapse_dataset/ |
| DNA replication inhibitor FOV timelapse dataset | This paper | https://open.quiltdata.com/b/allencell/tree/aics/nuc-morph-dataset/hipsc_fov_nuclei_timelapse_dataset/hipsc_fov_nuclei_timelapse_data_used_for_analysis/dna_replication_inhibitor_fov_timelapse_dataset/ |
| Nuclear import inhibitor FOV timelapse dataset | This paper | https://open.quiltdata.com/b/allencell/tree/aics/nuc-morph-dataset/hipsc_fov_nuclei_timelapse_dataset/hipsc_fov_nuclei_timelapse_data_used_for_analysis/nuclear_import_inhibitor_fov_timelapse_dataset/ |
| Feeding control FOV timelapse dataset: | This paper | https://open.quiltdata.com/b/allencell/tree/aics/nuc-morph-dataset/hipsc_fov_nuclei_timelapse_dataset/hipsc_fov_nuclei_timelapse_data_used_for_analysis/feeding_control_fov_timelapse_dataset/ |
| Fixed control FOV timelapse dataset | This paper | https://open.quiltdata.com/b/allencell/tree/aics/nuc-morph-dataset/hipsc_fov_nuclei_timelapse_dataset/hipsc_fov_nuclei_timelapse_data_used_for_analysis/fixed_control_fov_timelapse_dataset/ |
| PCNA + DNA dye FOV timelapse dataset | This paper | https://open.quiltdata.com/b/allencell/tree/aics/nuc-morph-dataset/hipsc_fov_nuclei_timelapse_dataset/hipsc_fov_nuclei_timelapse_data_used_for_analysis/pcna_dna_dye_fov_timelapse_dataset/ |
| hiPSC single-nuclei timelapse analysis datasets | This paper | https://open.quiltdata.com/b/allencell/tree/aics/nuc-morph-dataset/hipsc_single_nuclei_timelapse_analysis_datasets/ |
| hiPSC nuclei image datasets for training deep-learning models | This paper | https://open.quiltdata.com/b/allencell/tree/aics/nuc-morph-dataset/hipsc_nuclei_image_datasets_for_training_deep_learning_models/ |
| Segmentation decoder training FOV dataset | This paper | https://open.quiltdata.com/b/allencell/tree/aics/nuc-morph-dataset/hipsc_nuclei_image_datasets_for_training_deep_learning_models/segmentation_decoder_training_fov_dataset/ |
| Interphase detector single-nuclei image dataset | This paper | https://open.quiltdata.com/b/allencell/tree/aics/nuc-morph-dataset/hipsc_nuclei_image_datasets_for_training_deep_learning_models/interphase_detector_single_nuclei_image_dataset/ |
| Timelapse Feature Explorer datasets | This paper | https://open.quiltdata.com/b/allencell/tree/aics/nuc-morph-dataset/timelapse_feature_explorer_datasets/ |
| Software and algorithms |  |  |
| Zen 2.3, 2.6 software (blue edition) | ZEISS |  |
| Original/source data for figures in the paper | This paper | https://github.com/AllenCell/nuc-morph-analysis |
| The Vision Transformers are implemented using CytoDL, a Python package that we developed for configurable 2D and 3D image-to-image deep learning transformations and representation learning | N/A | https://github.com/AllenCellModeling/cyto-dl |
| Code to perform nuclear instance segmentation, formation and breakdown classification, and extraction of quantitative shape features | This paper | https://github.com/AllenCell/morflowgenesis |
| Configuration files to execute image processing workflows | This paper | https://open.quiltdata.com/b/allencell/tree/aics/nuc-morph-dataset/supplemental_files/ |
| Code used to track single nuclei over timelapse movie | This paper | https://github.com/AllenCell/aics-timelapse-tracking |
| Code used to perform morphing, compute Shape Modes | Viana et al., 2023 | https://github.com/AllenCell/cvapipe_analysis |
| Code for custom Paraview plugin used to support manual curation of nuclear trajectories | This paper | https://github.com/AllenCell/aics-track-curator |
| Code used to prepare data for the Timelapse Feature Explorer | This paper | https://github.com/allen-cell-animated/colorizer-data |
| Code used to create the Timelapse Feature Explorer | This paper | https://github.com/allen-cell-animated/nucmorph-colorizer. |
| Timelapse Feature Explorer | This paper | https://timelapse.allencell.org. |
| Paraview | Ayachit 2015 | https://www.paraview.org/ |
| AGAVE | Toloudis 2024 | https://allencell.org/pathtrace-rendering.html. |
| Source code for volume viewer front end web application | N/A | https://github.com/allen-cell-animated/website-3d-cell-viewer |
| Volume viewer volume loading and rendering component | N/A | https://github.com/allen-cell-animated/volume-viewer |
| Volume viewer | N/A | https://volumeviewer.allencell.org |
| Other |  |  |
| Falcon 60 mm TC-treated Cell Culture Dish | Corning | Cat# 353002 |
| Falcon 6-well Clear Flat Bottom TC-treated Multiwell Cell Culture Plate | Corning | Cat# 353046 |
| 96-well glass bottom plate with high performance #1.5 grade cover glass | Cellvis | Cat# P96-1.5H-N |
| 24-well glass bottom plate with high performance #1.5 grade cover glass | Cellvis | Cat# P24-1.5H-N |

