## Supplemental Information for "Colony context and size-dependent compensation mechanisms give rise to variations in nuclear growth trajectories"

**Supplemental information in this document includes:**
Figures S1-S17
Captions for Movies S1-S11

**Other Supplemental Materials for this manuscript include:**
Movies S1-S11

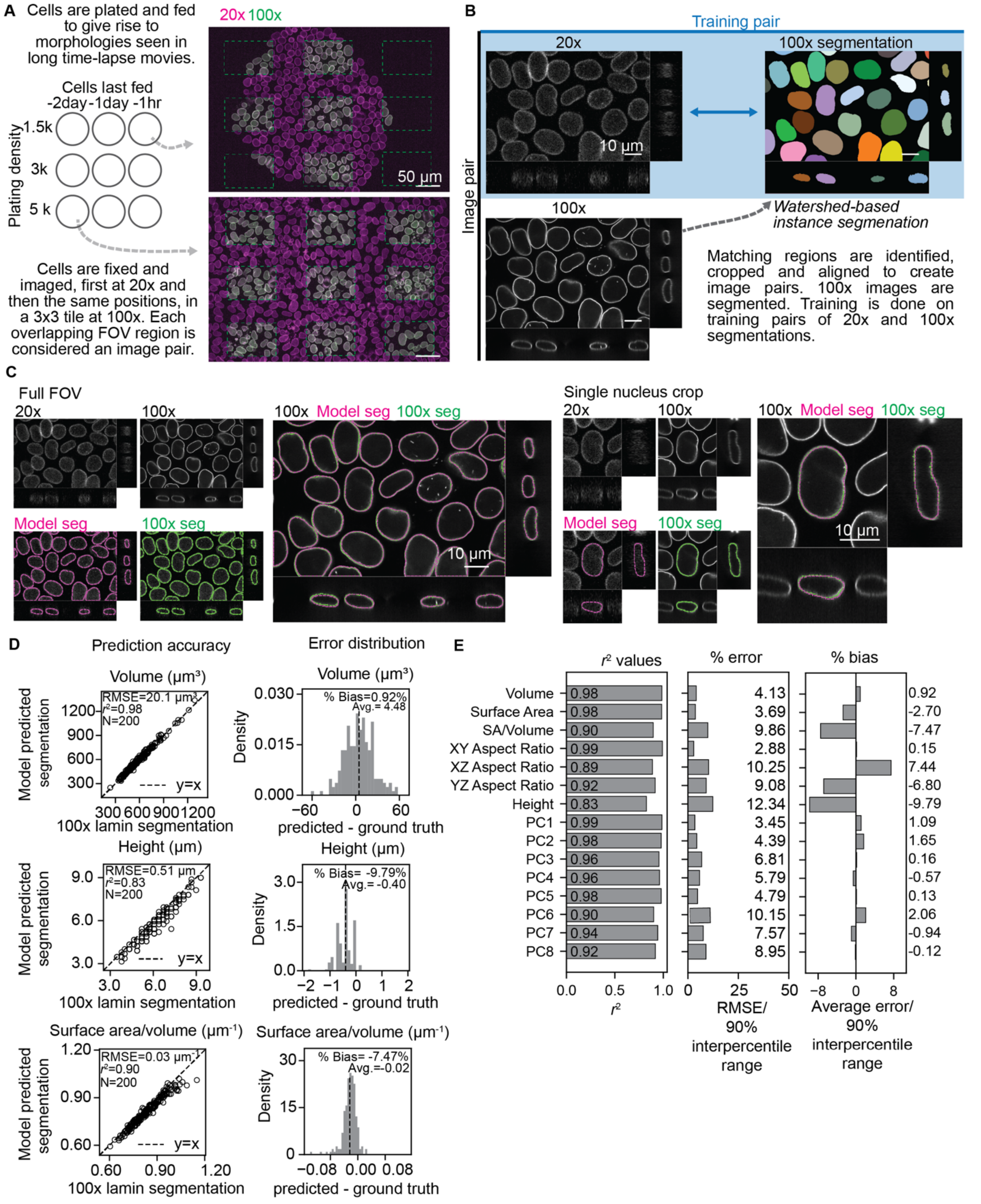


**Figure S1. Development and quantitative validation of a Vision Transformer-based deep-learning segmentation model. A.** Diagram illustrating cell plating and imaging strategy for acquisition of segmentation training data (Methods). **Left:** hiPS cells were plated in a matrix of conditions, varying both the feeding time and plating density to account for the variations in nuclear morphologies arising in a long timelapse acquisition. This includes images of colonies that are initially small and recently fed (top right image) and images of colonies grown for two days, increasing in size and remaining unfed (bottom right image). **Right:** Every colony was imaged in 3D at both 20x and 100x magnification (colored magenta and boxed in green, respectively) generating 9 matched image pairs. Images from two example colonies are shown as top views via maximum intensity projections. **B.** These matched pairs of images are used to generate the segmentation training data for the Vision Transformer-based deep-learning segmentation model. Watershed-based instance segmentations are generated from the matched 100x images to provide high resolution ground truth data for training (see Methods and Supplemental Fig. S14). For details about model training, including pretraining of the Vision Transformer, see Supplemental Fig. S2 and Methods. **C.** To qualitatively validate the segmentation model accuracy, the trained model was applied to 20x images from test set data (i.e. not used for training) and model-produced instance segmentations ("Model seg", magenta) were overlayed with corresponding 100x image derived ground truth instance segmentations ("100x seg", green). The consistent overlap shows the high level of segmentation accuracy achieved in the model-produced segmentations. The single nucleus crop images (right) further show the ability of the model to accurately segment the complex curvature of the tops and bottoms of the nuclei despite the very low contrast in these regions in the original 20x data. **D.** To quantitatively validate the segmentation model accuracy, nuclear shape features were quantified from matched single nuclei in the model-produced segmentations and the test set 100x ground truth segmentations. **Left:** The nuclear shape features from the ground truth and predicted segmentations were then plotted and correlated. Scatter plots are for volume, height and surface area to volume ratio (top to bottom). The $r^2$ for the goodness-of-fit to the y=x line is reported. Each point represents a single nucleus. **Right:** To determine if the model-predictions were biased (e.g., systematically predicted smaller or larger than expected volumes), the prediction error for each feature was determined ($error\ =\ predicted - ground\ truth$) and plotted as a histogram. The volumes from model-predicted segmentations (top panel) did not show significant bias; the average error was only 4.48 $\mu m^3$, which corresponds to 1.19% of the range of possible nuclear volumes (quantified here as the 90% interpercentile range). The largest prediction bias was observed for surface area to volume ratio (-9.86%, or -0.02 $\mu m^{-1}$). **E.** A summary of the $r^2$ values, percent error (% error) and percent bias (% bias) for all features (see Methods). Percent error and percent bias were calculated relative to the 90% interpercentile range. The segmentation model's high quantitative accuracy (i.e. the strong correlation and small bias), supports reporting values for shape features using physical units, e.g. units of $\mu m^3$ for volume, in all subsequent analyses.

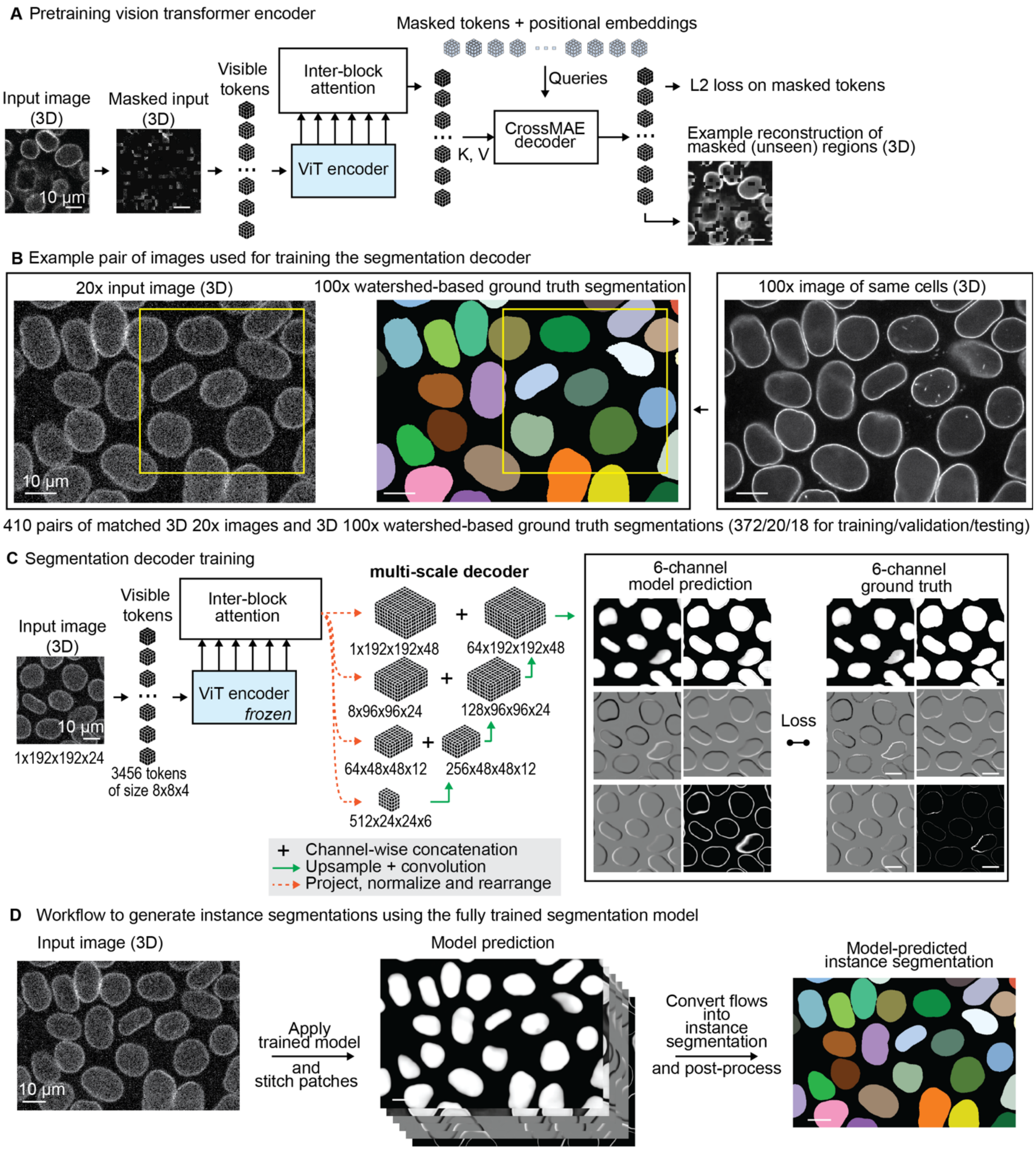


**Figure S2. Illustration of training the Vision Transformer-based deep-learning segmentation model. A.** Illustration of the self-supervised pretraining of the Vision Transformer (ViT) Encoder. We pretrained a ViT Encoder to learn efficient representations of 3D 20x images of lamin B1 using Masked Autoencoding as a pretext task. Briefly, a 192x192x24 (XYZ) voxel patch was extracted from a 20x image and masked at random in 8x8x4 patches, such that only 25% of the image remained visible. The Masked Autoencoder was trained to reconstruct the unseen (masked) 75% of the image. The visible portions of the image were embedded using a single convolution to form visible tokens and passed into the ViT Encoder. A learnable weighted combination of the intermediate representations from each transformer block (Inter-block Attention) were then passed as inputs to the CrossMAE Decoder block along with the masked tokens (which contain no pixel intensity information) and their positional embeddings.[50] Decoded tokens were then linearly projected and rearranged back into an image and the MSE loss was computed by comparing the original pixels within the masked region of the input image with the pixels from decoder-reconstructed masked tokens. The example reconstruction image shows the high quality with which the model reconstructs the 75% percent of the image that was masked. **B.** Example of one training pair of images used for training the segmentation decoder. A watershed-based instance segmentation derived from the 100x image was paired with the 20x image of the same nuclei. **C.** Illustration segmentation decoder training. To adapt the pretrained ViT Encoder for instance segmentation, key changes to the architecture were made: 1) the pretrained ViT Encoder was frozen (the Encoder learning was done during pretraining, see panel B), and 2) the CrossMAE decoder module was swapped out for a multi-scale decoder inspired by UNET.[56] For training, patches of size 1x192x192x24 (XYZ) and 480x480x62 (XYZ) were extracted from the 20x image and 100x instance segmentation, respectively (yellow boxes in **B**). The 20x image was passed through the pretrained ViT encoder. The outputs from the ViT Encoder were then passed into the decoder. Each layer of the decoder received as input a distinct Inter-block Attention output concatenated with the upsampled output from the previous decoder layer. The final output of the decoder is a 6-channel output of size 6x480x480x62. Loss was computed by comparing this 6-channel model prediction image to the 6-channel ground truth image, which is a representation derived from the input ground truth image. For specifics about the generation of each of the 6 channels and the loss functions used for each channel, see Methods. **D.** Illustration of how the trained Vision Transformer-based deep-learning segmentation model was applied to new images to generate instance segmentations. **Left:** A 3D 20x image of lamin B1 mEGFP fluorescence was provided as an input. Overlapping patches of size 192x192x24 were fed into the model sequentially. **Middle**: The resulting 6-channel model predictions were stitched together by overlapping patches by 30x30x4. **Right:** The model predictions were then converted into an instance segmentation

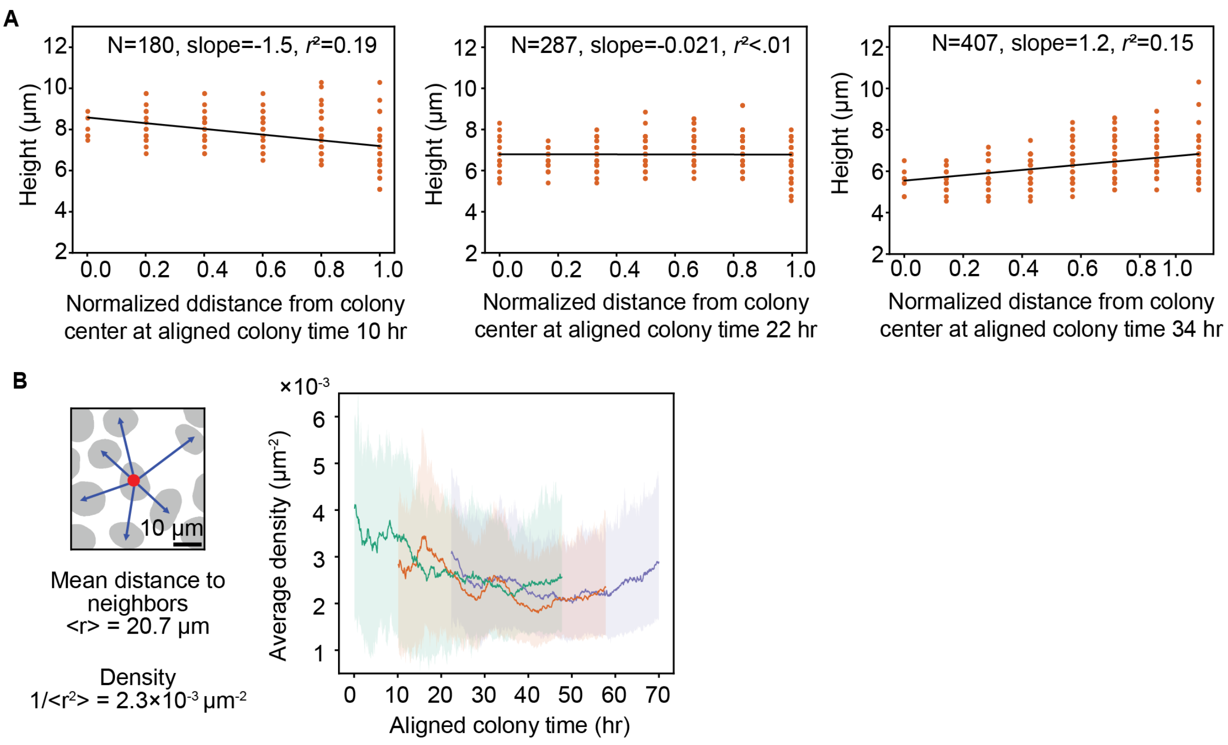


**Figure S3. Colony development and neighborhood density give rise to changes in nuclear height dynamics. A.** The relationship between height and the normalized distance from the colony center is shown for three example timepoints in aligned colony time (10 hours representing early, 22 hours representing middle and 34 hours representing late aligned colony times, from left to right). These examples are all taken from analysis of the Medium colony (standard orange coloring for this colony). The slope of the linear fit and the goodness of fit for all timepoints for all the colonies are reported in Figure 3. **B. Left:** The local density for a given nucleus is approximated by calculating the average of centroid-to-centroid distances between a nucleus and all its neighbors and converting this average distance into a per area metric (Methods). **B. Right:** The local density is averaged for each nuclear trajectory from the baseline colonies analysis dataset and plotted over the aligned colony time at the start of growth for each trajectory. The rolling mean is shown for each colony to guide the eye.

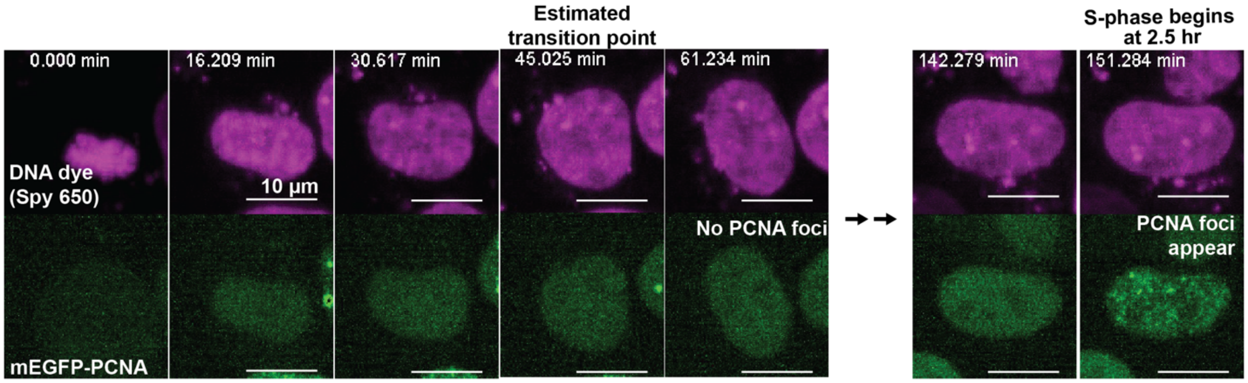


**Figure S4. S-phase onset occurs much later in interphase than the transition point. A.** Image sequence following a single nucleus expressing mEGFP-PCNA stained with a DNA dye (Spy650). The transition point was estimated by identifying the time at which the nucleus (via Spy650 fluorescence, top row) no longer showed rapid expansion in size. The transition point of this nucleus was estimated to occur at ~45 minutes, which is consistent the average transition point timing of 38±12 minutes determined from the nuclear volume trajectories of segmented lamin B1 in the full-interphase analysis dataset. The onset of S-phase is indicated by the appearance of PCNA foci. In this nucleus S-phase begins at ~150 minutes, which is 105 minutes after the estimated transition point. See Methods.

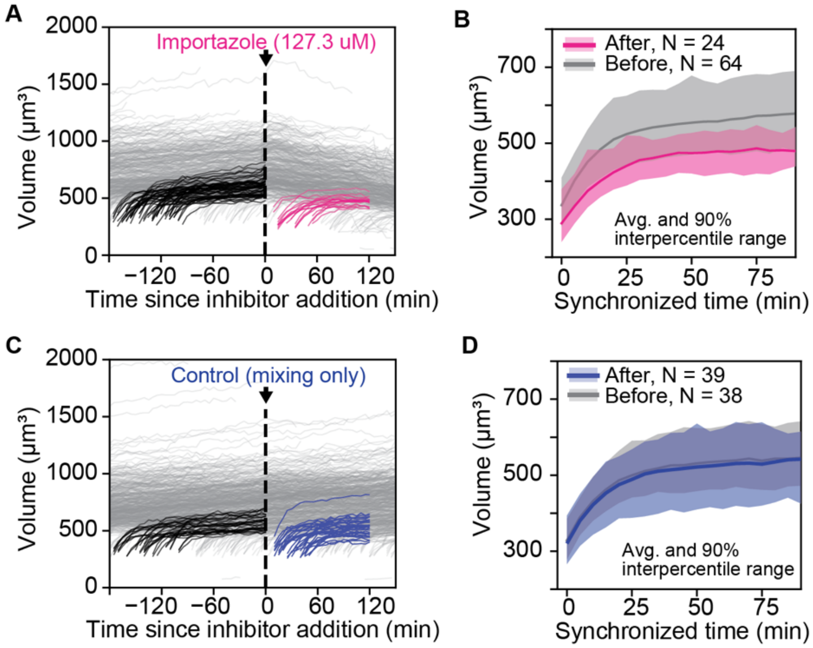


**Figure S5. Cells treated with an inhibitor of nuclear import reach smaller nuclear volumes than untreated cells by the end of the nuclear expansion phase. A.** All nuclear volume trajectories (gray) from a colony of cells treated with importazole, an inhibitor of nuclear import. Cells received a 127.3 µM dose of importazole after 175 minutes of initial imaging. Nuclei born after the addition of importazole were selected for analysis (magenta, "After"), along with nuclear volume trajectories completing the expansion phase prior to addition of importazole as a control (black, "Before"). To account for differences in nuclear growth that can occur from colony to colony, the control and inhibitor-treated nuclear volume trajectories were selected from the same colony. To avoid any effects due to cell death, no nuclei were analyzed once the fraction of cells dying in the importazole-treated colony reached 25%, which occurred 150 minutes after inhibitor addition. **B)** To determine the effect of inhibiting nuclear import, the nuclear volume trajectories from "Before" and "After" addition of importazole were synchronized by the time of lamin shell formation and compared. Plotted is the average nuclear volume (thick line) with the shaded area indicating the 90% interpercentile range. **C and D.** Same as in **A and B** except the nuclei come from a colony receiving no importazole (mixing only control). As expected, the control "After" nuclear volume trajectories have almost indistinguishable behavior to the control "Before" trajectories.

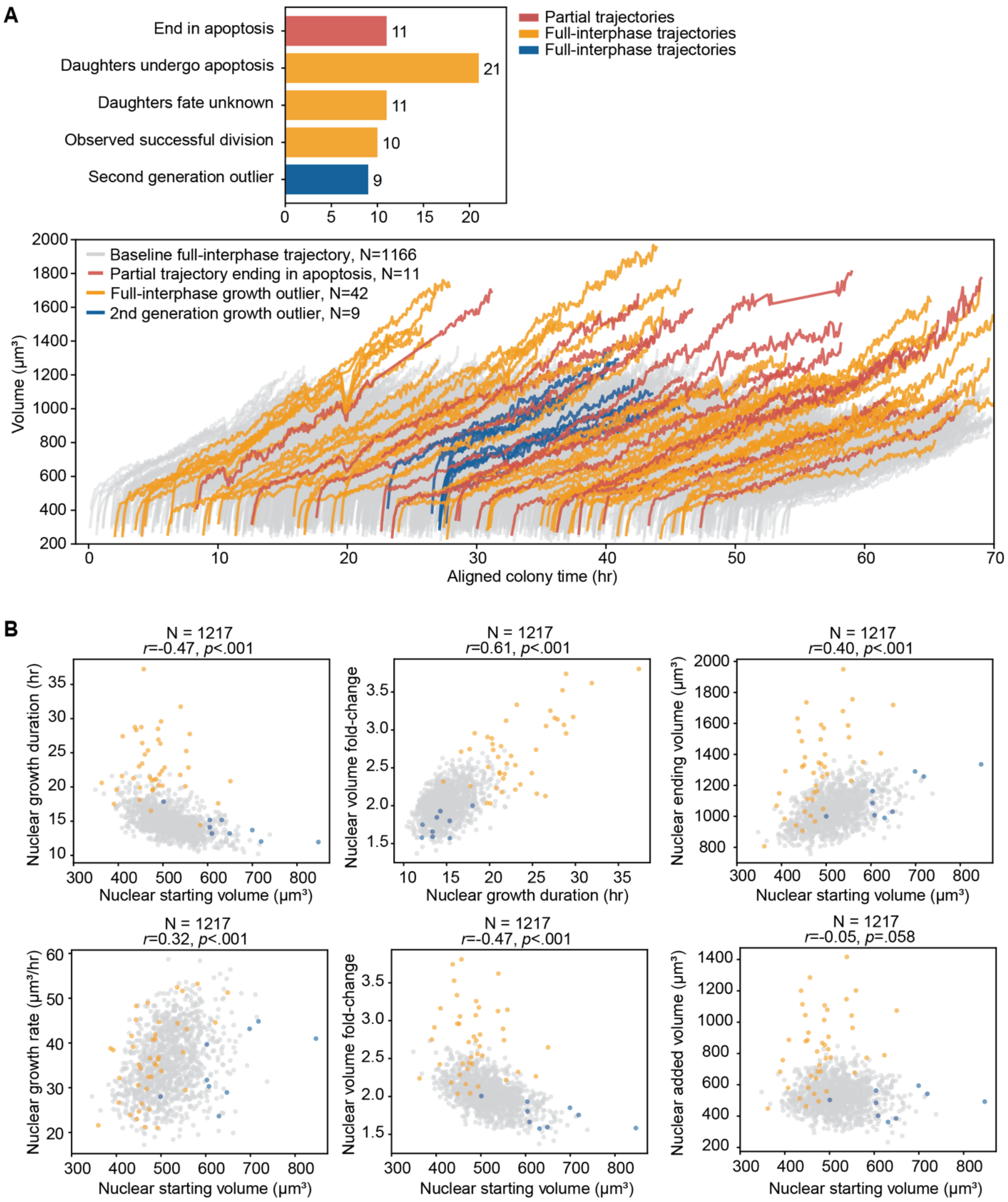


**Figure S6. Identifying and characterizing outlier nuclei with extremely long growth durations. A. Top:** Bar graph showing statistics for nuclei identified to have abnormally long growth trajectories compared to the population and their daughters**.** Of the 53 nuclei identified, 11 end in apoptosis (red) and 42 divide (yellow). Within this population of 42 nuclei, 21 successfully undergo division but have daughters that undergo apoptosis as they exit mitosis or shortly after it; 11 undergo division right as the timelapse movie ends (and so the fate of their daughters was not observed); and the last 10 seem to undergo division successfully despite having abnormally large fold-changes and large ending volumes (similar to the outliers that ended in apoptosis). The nine resulting daughters with full-interphase trajectories (blue) are excluded from the full-interphase analysis dataset, consistent with the exclusion of their mother nuclei. **Bottom:** Nuclear volume trajectories for full-interphase trajectories over aligned colony time with long trajectories ending in apoptosis highlighted in red, growth outliers highlighted in yellow and daughters of growth outliers highlighted in blue. **B.** The relationships previously shown between growth features (nuclear starting volume, growth duration, ending volume, added volume, volume fold-change, and growth rate) for all nuclear trajectories in the full-interphase analysis dataset (gray) with the growth outliers (yellow) and their daughters (blue) included. Volume trajectories with abnormally long durations are often also the outliers in other growth features (nuclear volume fold-change, ending volume, added volume). Daughters of growth outliers who successfully complete (and are tracked throughout) interphase (blue) tend to have large starting volumes; their growth behavior is consistent with that of other nuclei with large starting volumes from the full-interphase analysis dataset (gray). Previously shown in Fig. 6 and Supplemental Fig. S9 do not have significant differences in correlation or interpretation when growth outliers and their daughters are included.

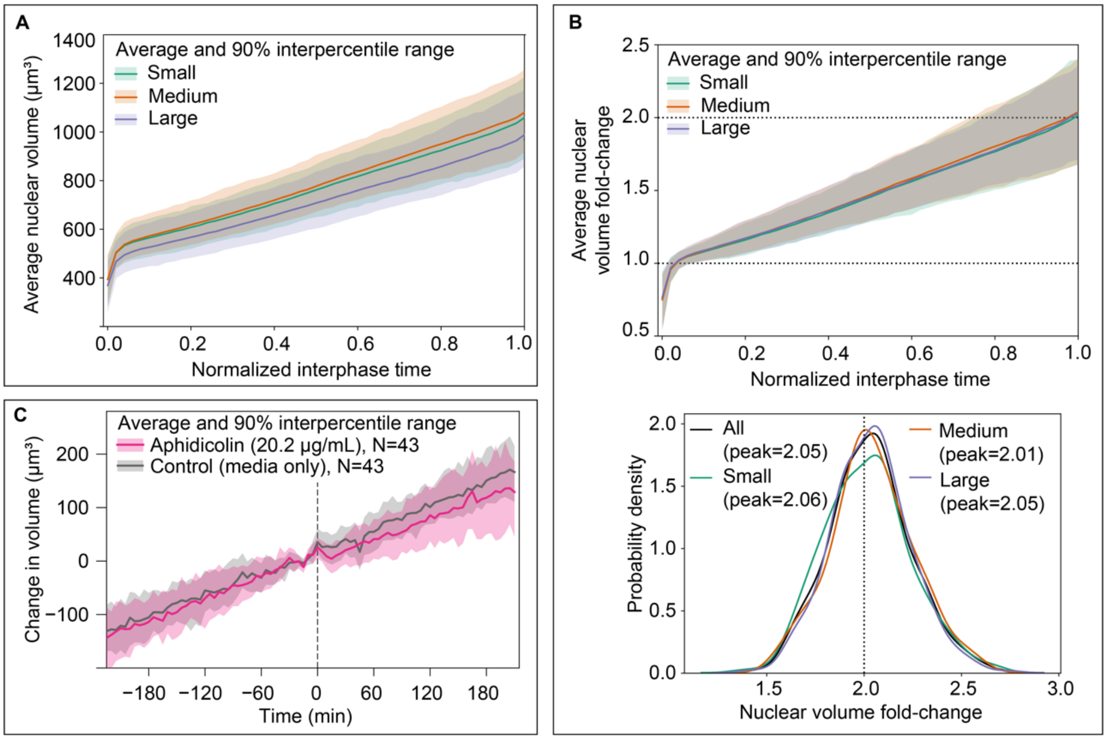


**Figure S7. Characterizing nuclear growth trajectories for the Small, Medium, and Large colonies.** **A.** The per-colony average of all nuclear volume trajectories from full-interphase analysis dataset from each of the Small (N=354), Medium (N=429) and Large (N=383) colonies. The 90% interpercentile range is shaded. Full-interphase nuclear volume trajectories were synchronized to start at a common t=0 (time of shell formation for each individual trajectory) and rescaled to normalized interphase time (see Methods). **B. Top:** The average nuclear volume fold-change and 90% interpercentile range of full-interphase nuclear trajectories from each colony, synchronized by their formation and rescaled to normalized interphase time. Nuclear volume at the start of growth (transition) was normalized to 1 (dashed line at fold-change of 1 for reference). Each colony reaches a population-level average volume that is approximately double from the start to the end growth (dashed line drawn at fold-change of 2 for reference). **Bottom:** The probability density of the nuclear growth volume fold-changes peaks (reaches the fold-change giving the maximum probability density) at just over 2 for each colony (dashed line at fold-change of 2 for reference). The variation around doubling for each colony is also consistent between colonies. **C.** The average change in nuclear volume for nuclei from a colony that received a dose of 20.2 µg/mL aphidicolin (magenta) and control colony receiving only media (gray). The change in volume was determined by subtracting the nuclear volume at the time of inhibitor addition (defined here as t=0) from the entire nuclear volume trajectory for all nuclei. Shaded regions represent the 90% interpercentile range. The single nucleus volume trajectories were selected such that each trajectory from the aphidicolin-treated trajectory was matched with a trajectory from the control colony that had the same volume at the frame before inhibitor addition, thus the sample sizes (N) are identical. This was done to account for potential size dependent effects of the inhibitor (see Methods). Additional aphidicolin-treated colonies and control colonies are plotted in Supplemental Fig. S13. Further, to rule out the possibility that the small effect is due to incomplete inhibition of DNA replication, a control experiment was performed to confirm that aphidicolin treatment rapidly halts the progression of DNA replication in hiPS cells (Supplemental Fig. S13 and Methods).

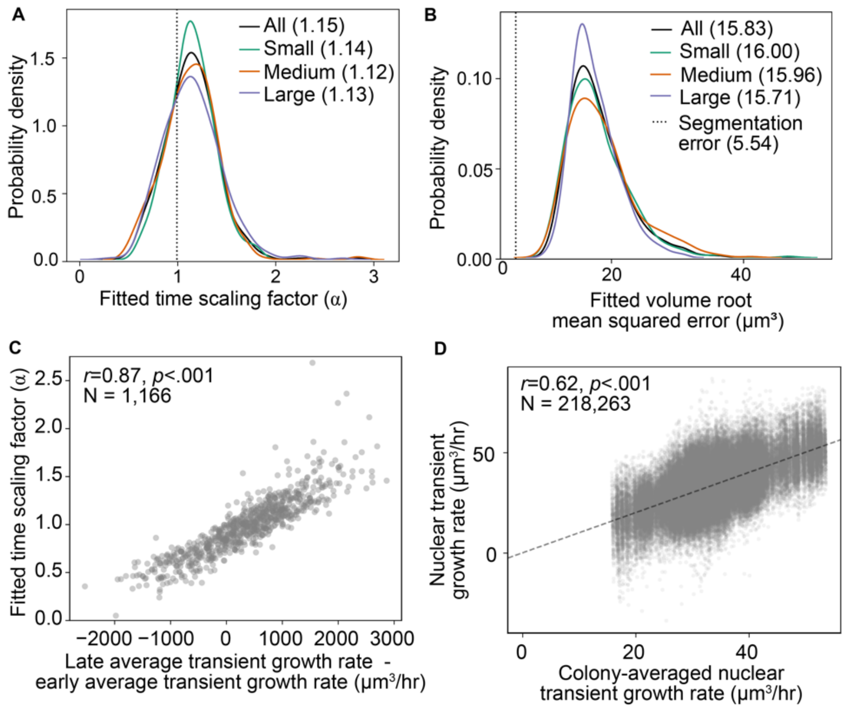


**Figure S8. The fitted time scaling factor can be recapitulated by the transient growth rate, which captures correlated changes in growth rate for individuals and across the colony. A.** The probability density of the fitted time scaling factor has modal value of 1.15 for the set of all trajectories (black); also shown are modal values for each colony individually (green, orange, and purple). Dashed line at fitted time scaling factor of 1 designating linear growth is shown for reference. **B.** The probability density of the fitted volume root mean squared error. The fitted volume root mean squared errors are greater than the mean segmentation error (dashed line at 5.54, Supp. Fig. S17). **C.** The relationship between the fitted time scaling factor and the difference between the average early and late transient growth rates for all nuclei in the full-interphase analysis dataset with Pearson's correlation coefficient (*r*) and p-values (*p*). The average early and late transient growth rates for a given trajectory are the average of all transient growth rates in the first 30% and last 25% of time points, respectively, for that trajectory. **D.** The nuclear transient growth rates for individual nuclei compared to the colony-averaged nuclear transient growth rate at a given timepoint. Dashed unity line shown for reference.

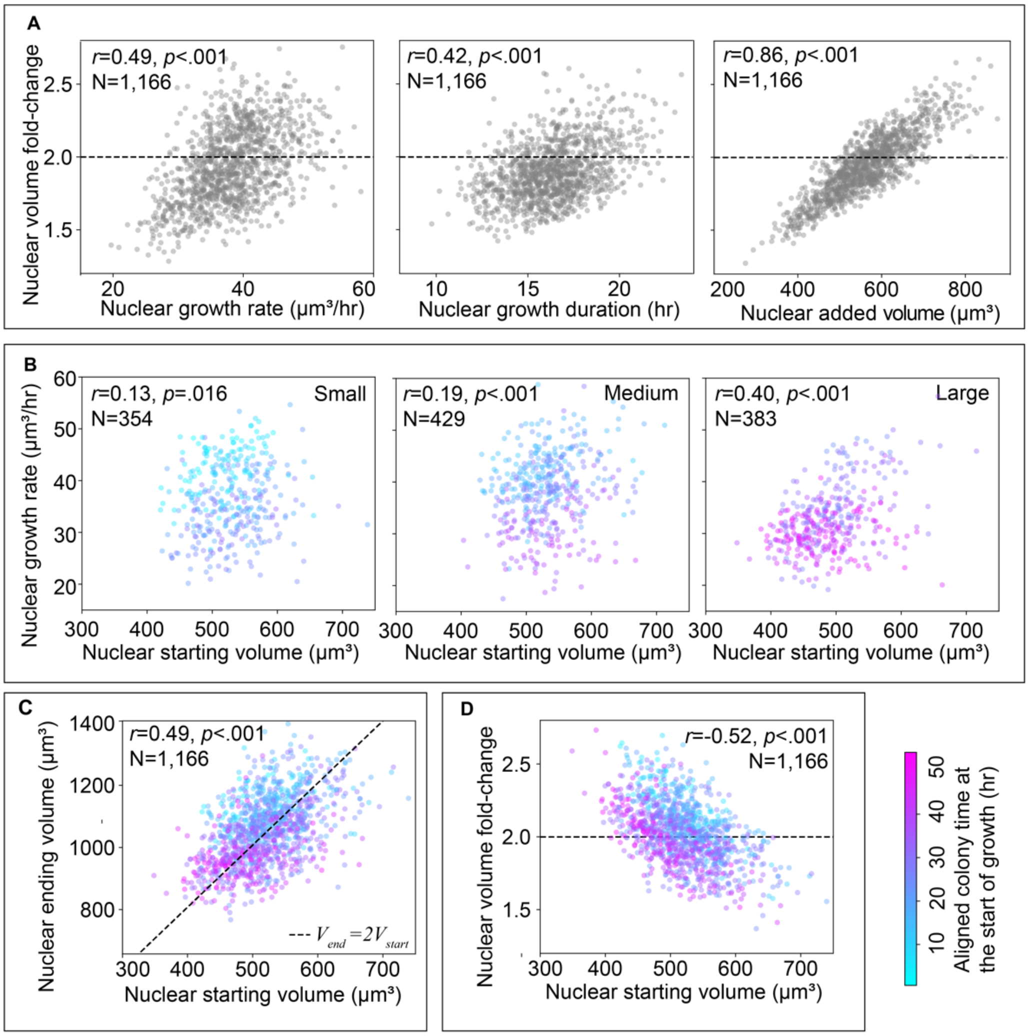


**Figure S9. Nuclear volume fold-change is dependent on various growth features, and nuclear growth rate, ending volume and volume fold-change all decrease slowly over time. A.** The relationships between nuclear volume fold-change and nuclear growth rate (left), growth duration (middle), and added volume (right) for trajectories from all colonies pooled together, with Pearson's correlation coefficient ($r$) and p-values ($p$). **B.** The relationship between nuclear growth rate and nuclear starting volume trajectories from the Small (left), Medium (middle), and Large (right) colonies, colored by aligned colony time at the start of growth. The same colormap is applied to all three plots, and a color bar is provided in bottom right of figure. **C.** The relationship between nuclear ending volume and starting volume for the trajectories from all colonies pooled together, colored by aligned colony time at the start of growth with doubling line (dotted line) for reference. **D.** The relationship between nuclear volume fold-change and starting volume for the full-interphase analysis dataset colored by aligned colony with doubling line (nuclear volume fold-change of 2; dotted line) for reference.

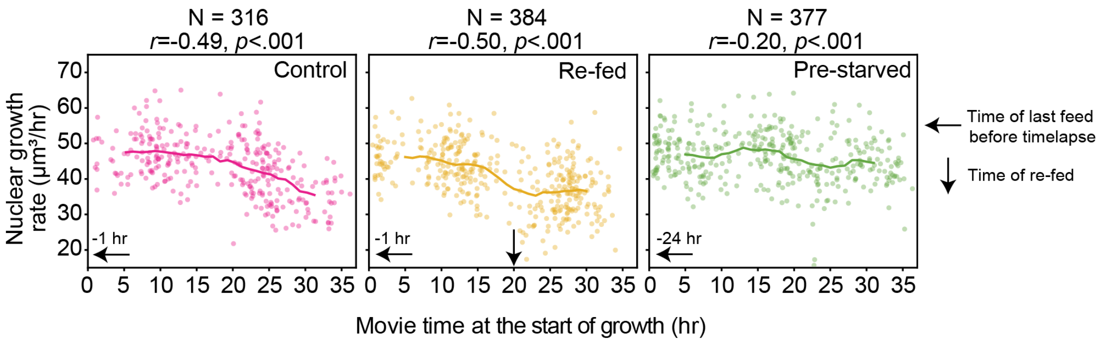


**Figure S10. The decrease in growth rate observed over aligned colony time is not explained by nutrient depletion.** To test the effect of nutrient depletion on the nuclear growth rate, the nuclear growth rates over movie time for nuclei from three colonies of the same size were analyzed. **Left:** The control colony is imaged under the same conditions as the baseline colonies, with cells fed one hour prior to the start of the timelapse. The correlation of nuclear growth rate and time at the start of growth is -0.49, consistent with the observation that the growth rate is decreasing over time in the baseline colonies analysis dataset. **Middle:** The re-fed colony was fed 1 hour prior to the start and halfway through the timelapse. The exchange of fresh media did not change the correlation of nuclear growth rate and time (-0.50). **Right:** The pre-starved condition was last fed 1 day prior to the start of acquisition. This condition has the smallest magnitude correlation between nuclear growth rate and time (-0.20), showing that nutrient depletion is not the cause of this observed decrease over time.

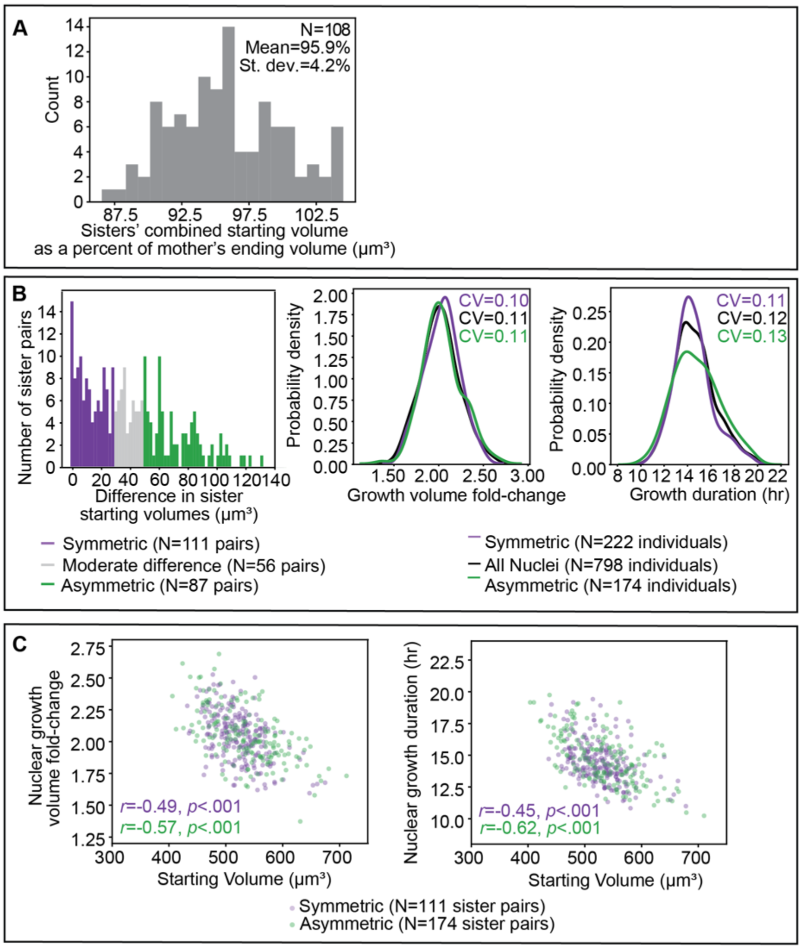


**Figure S11. Starting-size dependent growth trends do not arise from asymmetric division. A.** Distribution of the percent of the mother's nuclear ending volume reached by the sisters' combined nuclear starting volume for the lineage-annotated analysis dataset. The mean and standard deviation are shown. **B. Left:** The distribution of the difference in nuclear starting volumes between sister pairs. In this panel, and in subsequent panels, "symmetric" nuclei have a difference in starting volume less than 30 µm$^3$ (purple) and "asymmetric" nuclei (green) have a nuclear starting volume of greater than 50 µm$^3$ (see Methods). **Middle, Right:** The probability density of the nuclear volume fold-change (middle) and the growth duration (right) for symmetric sisters, asymmetric sisters, and all nuclei from the lineage-annotated analysis dataset with the coefficient of variation (CV) for each. **C.** The relationship of the nuclear volume fold-change (left) and the growth duration (right) with the nuclear starting volume for symmetric and asymmetric sister pairs with Pearson's correlation coefficient (*r*) and p-values (*p*).

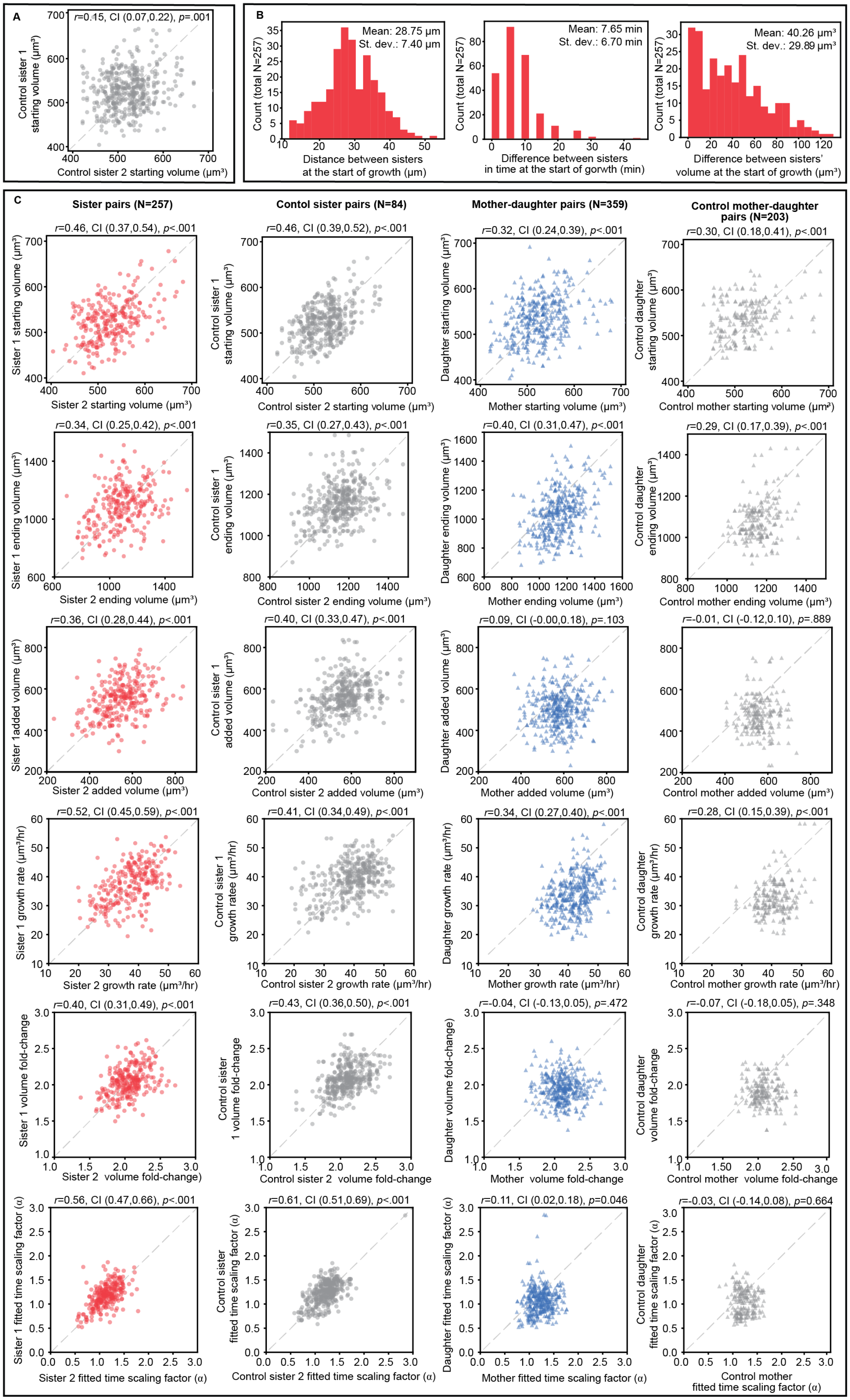
A
r=0.15, CI (0.07,0.22), p=.001
Control sister 1 starting volume (µm³)
Control sister 2 starting volume (µm³)
B
Mean: 28.75 µm
St. dev.: 7.40 µm
Count (total N=257)
Distance between sisters at the start of growth (µm)
Mean: 7.65 min
St. dev.: 6.70 min
Difference between sisters in time at the start of gorwth (min)
Mean: 40.26 µm³
St. dev.: 29.89 µm³
Difference between sisters' volume at the start of growth (µm³)
C
Sister pairs (N=257)
Contol sister pairs (N=84)
Mother-daughter pairs (N=359)
Control mother-daughter pairs (N=203)
r=0.46, CI (0.37,0.54), p<.001
Sister 1 starting volume (µm³)
Sister 2 starting volume (µm³)
r=0.46, CI (0.39,0.52), p<.001
Control sister 1 starting volume (µm³)
Control sister 2 starting volume (µm³)
r=0.32, CI (0.24,0.39), p<.001
Daughter starting volume (µm³)
Mother starting volume (µm³)
r=0.30, CI (0.18,0.41), p<.001
Control daughter starting volume (µm³)
Control mother starting volume (µm³)
r=0.34, CI (0.25,0.42), p<.001
Sister 1 ending volume (µm³)
Sister 2 ending volume (µm³)
r=0.35, CI (0.27,0.43), p<.001
Control sister 1 ending volume (µm³)
Control sister 2 ending volume (µm³)
r=0.40, CI (0.31,0.47), p<.001
Daughter ending volume (µm³)
Mother ending volume (µm³)
r=0.29, CI (0.17,0.39), p<.001
Control daughter ending volume (µm³)
Control mother ending volume (µm³)
r=0.36, CI (0.28,0.44), p<.001
Sister 1added volume (µm³)
Sister 2 added volume (µm³)
r=0.40, CI (0.33,0.47), p<.001
Control sister 1 added volume (µm³)
Control sister 2 added volume (µm³)
r=0.09, CI (-0.00,0.18), p=.103
Daughter added volume (µm³)
Mother added volume (µm³)
r=-0.01, CI (-0.12,0.10), p=.889
Control mother added volume (µm³)
r=0.52, CI (0.45,0.59), p<.001
Sister 1 growth rate (µm³/hr)
Sister 2 growth rate (µm³/hr)
r=0.41, CI (0.34,0.49), p<.001
Control sister 1 growth ratee (µm³/hr)
Control sister 2 growth rate (µm³/hr)
r=0.34, CI (0.27,0.40), p<.001
Daughter growth rate (µm³/hr)
Mother growth rate (µm³/hr)
r=0.28, CI (0.15,0.39), p<.001
Control daughter growth rate (µm³/hr)
Control mother growth rate (µm³/hr)
r=0.40, CI (0.31,0.49), p<.001
Sister 1 volume fold-change
Sister 2 volume fold-change)
r=0.43, CI (0.36,0.50), p<.001
Control sister 1 volume fold-change
Control sister 2 volume fold-change
r=-0.04, CI (-0.13,0.05), p=.472
Daughter volume fold-change)
Mother volume fold-change
r=-0.07, CI (-0.18,0.05), p=.348
Control daughter volume fold-change
Control mother volume fold-change
r=0.56, CI (0.47,0.66), p<.001
Sister 1 fitted time scaling factor (α)
Sister 2 fitted time scaling factor (α)
r=0.61, CI (0.51,0.69), p<.001
Control sister fitted time scaling factor (α)
Control sister 2 fitted time scaling factor (α)
r=0.11, CI (0.02,0.18), p=0.046
Daughter fitted time scaling factor (α)
Mother fitted time scaling factor (α)
r=-0.03, CI (-0.14,0.08), p=0.664
Control daughter fitted time scaling factor (α)
Control mother fitted time scaling factor (α)

**Figure S12. Nuclear growth duration is inherited from one generation to the next, while other nuclear growth features are controlled for by their colony context. A.** The relationship between the starting volumes for control pairs of nuclei. The control sister pairs are unrelated nuclei born within 10 minutes and a 60 μm radius. These thresholds were selected based on the relative timing and locations of sister pairs growth (**B** left and middle). The unity line is shown for reference. **B.** The distribution of differences between sister nuclei in the locations (left), times (middle) and volumes (right) at their respective starts of growth. These distributions inform the thresholds used to create the control of unrelated pairs (Methods). **C.** The relationship between the nuclear starting volumes, ending volumes, added volumes, growth rates, volume fold-changes, and fitted time scaling factors (top to bottom) for all sister pairs, control sister pairs, mother-daughter pairs, and control mother daughter pairs (left to right) in the lineage-annotated analysis dataset. These are the plots used to create Figure 7G). The unity line (dashed line) is shown for reference and confidence intervals (CI) are the bootstrapped 90% CI (Methods). The control sister pairs are unrelated nuclei born within 10 minutes, a 60 μm radius, and a difference in starting volume less than 80 $\mu m^3$. The control mother-daughter pairs are unrelated nuclei where the control mother's breakdown is within 60 minutes and a 60 μm radius of the control daughter's formation. Their size is constrained such that the control daughter's starting volume is within 60 $\mu m^3$ of half the control mother's ending volume (Methods).

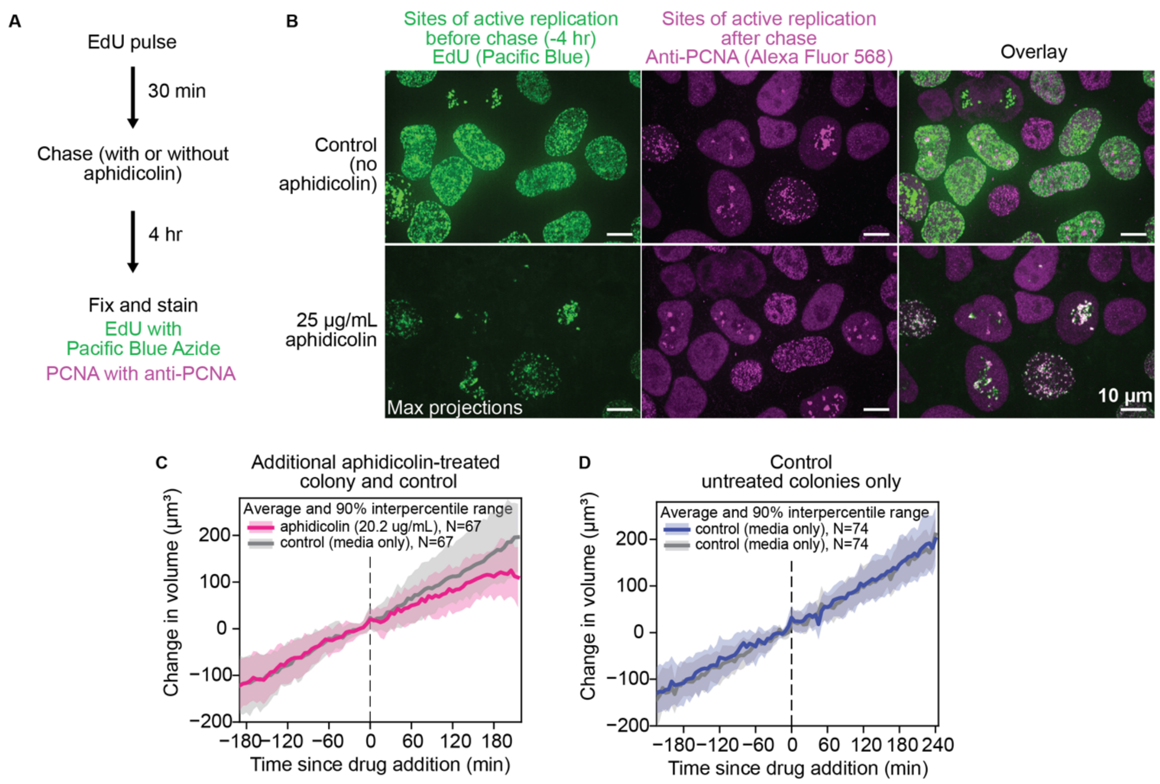


**Figure S13. Treatment with aphidicolin rapidly inhibits progression of DNA replication in hiPS cells. A.** Schematic illustrating this experiment. Cells were given a brief 30 minute pulse with 10 µM EdU, after which they were washed and chased with media only (control) or 25 µg/mL of aphidicolin for 4 hrs and then the cells were fixed and stained. Incorporated EdU was labeled with Pacific Blue Azide, and PCNA was labeled with an antibody against PCNA (for details, see Methods). **B.** Images of cells in the control condition (no aphidicolin, top panel) and aphidicolin treated condition (bottom panel). PCNA localization (magenta) indicates sites of active DNA replication just before fixation. The EdU (via Pacific Blue, green) indicates sites of activate replication 4 hours before fixation. When no inhibitor was added (top panel) sites of active DNA replication were no longer in the same location as 4 hrs earlier (i.e. no overlap between EdU and PCNA). In contrast, the cells treated with aphidicolin show a strong overlap in localization of EdU and PCNA, indicating that aphidicolin treatment caused the DNA replication machinery to stall and remain stalled for the full 4 hrs. Therefore, we are confident that the very minimal change in nuclear growth caused by aphidicolin (seen in Supplemental Fig. S7C and in panels C and D) were not due to low levels of continuing replication. The images shown are maximum intensity projections of 3D fluorescence images acquired with a 100x/1.25 NA objective. **C.** The average change in nuclear volume for nuclei from an additional colony that received a dose of 20.2 µg/mL aphidicolin (magenta) and control colony receiving only media (gray). The change in volume was determined by subtracting the nuclear volume at the time of inhibitor addition (defined here as t=0) from the entire nuclear volume trajectory for all nuclei. Shaded regions represent the 90% interpercentile range. The single nucleus volume trajectories were selected such that each trajectory from the aphidicolin-treated trajectory was matched with a trajectory from the control colony that had the same volume at the frame before inhibitor addition, thus the sample sizes (N) are identical. This was done to account for potential size dependent effects of the inhibitor (see Methods). **D.** The same analysis as in panel C here performed on two untreated control colonies.

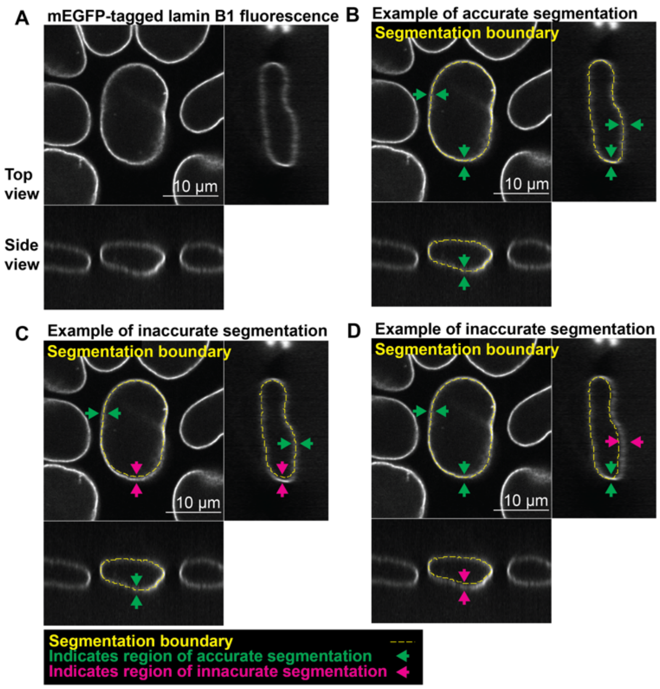


**Figure S14. Illustration of visual evaluation used to curate accuracy of ground truth segmentations of lamin B1 from 100x/1.25 NA images. A.** A 3D image of mEGFP-tagged lamin B1 fluorescence acquired with a 100x/1.25 NA objective is shown in top view and side views (middle slices). The same image is shown in all four panels. See Methods for details. **B.** Example of accurate segmentation. The segmentation is considered accurate if the boundary of segmentation (drawn over the image as a yellow dashed line) lies directly in middle of lamin shell at all points in all views. **C and D**. Examples of inaccurate segmentations that would not be included in the ground truth segmentation training set. This example segmentation in C. has inaccuracies seen in the top view (XY) where the segmentation boundary clearly deviates from the midline of the lamin B1 shell, as indicated by the magenta arrows (compare to green arrows in B). In the example in D., the segmentation has inaccuracies seen in the side views (ZX and ZY).

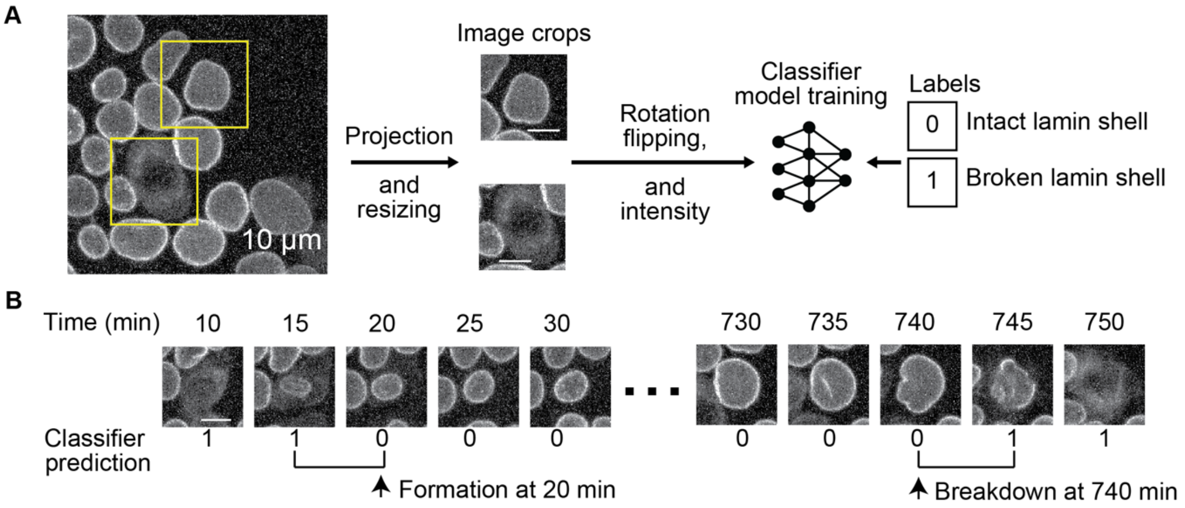


**Figure S15. Automated image analysis workflow for identifying interphase in trajectories of individual nuclei. A.** An illustration of training a deep-learning image classifier to detect nuclei in interphase (nuclei with intact lamin shells) or in mitosis (nuclei with broken lamin shell). To train the convolutional classifier, 64x64 crops were taken from maximum intensity projections along Z (MIPs) of 3D fluorescence images of mEGFP-tagged lamin B1 ("image crops"). Labels were then given to each of these crops to indicate whether the in-frame nucleus had an intact lamin shell (0) or a broken lamin shell (1). The model was then trained with ~65,000 of these image crop and label pairs from 505 single nucleus trajectories. **B.** After training, this image-classifier was applied to all timepoints of all tracked nuclei. Each timepoint is assigned a 0 if the lamin shell is intact, and 1 otherwise. "Formation" and "breakdown" are identified as the first and last timepoints where the nucleus was classified as having an intact lamin shell, respectively. For the example nuclear trajectory shown here, formation occurs at the 20-minute timepoint and breakdown occurs at the 740-minute timepoint, with interphase timepoints in between.

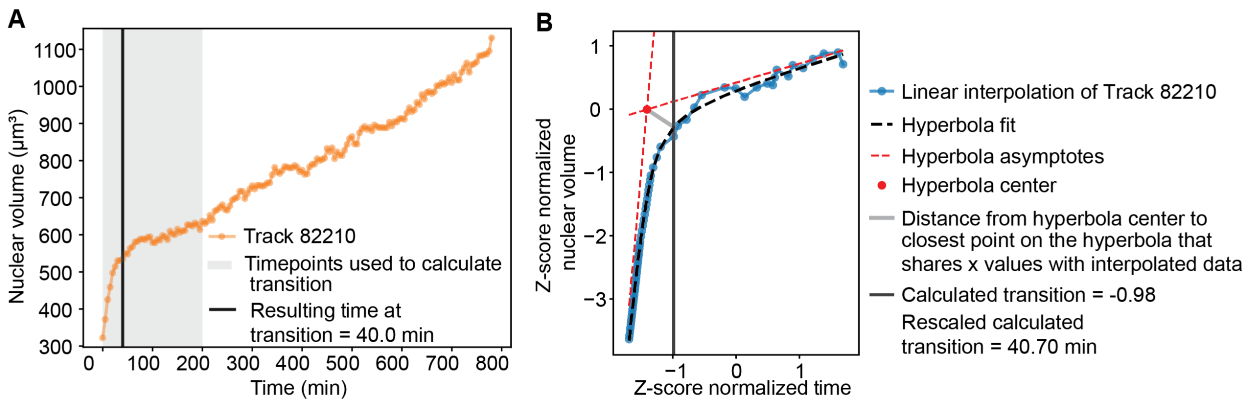


**Figure S16. Calculation of the "transition point" between "expansion" and "growth" in a nuclear volume trajectory. A.** The transition point is calculated by first truncating a nuclear volume trajectory to the first 40 timepoints (grey window). **B.** Illustration of the workflow used to determine transition point. Linear interpolation of the trajectory (blue) increases the number of datapoints in the early part of the trajectory. The volume and time axis were z-score normalized to simplify the hyperbola fit (dashed black) parameters. The hyperbola center (red point) is where the hyperbola asymptotes (dashed red lines) cross. The point on the hyperbola (that shares time values with the interpolated data) that is closest to the hyperbola center (distance shown in grey), is defined as the transition point. After rescaling, this value may not fall on the time axis of the acquired data (40.70 minutes for Track 82210). The nearest acquired frame, within 2 frames (10 minutes), is chosen (40.0 minutes shown in A).

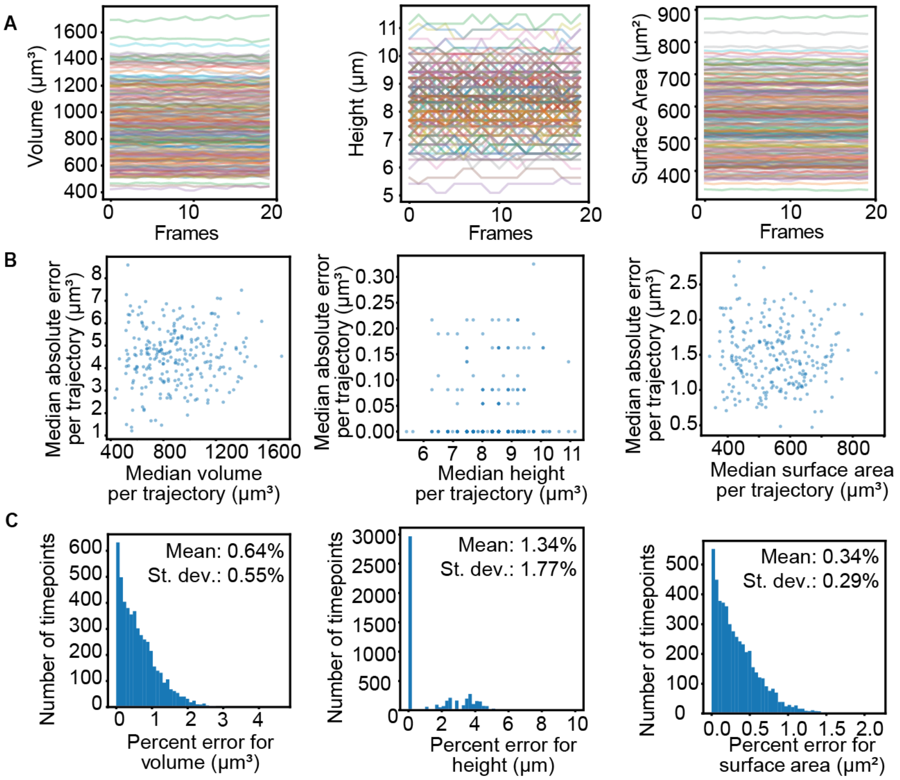


**Figure S17. Evaluation of segmentation model consistency using a fixed nucleus timelapse control. A.** The volume (left), height (middle) and surface area (right) for all nuclei tracked over 20 frames of the fixed nucleus timelapse control (see Methods). Because these nuclei are fixed, variation in the volume, height and surface area are due only to variation in model prediction of features. **B.** The relationship between the median absolute error and median feature value from each trajectory, shown for volume (left), height (middle), and surface area (right). To calculate the percent and absolute error in the measurements we assumed the median value for a given nuclear trajectory was representative of its true value and compared each of the 20 fixed timepoint measurements to the median. These results show that the magnitude of error was not strongly correlated with the magnitude of the feature (e.g., the median absolute errors for larger nuclei were similar to those of smaller nuclei). **C.** The distribution of percent error for volume (left), height (middle), and surface area (right) for each nuclear measurement at every timepoint with the mean and standard deviation. The mean percent error for volume was 0.64% (5.54 $\mu m^3$), and the 95th percentile was 1.71% (14.95 $\mu m^3$). The mean percent error for height was 1.34% (0.11 µm), and the 95th percentile was 4.34% (0.33 µm) which is on the order of the 100x/1.25 NA acquisition z-step (0.29 µm) for ground truth images used to train the Vision Transformer-based deep-learning segmentation model (see Methods). The mean percent error for surface area was 0.34% (1.83 $\mu m^2$) and the 95$^{th}$ percentile was 0.9% (4.89 $\mu m^2$). These results show that the segmentation model predictions were very consistent for the same nucleus over time.

**Supplemental Movies Captions:**

**Movie S1. Paired bright-field and fluorescence timelapses of the Medium colony.** Timelapses associated with Fig. 1A and B. A center slice of the 20x/0.8 NA transmitted light bright-field (left) and a maximum intensity projection of the lamin B1-mEGFP fluorescence (right) of the z-stack images for the Medium colony (see Methods for details). Images were acquired every 5 minutes and movies were generated with a frame rate of 10 frames per second (fps). The blurry, dark spots in the bright-field image are due to light refracted dust on the lid of the 96-well plate.

**Movie S2. Paired bright-field and fluorescence timelapses of the Small colony.** A center slice of the 20x/0.8 NA transmitted light bright-field (left) and a maximum intensity projection of the lamin B1-mEGFP fluorescence (right) of the z-stack images for the Small colony (see Methods for details). Images were generated as described in Movie S1.

**Movie S3. Paired bright-field and fluorescence timelapses of the Large colony.** A center slice of the 20x/0.8 NA transmitted light bright-field (left) and a maximum intensity projection of the lamin B1-mEGFP fluorescence (right) of the Z-stack images for the Large colony (see Methods for details). Images were generated as described in Movie S1.

**Movie S4. Segmentation timelapse of the Medium colony.** Timelapse associated with Fig. 1C showing maximum projections of 3D segmentations of nuclei in the Medium colony (see Methods for details). Nuclear segmentations are colored based on their Track IDs to facilitate visual tracking of each nucleus over time. Segmentations were generated for images acquired every 5 minutes and movies were generated with a frame rate of 10fps.

**Movie S5. Segmentation timelapse of the Small colony.** Segmentation timelapse of the Small colony, with details as in Movie S4.

**Movie S6. Segmentation timelapse of the Large colony.** Segmentation timelapse of the Large colony, with details as in Movie S4.

**Movie S7. Nuclear height varies on the timescale of days.** Timelapse corresponding to the images in Fig. 2A and Fig. 3B, showing maximum projections of 3D nuclear segmentations in the Medium colony colorized by nuclear height. They were created in the Timelapse Feature Explorer (Methods) and exported at a frame rate of 10 fps. Due to the nature of viewing 3D segmentations as 2D maximum projections, the apparent nuclear size and shape in these images can be misleading; the color-mapping of these images is the best way to interpret their size and shape. The colormap limits are chosen to highlight the variation of nuclear features across the population for nuclei present in the baseline colonies dataset (all others are shown in gray, Methods). The colormap limits are 4 μm (cyan) and 11 μm (magenta).

**Movie S8. Nuclear volume varies on the timescale of hours.** Timelapse corresponding to the images in Fig. 2B, showing maximum projections of 3D nuclear segmentations in the Medium colony color-mapped to nuclear volume. Details of movie generation as in Movie S7. The colormap limits are 400 $\mu m^3$ (cyan) and 1200 $\mu m^3$ (magenta).

**Movie S9. Nuclear aspect ratio varies on the timescale of minutes.** Timelapse corresponding to the images in Fig. 2C, showing maximum projections of 3D nuclear segmentations in the Medium colony color-mapped to nuclear aspect ratio (the length of the longest axis divided by the orthogonal in-plane axis). Details of movie generation as in Movie S7. The colormap limits are 1.5 (cyan) and 11 μm (magenta).

**Movie S10. Nuclear height varies on the timescale of days (Small colony).** Timelapse corresponding to the images in Fig. 3A, showing maximum projections of 3D nuclear segmentations in the Small colony color-mapped to nuclear height. Details of movie generation as in Movie S7. The colormap limits are 4 μm (cyan) and 11 μm (magenta).

**Movie S11. Nuclear height varies on the timescale of days (Large colony).** Timelapse corresponding to the images in Fig. 3C, showing maximum projections of 3D nuclear segmentations in the Large colony color-mapped to nuclear height. Details of movie generation as in Movie S7. The colormap limits are 4 μm (cyan) and 11 μm (magenta).
